# A unified framework for prioritizing habitat and connectivity conservation through analytical sensitivity

**DOI:** 10.64898/2026.01.05.697654

**Authors:** Bram Van Moorter, Ilkka Kivimäki, Manuela Panzacchi, Rafael Schouten, Bert Wuyts, Bernardo Brandão Niebuhr, Marco Saerens, Victor Boussange

## Abstract

**Context:** Connected habitats underpin key ecological processes such as movement, gene flow, and recolonization. As land-use change accelerates habitat loss and fragmentation, there is an urgent need to identify priority areas for conserving habitat and maintaining connectivity.

**Objectives:** We develop a unified spatial prioritization framework for functional connectivity that is grounded in metapopulation theory, accommodates species-specific movement behavior, and captures the dual role of landscape units as both habitat providers and connectivity facilitators.

**Methods:** We formalize the landscape as a network in which nodes represent spatial units characterized by habitat quality and edges encode permeability for movement. Pairwise ecological proximities are defined from species-specific movement capacity and an ecological distance that integrates costs across paths, expressed as least-cost distance, effective resistance, expected cost, or survival probability depending on the movement ecology of the focal species. From these proximities and node qualities, we construct the landscape matrix. Based on the sensitivity of the landscape matrix to local perturbations in habitat quality or permeability, we develop a prioritization approach that identifies where local changes most strongly influence landscape-scale habitat connectivity.

**Results:** Our framework unifies widely used landscape connectivity metrics across the full spectrum of movement models, including least-cost paths, circuit theory, spatial absorbing Markov chains, and randomized shortest paths. We show that summation-based metrics serve as computationally efficient approximations of metapopulation capacity, enabling evaluation of functional connectivity in large, high-resolution landscapes. We derive sensitivities for these metrics with respect to habitat quality and permeability and show that they correspond to two families of centrality measures: a closeness-like centrality reflecting the influence of habitat quality, and a betweenness-like centrality reflecting the influence of habitat permeability. Applying our prioritization approach based on these sensitivities to a case study involving wild reindeer in Norway, we identify high-value habitat and key movement corridors that closely align with a reference node-removal approach while reducing computation time by several orders of magnitude.

**Conclusions:** By linking metapopulation theory with sensitivity analysis, our work provides a unified and scalable framework for spatial prioritization of functional connectivity. It supports evidence-based management of fragmented landscapes and advances both theoretical understanding and practical application in landscape ecology.

## 1. Introduction

As human land-use transforms much of the Earth’s surface (Díaz et al., 2019; Barnosky et al., 2012), habitat loss and fragmentation have become major drivers of biodiversity decline, contributing to the so-called ‘sixth mass extinction’ (Foley et al., 2005; Dirzo et al., 2014). Halting this trend and promoting sustainable development require targeted conservation strategies that identify where interventions can most effectively preserve biodiversity. Spatial conservation prioritization, defined as the systematic identification of ecologically significant areas and the optimization of limited resources (Moilanen et al., 2009), provides a key framework for this task. However, understanding where and how local changes can propagate to affect whole landscapes demands concepts and methods that link the properties of fine-scale spatial units to emergent landscape-level metrics. Here, landscape connectivity plays a central role: it supports ecological processes and facilitates species movement across fragmented habitats, serving as a bridge between local and landscape scales (Haddad et al., 2015; Beger et al., 2022).

To quantify landscape connectivity, ecologists have developed increasingly complex connectivity models, most of which are based on a network representation of the landscape. Initial approaches in metapopulation theory represented landscapes as graphs in which edges between vertices were defined solely by their Euclidean distance (Hanski, 1999). However, Euclidean distance assumes that movement between two locations depends only on their spatial proximity and ignores the heterogeneity of the intervening landscape. Hence, subsequent models incorporated spatial heterogeneity in landscape permeability: least-cost path (LCP) models compute the path with the lowest cumulative cost of traversing the landscape, assuming organisms follow globally optimal routes (Adriaensen et al., 2003). In contrast, models based on circuit theory (CT) treat movement as electrical currents, capturing diffusive, random-walk behavior without assuming knowledge of the destination (McRae, 2006). More recently, the randomized shortest paths (RSP) framework has unified these paradigms, providing a generalized movement model that can be tuned to species-specific cognition and decision rules (Saerens et al., 2009; Van Moorter et al., 2021). These movement models underpin a range of *landscape functionality* metrics that quantify the total amount of functionally connected habitat for a landscape (see Table 1 for definition), such as metapopulation capacity (Hanski, 1999) or equivalent connected habitat (Saura et al., 2011; Van Moorter et al., 2023b).

**Table 1:**
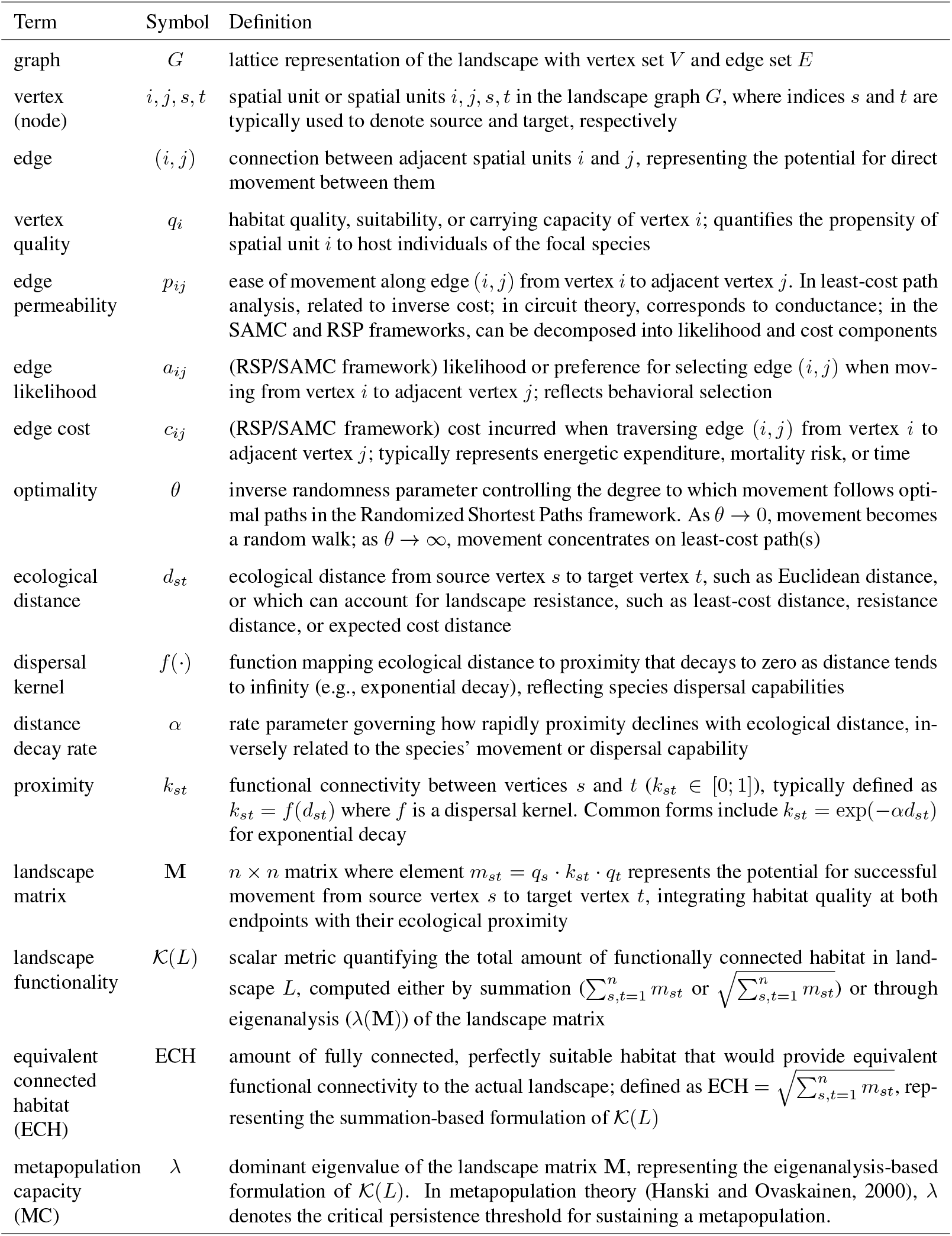
Notation and definitions of key quantities. The landscape is formalized as a graph *G* with vertices representing spatial units and edges representing adjacent neighbors. For further details see main text and Appendix C.

| Term | Symbol | Definition |
| --- | --- | --- |
| graph | $G$ | lattice representation of the landscape with vertex set $V$ and edge set $E$ |
| vertex (node) | $i, j, s, t$ | spatial unit or spatial units $i, j, s, t$ in the landscape graph $G$, where indices $s$ and $t$ are typically used to denote source and target, respectively |
| edge | $(i, j)$ | connection between adjacent spatial units $i$ and $j$, representing the potential for direct movement between them |
| vertex quality | $q_i$ | habitat quality, suitability, or carrying capacity of vertex $i$; quantifies the propensity of spatial unit $i$ to host individuals of the focal species |
| edge permeability | $p_{ij}$ | ease of movement along edge $(i, j)$ from vertex $i$ to adjacent vertex $j$. In least-cost path analysis, related to inverse cost; in circuit theory, corresponds to conductance; in the SAMC and RSP frameworks, can be decomposed into likelihood and cost components |
| edge likelihood | $a_{ij}$ | (RSP/SAMC framework) likelihood or preference for selecting edge $(i, j)$ when moving from vertex $i$ to adjacent vertex $j$; reflects behavioral selection |
| edge cost | $c_{ij}$ | (RSP/SAMC framework) cost incurred when traversing edge $(i, j)$ from vertex $i$ to adjacent vertex $j$; typically represents energetic expenditure, mortality risk, or time |
| optimality | $\theta$ | inverse randomness parameter controlling the degree to which movement follows optimal paths in the Randomized Shortest Paths framework. As $\theta \rightarrow 0$, movement becomes a random walk; as $\theta \rightarrow \infty$, movement concentrates on least-cost path(s) |
| ecological distance | $d_{st}$ | ecological distance from source vertex $s$ to target vertex $t$, such as Euclidean distance, or which can account for landscape resistance, such as least-cost distance, resistance distance, or expected cost distance |
| dispersal kernel | $f(\cdot)$ | function mapping ecological distance to proximity that decays to zero as distance tends to infinity (e.g., exponential decay), reflecting species dispersal capabilities |
| distance decay rate | $\alpha$ | rate parameter governing how rapidly proximity declines with ecological distance, inversely related to the species' movement or dispersal capability |
| proximity | $k_{st}$ | functional connectivity between vertices $s$ and $t$ ($k_{st} \in [0; 1]$), typically defined as $k_{st} = f(d_{st})$ where $f$ is a dispersal kernel. Common forms include $k_{st} = \exp(-\alpha d_{st})$ for exponential decay |
| landscape matrix | $\mathbf{M}$ | $n \times n$ matrix where element $m_{st} = q_s \cdot k_{st} \cdot q_t$ represents the potential for successful movement from source vertex $s$ to target vertex $t$, integrating habitat quality at both endpoints with their ecological proximity |
| landscape functionality | $\mathcal{K}(L)$ | scalar metric quantifying the total amount of functionally connected habitat in landscape $L$, computed either by summation ($\sum_{s,t=1}^n m_{st}$ or $\sqrt{\sum_{s,t=1}^n m_{st}}$) or through eigenanalysis ($\lambda(\mathbf{M})$) of the landscape matrix |
| equivalent connected habitat (ECH) | ECH | amount of fully connected, perfectly suitable habitat that would provide equivalent functional connectivity to the actual landscape; defined as $\text{ECH} = \sqrt{\sum_{s,t=1}^n m_{st}}$, representing the summation-based formulation of $\mathcal{K}(L)$ |
| metapopulation capacity (MC) | $\lambda$ | dominant eigenvalue of the landscape matrix $\mathbf{M}$, representing the eigenanalysis-based formulation of $\mathcal{K}(L)$. In metapopulation theory (Hanski and Ovaskainen, 2000), $\lambda$ denotes the critical persistence threshold for sustaining a metapopulation. |

Identifying priority areas for conservation requires understanding how individual spatial units contribute to overall landscape functionality. Spatial units play a dual role in landscape networks: (1) they provide functional habitat and (2) they act as connectors that facilitate movement between habitat areas (Saura and Rubio, 2010). Losing these functions leads to habitat loss and fragmentation, respectively – the primary drivers of biodiversity decline (Fahrig, 2003; Haddad et al., 2015). Connectivity is often incorporated into spatial conservation prioritization through measures such as patch contiguity or dispersal kernels (Hanson et al., 2022), but these rely on Euclidean distances and therefore ignore the connector role of spatial units. This connector role is made explicit in betweenness-like or medial centrality metrics (sensu Borgatti and Everett, 2006), which quantify the number of paths that pass through a spatial unit. Least-cost betweenness (Bodin and Saura, 2010) and current flow derived from circuit theory (Hodgson et al., 2016) have been proposed to identify areas important for maintaining landscape connectivity and can be included as features in spatial conservation prioritization (Daigle et al., 2020). However, these approaches focus on the connector function of a spatial unit and do not explicitly represent its contribution as habitat. Prioritization based on the effect of iterative node-removal analysis on landscape functionality is conceptually straightforward and captures both habitat and connector roles (Saura and Torne, 2009). Yet this approach becomes computationally prohibitive for large or high-resolution networks (Van Moorter et al., 2023b). Thus, existing connectivity metrics for conservation prioritization typically emphasize only one of the dual roles played by spatial units in the landscape network, while more comprehensive approaches remain computationally infeasible at the spatial resolutions needed for land planning.

In this study, we develop a unified framework for spatial prioritization based on sensitivity analysis (Kivimäki et al., 2024). We formalize the landscape as a weighted graph in which vertices represent spatial units with associated habitat quality and edges encode habitat permeability. Ecological proximity between two areas is defined using species-specific movement capacity and an ecological distance that integrates permeability across paths, which can take the form of least-cost distance, resistance distance, expected-cost distance, or survival probability. From these proximities and vertex qualities, we construct a landscape matrix that can be summarized by summation or eigenanalysis (spectral decomposition) to quantify landscape functionality. This formulation links to metapopulation capacity, yields sensitivity measures with clear interpretations in terms of weighted-network centralities, and enables scalable prioritization of both habitat quality and permeability. We illustrate the framework with a case study of wild reindeer in Norway, identifying areas important for habitat and connectivity protection and comparing the outcomes of a sensitivity-based spatial conservation prioritization with iterative node-removal analysis.

## 2. Methods

We present a quantitative framework for assessing the contribution of individual spatial units to landscape-level connectivity via sensitivity analysis. First, we formalize the landscape as a weighted graph and define landscape functionality as a summary statistic of the landscape matrix, which is constructed from pairwise ecological proximities between vertices and their habitat quality (see Table 1). These proximities reflect the permeability of the landscape to movement by accommodating different movement models spanning least-cost path, circuit theory, absorbing random walks, and randomized shortest paths. We then characterise the marginal contribution of habitat quality and landscape permeability of a single spatial unit to the landscape functionality, deriving analytical expressions for the partial derivatives of the summary statistic with respect to quality and permeability. On this basis, we propose a prioritization heuristic that uses these quantities to identify priority areas for conservation. Finally, we introduce an empirical case study from a wild reindeer population in Norway to evaluate the framework’s computational efficiency and ecological relevance.

### 2.1. Formal representation of the landscape

#### Landscape as a weighted graph

We formalize the landscape as a weighted graph *G* = (*V, E*), where the vertex set *V* represents discrete spatial units and the edge set *E* encodes movement potential between adjacent units. Each vertex *i* ∈ *V* corresponds to a spatial unit (i.e., a habitat patch or a grid cell) and is characterized by a non-negative quality value *q*_*i*_ ≥ 0 that quantifies the unit’s propensity to support individuals of the focal species. This vertex attribute may represent habitat suitability derived from species distribution models, patch area, carrying capacity, or observed population size, depending on the ecological context and data availability. Similarly, we parameterize edge (*i, j*) through its permeability *p*_*ij*_, which quantifies the ease of movement along that edge (Fig. 1b). Permeability serves as a unifying concept across movement frameworks: in circuit theory, permeability corresponds to conductance (the inverse of resistance); in least-cost path analysis, it relates to the inverse of movement cost. The randomized shortest paths and absorbing random walks frameworks introduce a richer parameterization of permeability by distinguishing between the likelihood and cost of movement (see section below).

**Figure 1.**
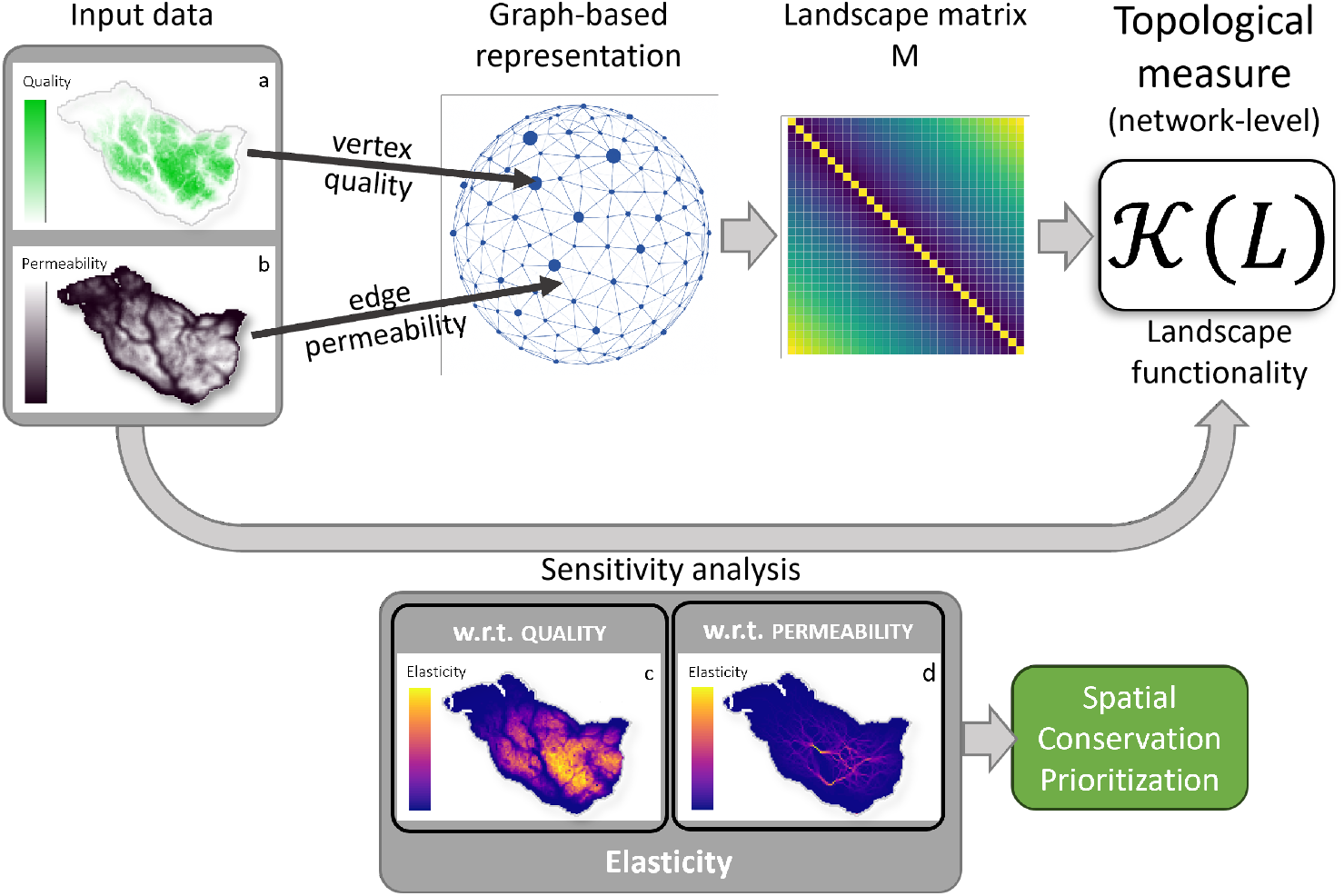
Sensitivity analysis of the landscape network. The landscape graph is parameterized through the habitat quality (*q*_*i*_ in panel a) and permeability (*c*_*j*_ = −*log*(*a*_*j*_) in panel b) inputs, from which the landscape matrix M is derived. The landscape matrix can be summarized through either summation or eigenanalysis into a topological (i.e., network-level) metric of the amount and connectedness of the habitat in the landscape. The sensitivity analysis computes the response of this topological measure – the landscape functionality 𝒦 (*L*) – to a local perturbation of the network (i.e. *q, a*, or *c*). We computed the sensitivity to the proportional change – i.e., the elasticity to vertex quality (c) and edge permeability (d) – to inform spatial conservation prioritization. See main text for detailed explanation.

#### Ecological proximity between spatial units

From the local edge permeabilities *p*_*ij*_, we derive a pairwise proximity metric *k*_*st*_ between any two vertices *s, t ∈ V*, not necessarily adjacent, that quantifies the functional connectivity between them (Fig. 1c). *k*_*st*_ integrates information across the set of possible paths connecting *s* to *t*, weighted according to an assumed movement model, the permeabilities of edges along those paths, and the limited capacity of the focal species to travel long distances. Specifically, we define proximity as a decreasing function of an ecological distance *d*_*st*_:

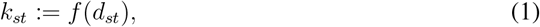

where *d*_*st*_ is a quasimetric on the graph that reflects the effective distance from source *s* to target *t*, and *f* : ℝ_≥0_ → [0, 1] is a dispersal kernel function chosen to match the species’ movement or dispersal capability, with *f* (*d*) → 0 as *d* → ∞. A common choice is the exponential kernel (Hanski, 1999), *k*_*st*_ = exp(−*αd*_*st*_), where the decay rate *α* ≠ 0 is inversely related to dispersal ability: species with greater mobility have smaller *α*, yielding higher connectivity over long distances. Under certain formulations, *k*_*st*_ can be directly interpreted as the probability of successful movement or gene flow between *s* and *t* (Hanski, 1999; *Bullock et al., 2017)*.

The choice of ecological distance *d*_*st*_ encodes the assumed movement behavior of the focal species (reviewed in: Van Moorter et al., 2021). When movement is unconstrained by landscape structure – typical of aerial dispersers – *d*_*st*_ is simply the Euclidean distance. For species with perfect knowledge and optimal movement across the landscape, the least-cost path (LCP) distance is appropriate; here, *d*_*st*_ minimizes the cumulative cost along a single optimal trajectory. For species exhibiting diffusive or exploratory behavior with no knowledge of the destination, the resistance distance of CT can be used for *d*_*st*_, corresponding to the commute time of a random walk on the graph between vertices *s* and *t*. The Randomized Shortest Paths framework (RSP; Saerens et al., 2009; Van Moorter et al., 2021) unifies these approaches by introducing a parameter *θ* that continuously interpolates between random-walk behavior of CT (*θ* → 0) and the deterministic optimality of LCP (*θ* → ∞). Finally, while standard models typically conflate movement likelihood and cost into a single permeability value, the absorbing random walks model (i.e., spatial absorbing Markov chain – SAMC; Fletcher Jr et al., 2019) incorporates mortality by separating the likelihood and cost of movement. Because RSP is based on absobring random walks, it similarly distinguishes between the *likelihood* of selecting an edge, *a*_*ij*_ (reflecting behavioral preference or structural guidance), and the *cost* of traversing it, *c*_*ij*_ (reflecting energy expenditure or mortality risk). This separation allows modelers to capture more complex behaviors in which, for instance, an organism might prefer a path (high *a*_*ij*_) that is nonetheless costly (high *c*_*ij*_), or avoid a low-cost path due to behavioral avoidance. The two can be coupled via, e.g., *c*_*ij*_ = − log(*a*_*ij*_), recovering standard formulations, but their decoupling may provide flexibility needed for realistic ecological modeling.

#### Landscape matrix

From the vertex qualities *q*_*i*_ and pairwise proximities *k*_*st*_, we construct the landscape matrix M (Hanski, 1999; Ovaskainen, 2003), an *n* × *n* matrix where *n* = |*V* | is the number of spatial units. Each element of M is defined as

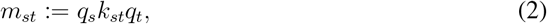

representing the potential for successful movement-mediated interaction between source *s* and target *t* (Fig. 1d). This formulation integrates three components: the quality of the source (reflecting emigration potential or propagule production), the proximity between source and target (reflecting landscape-mediated connectivity), and the quality of the target (reflecting immigration potential or establishment probability). The landscape matrix **M** thus encapsulates how habitat amount, habitat configuration, and landscape permeability jointly determine effective connectivity (Van Moorter et al., 2021).

#### Landscape functionality

We define landscape functionality, 𝒦 (*L*), of landscape *L* as a scalar summary metric of the landscape matrix **M** that quantifies the total amount of functionally connected habitat (see also Appendix A). We consider two canonical formulations of 𝒦 (*L*), corresponding to the two primary perspectives in metapopulation theory (see Appendix B): a summation-based metric,

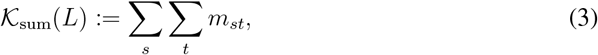

and a spectral formulation based on the dominant eigenvalue,

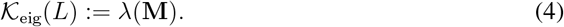

Because 𝒦(*L*) provides a global summary of landscape function, it serves as a natural quantity for conservation. By maximizing 𝒦(*L*), we maximize either the total connected habitat or the capacity for long-term persistence.

### 2.2. Sensitivity analysis of landscape functionality

To characterise the contribution of a spatial unit to landscape functionality, we quantify the effect of a small perturbation in quality or permeability on 𝒦(*L*). Formally, this corresponds to the sensitivity – equivalently, the partial derivative – of 𝒦(*L*) with respect to vertex quality *q*_*i*_ or to the edge-level permeability parameters. The sensitivity to vertex quality, 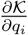, measures the marginal contribution of the habitat in spatial unit *i* to overall landscape functionality. Units with high sensitivity contribute disproportionately to landscape functionality. The sensitivities to permeability, 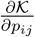, measure how strongly landscape functionality depends on the capacity to move along vertex *i* − *j*. High sensitivities indicate edges (or the spatial units they traverse) that disproportionately sustain connectivity supporting landscape functionality. We derived closed-form expressions for these sensitivities across the full spectrum of movement models (LCP, CT, SAMC, RSP) and for both connectivity formulations (summation and spectral). These analytical results (detailed in Appendix C) allow us to compute sensitivities exactly and to link them directly to well-known centrality metrics in network analysis.

#### Numerical implementation

We implemented the sensitivity analysis within the `ConScape` library (see Appendix D for an example and tutorial of the workflow).

For a landscape with *n* spatial units, constructing the landscape matrix **M** requires computing *n* × *n* = *n*^2^ pairwise proximities (**K**), which becomes demanding both in computation time and working memory for high-resolution, large-extent landscapes. To address this, we implemented landmark-based algorithms (Van Moorter et al., 2023a), where distances or proximities are computed from *n* vertices to *m* landmarks (see the *n*-to-*m* method, pp. 111 in Kivimäki, 2018). Depending on the number of landmarks (*m*), this approach can substantially reduce computational load with only limited loss of accuracy (Van Moorter et al., 2023a).

Here, we used this landmark approach to (1) evaluate how the accuracy of estimated sensitivities changes with decreasing numbers of landmarks (*m*), and (2) assess the accuracy of sensitivities estimated when all *n* spatial units are distributed across *n/m* parallel processes. These tests illustrate how the analytical method can be scaled to large landscapes while maintaining computational efficiency. Results are presented in the Supplementary Material (Appendix E).

### 2.3. Spatial conservation prioritization

#### Elasticity-based prioritization approach

We propose a conservation prioritization approach that ranks spatial units according to their degradation potential and how strongly changes in habitat quality or permeability affect overall landscape functionality. Raw sensitivities capture only the responsiveness of landscape functionality to unit local perturbations and do not account for the magnitude of the perturbation itself. To obtain a metric suitable for prioritization, we therefore compute the elasticity of 𝒦 (*L*) with respect to each parameter *x* (where *x* is quality *q*_*i*_ or permeability *p*_*ij*_):

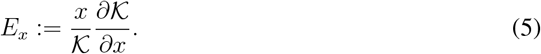

Elasticity *E*_*x*_ measures the proportional change in landscape functionality caused by a proportional change in habitat quality or permeability at a given location. This makes elasticities directly interpretable as indicators of where degradation would have the greatest proportional impact, while also providing a scale-free metric that enables comparison across variables.

##### 1. Habitat prioritization

Rank spatial units by 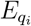. High-ranking units are habitats whose degradation would disproportionately reduce landscape functionality.

##### 2. Connectivity prioritization

Rank spatial units by 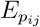. High-ranking units represent connectivity bottlenecks whose reduced permeability would disproportionately diminish movement-mediated landscape functionality.

This elasticity-based prioritization therefore highlights spatial units exerting the strongest leverage on landscape functionality under small proportional changes.

#### Validation against node-removal analysis

Elasticities reflect the response of landscape functionality to infinitesimally small perturbations and therefore do not represent the complete loss of a spatial unit. To assess how well elasticities approximate larger changes in empirical landscapes, we validate the approach against a node-removal analysis, in which landscape functionality is recomputed after removing each spatial unit; we refer to the resulting finite loss in functionality as the node’s criticality. Despite its computational challenges for large or high-resolution networks, we treat node-removal as the reference against which to benchmark our analytical approach for landscape conservation (Saura and Torne, 2009; Saura and Rubio, 2010).

We considered two node-removal scenarios to quantify the criticality of a node: (1) removal of habitat quality only, where the node remains in the network and continues to act as a connector, and (2) complete removal of the node, which eliminates both its habitat contribution and its role in facilitating movement. The first scenario represents situations where habitat suitability is lost while connectivity remains – such as when forest is replaced by a non-suitable land cover that still allows movement between adjacent habitat areas. Because a node’s connector role cannot be removed independently of its habitat, we isolated the additional effect of lost connectivity as the difference between scenarios (2) and (1).

These comparisons allow us to evaluate how closely elasticity-based prioritization corresponds to rankings obtained from node-removal analysis and to quantify the computational advantages of the analytical approach in a realistic landscape prioritization context.

### 2.4. Case study: wild reindeer in Norway

We conducted the benchmark using a spatial prioritization case study in the Snøhetta wild reindeer area in Norway. Although Norwegian wild reindeer have not experienced the severe population declines observed elsewhere (Vors and Boyce, 2009), their habitat has become smaller and increasingly fragmented due to anthropogenic land use and infrastructure development (Nellemann et al., 2003). Quantifying how connected habitat is distributed, and how it is affected by human activity, is therefore crucial for effective land management and for ensuring the long-term availability of suitable habitat for reindeer populations in Norway (Rolandsen et al., 2022). In previous work, GPS tracking data from wild reindeer were used in a habitat selection analysis to estimate habitat quality (Panzacchi et al., 2015) and in a step selection analysis to estimate habitat permeability and step likelihood (Panzacchi et al., 2016). We refer readers to these earlier studies for details on data and modelling procedures. Here, we use the predicted habitat quality and permeability from those analyses. Specifically, we linked movement cost to step likelihoods by assuming that the cost of moving into vertex *j* follows a negative logarithm with the likelihood, *c*_*j*_ = −log(*a*_*j*_), where *a*_*j*_ > 10^−12^ for connected nodes. For the present illustrations – particularly to enable iterative node-removal analysis – the habitat and permeability maps were resampled to approximately 5, 000 spatial units of 1 × 1 km resolution (for more details on this dataset, see Van Moorter et al., 2023a).

## 3. Results

### 3.1. Unifying landscape functionality metrics

We show that widely used landscape functionality metrics arise as special cases of a single landscape-matrix formulation and that both summation-based and eigenanalysis-based summaries of this matrix connect naturally to metapopulation persistence (see Appendix A and Appendix B).

The landscape matrix **M** is often summarized through its dominant eigenvalue (Eq. 4), which corresponds to metapopulation capacity (MC) (Hanski and Ovaskainen, 2000). The MC provides a rigorous persistence threshold determined by colonization and extinction rates. However, spectral decomposition of **M** becomes computationally infeasible for large or high-resolution landscapes.

Motivated by these computational constraints, a broad class of landscape functionality metrics instead summarizes **M** by summing its entries (Eq. C.7), which aggregates all pairwise connections and treats the landscape as a pool of functionally connected habitat (Van Moorter et al., 2023b). When *d*_*st*_ is Euclidean distance, 𝒦_sum_(*L*) recovers Hanski’s neighbourhood habitat area index (NHAI; Hanski, 1999); when *d*_*st*_ is least-cost distance, it yields the probability of connectivity (PC; Saura and Pascual-Hortal, 2007). When costs are defined as the logarithm of step probabilities, the exponential proximity equals the product of those probabilities along the least-cost path, and PC can be interpreted as the probability that two randomly located individuals occupy mutually reachable habitat (Saura and Pascual-Hortal, 2007; Rubio and Saura, 2012). Taking the square root of PC gives the equivalent connected area (ECA; Saura et al., 2011), which expresses landscape functionality in area units as the amount of perfectly connected habitat equivalent to the observed landscape. Extending this concept from patch area to continuous habitat quality defines the equivalent connected habitat (ECH; Van Moorter et al., 2023a), which generalizes ECA to heterogeneous, grid-based landscapes. All these metrics are variants of 𝒦_sum_(*L*), which can be computed efficiently using a moving-window procedure that avoid explicit construction of the full landscape matrix, an essential advantage when the number of spatial units is large.

In Appendix B we show that 𝒦_sum_(*L*) arises naturally from a landscape-scale mean-field approximation of a stochastic colonization–extinction model, yielding a persistence condition

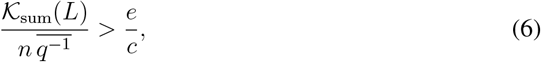

where 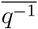 is the mean of the inverse habitat quality across spatial units and *e/c* is the ratio of extinction to colonization rates. This condition mirrors the eigenvalue-based threshold and establishes 𝒦_sum_(*L*) as a computationally tractable, theoretically grounded proxy for metapopulation capacity in large, high-resolution landscapes.

These results clarify how widely used connectivity metrics fit into a unified landscape-matrix framework and justify the use of the summation-based formulation as a scalable proxy for metapopulation capacity.

### 3.2. Sensitivity analysis

The sensitivity of 𝒦_sum_(*L*) with respect to habitat quality and permeability of a given spatial unit quantifies the marginal contribution of that spatial unit to landscape functionality. In Appendix C we derive closed-form expressions of these sensitivities for least-cost path (LCP), circuit theory (CT), spatial absorbing Markov chains (SAMC), and randomized shortest paths (RSP). Here, we highlight the main insights and their ecological interpretation.

First, the sensitivity of landscape functionality to habitat quality generalizes closeness centrality. For any proximity *k*_*st*_, the derivative of 𝒦_sum_(*L*) with respect to the quality of spatial unit *j* is

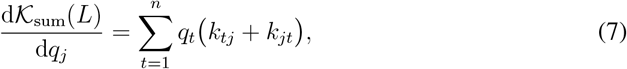

where *k*_*st*_ represents the ecological proximity between spatial units *s* and *t*. This expression shows that a unit’s influence depends jointly on how well it can send individuals to, and receive individuals from, other high-quality units. When qualities are set to one and proximities are defined as the inverse of ecological distance, this reduces to harmonic centrality (Boldi and Vigna, 2014). The quality sensitivity is therefore a habitat-weighted extension of closeness centrality that applies consistently across Euclidean, least-cost, circuit, SAMC, and RSP-based distances. Ecologically, spatial units with high sensitivity to quality are those where even small changes in habitat value disproportionately alter landscape functionality.

Second, the sensitivity of landscape functionality to permeability generalizes betweenness-like measures of movement flow. For summation-based landscape functionality, the derivatives with respect to movement cost (and step likelihood in SAMC and RSP frameworks) at unit *j* can be expressed in terms of the expected number of passages through *j* and related negative covariances between passage frequency and path cost (Table C.3). In the least-cost path limit, this sensitivity is proportional to the number of optimal paths that traverse *j*, weighted by the qualities and proximities of the corresponding source–target pairs, recovering a generalized betweenness metric (Bodin and Saura, 2010). In the circuit-theory limit, the relevant term corresponds to the electrical power (or stress) carried by edges entering *j*, so edges that concentrate current have large influence on landscape functionality – consistent with the use of cumulative current flow for connectivity conservation prioritization (McRae et al., 2008; Hodgson et al., 2016). Along the same lines, in the SAMC framework sensitivity is proportional to the number of paths through an edge, whereas for the RSP framework sensitivities depend on how much more often cost-aware movement uses *j* compared to an unbiased random walk, highlighting locations where global route optimization strongly alters movement patterns. Across these movement models, high permeability sensitivity identifies structural bottlenecks where small changes in permeability cause large changes in landscape functionality.

Finally, we show that the same structural patterns govern sensitivities for both summation- and eigenvalue-based landscape functionality metrics. For eigenvalue-based functionality 𝒦_eig_(*L*) = *λ*(**M**), the sensitivities to habitat quality and permeability involve the same betweenness- and covariance-like terms as above, but weighted by the left and right eigenvectors of **M** (Table C.3 and Appendix C). This parallels classical results on the sensitivity of population projection matrices (Caswell, 2019; Ovaskainen and Hanski, 2003) and links directly to the metapopulation capacity interpretation introduced in Appendix B. As a result, ranking spatial units by their influence on summation-based and eigenvalue-based connectivity produces closely aligned prioritizations. Sensitivity maps based on 𝒦_sum_(*L*) can therefore be interpreted as computationally efficient surrogates for the more computationally demanding eigenvalue sensitivities, while retaining a direct connection to persistence thresholds.

These results show that our analytical sensitivities provide a principled bridge between network centrality, movement ecology, and metapopulation theory. They reveal which spatial units act as key habitat providers or connectivity facilitators under realistic movement models, in a manner that is interpretable via familiar centrality measures and scalable to large landscapes.

### 3.3. Prioritization approach benchmark with application in wild reindeer in Norway

We next benchmarked the use of the derived sensitivities for spatial conservation prioritization, applying the framework introduced in Section 2.3. Specifically, we calculated elasticities with respect to both habitat quality and permeability for the Snøhetta wild reindeer area in Norway, and compared the resulting prioritization maps to outcomes from node-removal analysis.

The elasticity with respect to habitat quality (Fig. 2c and 3a) identifies the central part of the reindeer area as having the strongest influence on landscape functionality. Habitat quality in the Snøhetta region reflect both environmental conditions and human activity. Urban expansion and transport infrastructure reduce habitat suitability around the periphery, while tourist and hydropower developments affect more central areas (Fig. 2a; Panzacchi et al. 2015). The elasticity with respect to permeability (Fig. 2d and 3b) highlights narrow movement corridors linking eastern and western parts of the range. Movement is constrained by rugged topography (Fig. 2b; Panzacchi et al. 2016). Within this context, the narrow land bridge north of the Aursjøen reservoir emerges as a key connector, and the corridor immediately south of it provides an additional link between eastern and western subpopulations (Fig. 3). Conversely, the large, high-quality area east of Aursjøen – located within the Dovrefjell–Sunndalsfjella National Park – contains the highest-quality habitat in the region, whereas both movement bottlenecks lie outside formal protection but are recognized as management priorities (Jordhøy et al., 2012). These maps translate the analytical sensitivities into spatially explicit indicators of where proportional degradation in habitat or permeability would most reduce landscape functionality.

**Figure 2.**
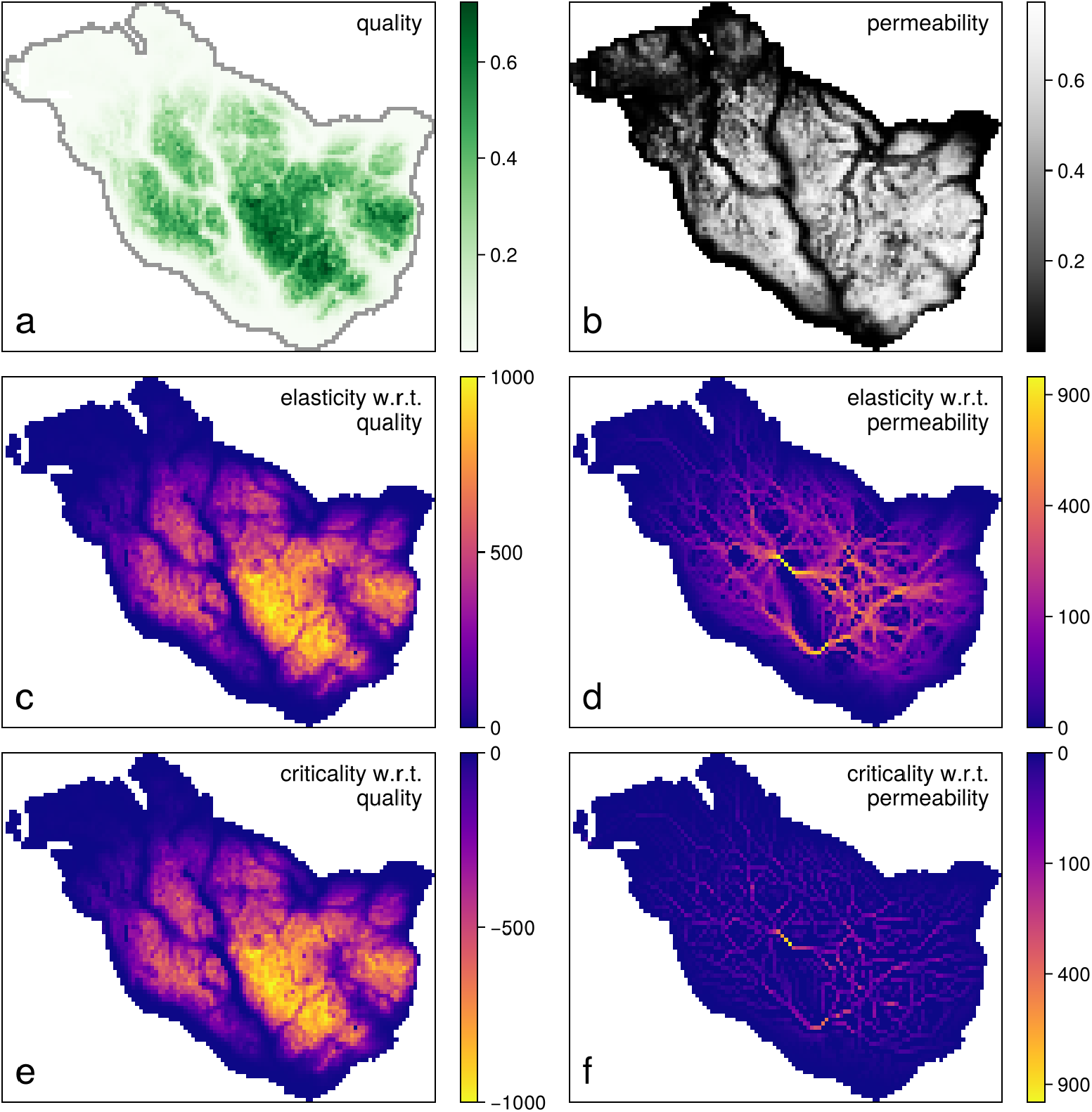
Demonstration and validation of the analytical sensitivity framework for the Snøhetta wild reindeer area in Norway. (a) Habitat quality and (b) permeability data for the network parameterization were taken from Van Moorter et al. (2023a). We computed the elasticity of landscape functionality – based on the summation formulation of the landscape matrix M – with respect to (c) habitat quality and (d) permeability. To validate these results, we compared them with two node-removal analysis to quantify the criticality of a node: one in which the habitat quality of a node was set to zero while retaining it as a connector (e), and another in which the node was completely removed from the network. The difference between these two node-removal analyzes isolates the effect of removing only a node’s connector role (f). See main text for details.

**Figure 3.**
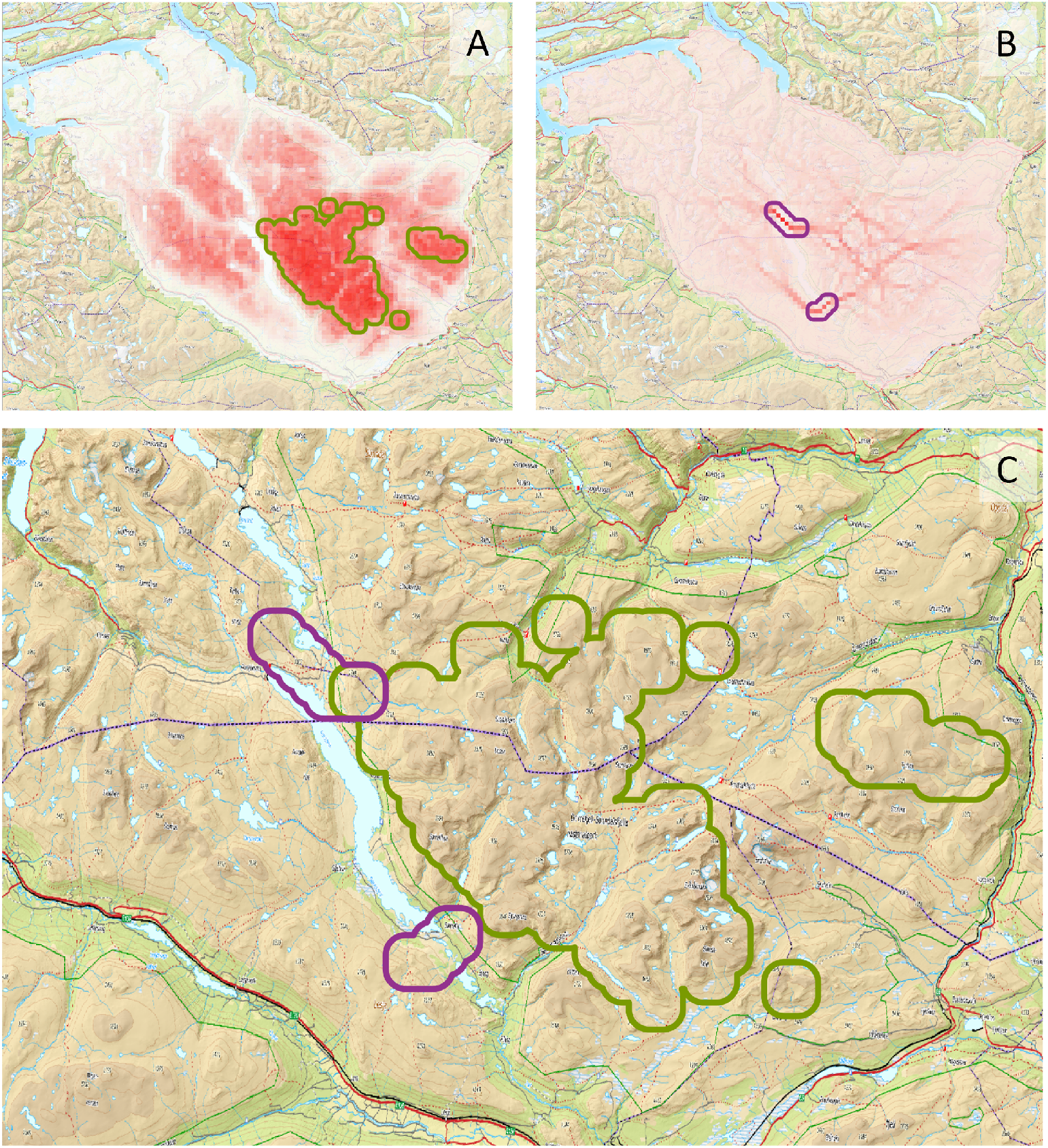
Spatial sensitivity analysis for conservation prioritization. Panels (a) and (b) show elasticities with respect to habitat quality and permeability, respectively; the most sensitive areas are highlighted in red. Panel (c) overlays the most influential areas (purple: permeability; green: quality) on a topographic map of the region. See main text for further discussion.

Prioritization based on elasticities assumes a linear response to infinitesimal perturbations and therefore does not represent the complete destruction of a spatial unit. We compared elasticity maps to node-removal analysis (Fig. 2e–f), in which landscape functionality is recomputed after systematically removing each unit and contributions are assessed based on the resulting loss in landscape functionality. Elasticity values showed strong negative correlations with the corresponding node-removal analysis (quality elasticity: *r* = −1.00; permeability elasticity: *r* = −0.70), confirming that the analytical approach closely reproduces the spatial vulnerability patterns revealed by iterative node-removal, while being orders of magnitude faster.

Analytical computation of elasticities for this landscape (≈5,000 nodes) required only 25 seconds, compared to approximately 12 hours (42,592 seconds) for the equivalent iterative node-removal analysis – a speed increase of more than three orders of magnitude. All computations were performed on a Windows 11 laptop equipped with a 13^th^ Gen Intel^®^ Core^™^ i7–1370P processor (14 cores, 1.9 GHz base frequency) and 16 GB RAM, using single-threaded execution without GPU acceleration. Further optimization using landmark-based computation can reduce runtime even further with minimal accuracy loss (Appendix E), demonstrating that the approach scales efficiently to larger, real-world landscapes.

## 4. Discussion

Connectivity is now a central concern in spatial conservation planning, reflected in the incorporation of connectivity metrics into widely used prioritization tools such as Zonation (Lehtomäki and Moilanen, 2013) and Marxan Connect (Daigle et al., 2020). Yet current existing implementations typically neglect the dual role of landscape units as both functional habitat and connectors (Saura and Rubio, 2010), rely on overly simplified movement models that may not reflect species-specific dispersal behaviour (McRae and Beier, 2007; Saura and Pascual-Hortal, 2007), or depend on computationally intensive iterative analysis that do not scale to relevant resolutions for spatial planning (Saura and Torne, 2009). Our work addresses these limitations by providing a unified, analytically tractable framework for assessing and prioritizing landscape functionality, grounded in metapopulation theory and movement ecology.

By generalizing the original landscape-matrix formulation (Hanski, 1999; Ovaskainen, 2003) to accommodate a broad class of movement models, we connect widely used connectivity metrics to explicit persistence conditions from metapopulation theory. Several established metrics – including neighbourhood habitat area index (Hanski, 1999), probability of connectivity (Saura and Pascual-Hortal, 2007), and equivalent connected area (Saura et al., 2011) – emerge as special cases of our framework when different ecological distances are used. The key distinction among these metrics lies in how proximity *k*_*st*_ is derived from the ecological distance *d*_*st*_. Species moving optimally with full knowledge of the landscape are best represented by least-cost distance; species moving randomly (e.g., exhibiting diffusive search behaviour) align with resistance distance from circuit theory; species whose movement lies between these extremes benefit from the RSP framework’s optimality parameter *θ*, which allows continuous tuning between optimal and random movement; and species experiencing mortality during dispersal are naturally represented by the survival probability derived from absorbing random walks (Fletcher Jr et al., 2019). This integrative view helps organise a wide array of connectivity measures within a single quantitative framework.

Importantly, our results link summation-based metrics to a persistence threshold based on metapopulation colonization extinction dynamics, therefore providing theoretical justification for their use in assessing landscape functionality. Summation-based summary of the landscape matrix lend themselves to spatial decomposition and moving-window computation (similar to Hughes et al., 2023), making them practical for large or spatially heterogeneous regions. In contrast, metapopulation capacity requires full matrix construction and eigenanalysis, which becomes computationally intractable for large landscapes. While metapopulation capacity has long been linked to persistence thresholds (Hanski and Ovaskainen, 2000), our derivation of a analogous persistence condition for the summation formulation offers a computationally efficient alternative that retains a direct connection to metapopulation dynamics.

We show that sensitivity analysis provides a principled framework for spatial prioritization. Sensitivity of landscape functionality to habitat quality corresponds to a closeness-like centrality metric, capturing how well a node is connected to all others, whereas sensitivity to permeability corresponds to a betweenness-like centrality metric, reflecting a node’s contribution to facilitating movement between other areas (Borgatti and Everett, 2006). These relationships explain why centrality metrics have often performed well as heuristic prioritization indicators for connectivity conservation (e.g. Hodgson et al., 2016; Daigle et al., 2020), while revealing that they are often simplified forms of the exact sensitivities or elasticities derived here. By deriving sensitivities across least-cost (Adriaensen et al., 2003), circuit-based (McRae and Beier, 2007), SAMC (Marx et al., 2020), and RSP formulations (Saerens et al., 2009), we reconcile movement models that have often been treated as conceptually distinct and show that they can be analysed within a single, differentiable framework.

The elasticity-based prioritization approach translates these analytical sensitivities into directly actionable maps for conservation. Elasticities identify spatial units where proportional changes in local habitat quality or permeability produce the largest proportional change in overall landscape functionality, thereby highlighting areas that disproportionately contribute to habitat availability and connectivity. Although the analytical sensitivities are derived under assumptions of linearity and independence among local perturbations, our results show that they remain informative even when these assumptions are partly violated. In the Snøhetta wild reindeer case study, elasticity maps based on habitat quality and permeability aligned closely with known ecological patterns and management priorities, recovering key movement corridors and highquality habitat areas identified in earlier expert-based assessments.The computational gains were substantial: sensitivity maps were obtained more than three orders of magnitude faster than iterative node-removal analysis, with additional speed-ups achievable through landmark-based approximations for very large landscapes.

The broader potential of analytical sensitivity extends beyond spatial conservation prioritization. Because our framework yields explicit gradients of connectivity metrics with respect to underlying parameters, it enables gradient-based optimization for improving the representation of permeability through, e.g., inverse landscape genetics (Peterman et al., 2019). By bridging network theory, metapopulation models, and applied spatial planning within a unified differentiable framework, our study advances the analytical foundations of landscape ecology and opens new avenues for integrating connectivity into evidence-based land-use planning and biodiversity conservation.

## 5. Conclusions

This study introduces a unifying framework for spatial prioritization of landscape functionality. The approach provides new insight into how local changes in habitat quality and permeability influence the landscape-scale connectivity – information essential for spatial conservation prioritization. In practice, sensitivity outputs can guide the early stages of scenario design by highlighting where conservation actions would most influence overall landscape functionality. Dedicated scenario analyses remain indispensable for evaluating the detailed outcomes of major interventions (Dorber et al., 2023), but sensitivity outputs can help shape, refine, and prioritize those scenarios. The framework offers an analytically transparent and computationally efficient means of spatial prioritization – more than three orders of magnitude faster than iterative node-removal analysis. Integrating sensitivity layers across species and embedding them within existing conservation planning tools can strengthen multi-species prioritization and improve connectivity management in fragmented landscapes. In particular, these layers could be incorporated into tools such as Marxan Connect (Daigle et al., 2020), replacing generic centrality metrics with sensitivity measures directly derived from the model’s objective function. By linking network theory, metapopulation models, and applied spatial planning within a unified differentiable framework, this work advances the analytical foundations of connectivity science and supports evidence-based strategies for biodiversity conservation and sustainable land use.

## Acknowledgments

The work was supported by grant 287925 from the Research Council of Norway.

## Data availability statement

Data are available in the ConScape library (Van Moorter et al., 2023a) and code is provided in Appendix D of the supplementary material.

