## appendices for "A unified framework for prioritizing habitat and connectivity conservation through analytical sensitivity"

#### Appendix A. Landscape functionality

Different metrics have been developed to quantify the amount and connectedness of habitat in a given landscape simultaneously (Saura and Pascual-Hortal, 2007; Kindlmann and Burel, 2008). The two major ones share a background in metapopulation ecology: extensions of the 'neighborhood habitat area index' (NHAI; pp. 83 in Hanski, 1999) and the 'metapopulation capacity' (MC; Hanski and Ovaskainen, 2000), which are based on respectively summation and eigenanalysis of the 'landscape Matrix' ($\mathbf{M}$). Note that we use the term 'matrix' in its mathematical meaning, not as the area in between habitat patches (cf. the patch-matrix representation of a landscape; Lausch et al., 2015). The matrix $\mathbf{M}$ consists of elements $m_{st} = q_s q_t k_{st}$, where $q$ is the quality and $k_{st}$ is the proximity between all pairs of areas as source $s$ and target $t$ (Van Moorter et al., 2021). The proximity is often derived from an (ecological) distance $d$ using an exponential function, $k_{st} = \exp(-\alpha d_{st})$, scaled with species-specific movement capabilities $\alpha$; however, several other distance decay functions can be considered (for a review see: Bullock et al., 2017). The difference between the NHAI and MC lies in the summary statistics derived from the landscape matrix $\mathbf{M}$, respectively the sum of its elements versus its dominant eigenvalue. Van Moorter et al. (2023b) defined 'functional habitat' ($\mathcal{K}(i)$) as a network-based concept to integrate the suitability of local environmental conditions – or habitat – in a spatial unit $i$ (e.g. pixel) with its connectivity to other suitable habitat derived from $\mathbf{M}$ (either through row- or column-wise summation or through the left or right eigenvectors). Van Moorter et al. (2023b) extended this pixel- or patch-based habitat functionality to the landscape level, as 'landscape functionality' ($\mathcal{K}(L)$).

We define 'landscape functionality' as a scalar metric quantifying the amount and interconnectedness of habitat across a landscape, derived from the landscape matrix $\mathbf{M}$ via summation or eigenanalysis. This generalises earlier quantities of connectivity or metapopulation persistence. We first discuss the summation-based landscape functionality metrics ($\mathcal{K}_{\text{sum}}(L)$) followed by the eigenvalue-based ones ($\mathcal{K}_{\text{eig}}(L)$).

The NHAI was the first measure of landscape functionality, $\mathcal{K}(L)$, based on the summation of $\mathbf{M}$:

$$\mathcal{K}_{\text{sum}}^{\text{Eucl}}(L) = \sum_{s=1}^{n} \sum_{t=1}^{n} q_s q_t \exp(-\alpha d_{st}^{\text{Eucl}}). \tag{A.1}$$

where $q$ is the size of a source $s$ and target $t$ patch and $d_{st}^{\text{Eucl}}$ is the Euclidean distance between these patches (Hanski, 1999). Unfortunately, Euclidean distance is often a poor approximation of the 'real' ecological distance (Sutherland et al., 2015), as barriers in the landscape may affect movement paths and their associated ecological distance.

Saura and Pascual-Hortal (2007) extended this metric by replacing the Euclidean distance with the ecological cost distance of the least-cost path between $s$ and $t$ to account for the spatial variation in cost of moving through the landscape:

$$\mathcal{K}_{\text{sum}}^{\text{LC}}(L) = \sum_{s=1}^{n} \sum_{t=1}^{n} q_s q_t \exp(-\alpha d_{st}^{\text{LC}}). \tag{A.2}$$

where $q$ can be either size or quality of a source $s$ and target $t$ patch or pixel and $d_{st}^{\text{LC}}$ is the least-cost distance between them. Saura and Pascual-Hortal (2007) used as a cost metric the logarithm of the movement probabilities between patches/pixels, therefore the exponential proximity is the product of these probabilities. This led Saura and Pascual-Hortal (2007) to interpret this metric as 'the probability that two animals randomly placed within the landscape fall into habitat areas that are reachable from each other', and they therefore named $\mathcal{K}_{\text{sum}}^{\text{LC}}(L)$ the 'probability of connectivity' (PC). This metric decreases with both habitat loss and/or fragmentation (Rubio and Saura, 2012).

By taking the square root of the PC, Saura et al. (2011) obtained a measurement of the 'equivalent connected area' $\left(ECA = \sqrt{\mathcal{K}_{\text{sum}}(L)}\right)$, where quality $q$ in $\mathbf{M}$ represents the area of the patches. The ECA of a landscape corresponds to the amount of habitat available, if it were to occur in a single perfectly connected patch. The main advantage of this metric is that it is expressed in area units, which makes it easier to interpret for land managers. A similar measure can be defined more generally using the quality of pixels, as the 'equivalent connected habitat' (Van Moorter et al., 2023a):

$$ECH = \sqrt{\mathcal{K}_{\text{sum}}(L)}. \tag{A.3}$$

Unfortunately, ECH is measured in 'habitat $\times$ area' units and not immediately in 'area' units as is the ECA and therefore requires an additional step to define habitat units (e.g. Dorber et al. 2023).

The use of the least-cost distance is an appreciable improvement over the Euclidean distance, as it accounts for barriers and stepping stones in between $s$ and $t$; however, it fails to account for the contribution of multiple paths to the connectedness of $s$ and $t$ by focusing exclusively on the cost of a single path. Hence, in connectivity studies, the random walk – closely related to circuit theory (McRae and Beier, 2007; McRae et al., 2008) – has become a popular approach to account for the number of pathways. Although, circuit theory has been used extensively to identify corridors and pinch points for connectivity on the landscape (Dickson et al., 2019), it has not found much usage in metrics of landscape functionality (but see: Avon and Bergès, 2016). Moreover, the random walk is not without its problems, as it suffers from the so called 'lost in space' phenomenon on large landscapes (Von Luxburg et al., 2010; Van Moorter et al., 2021). Instead of focusing exclusively either on the number or on the cost of paths, the Randomized Shortest Paths framework (RSP; Saerens et al., 2009) accounts simultaneously for the number of paths and the cost of those paths for the computation of $k_{st}$ in the $\mathbf{M}$.

An alternative approach to quantify landscape functionality using eigenanalysis focuses on the long-term persistence of a metapopulation from the extinction and recolonization dynamics of local population patches. Hanski and Ovaskainen (2000) showed that a metapopulation persists, if the 'metapopulation capacity' (MC) is above a species-specific threshold, where the MC is defined as the dominant eigenvalue, $\lambda$, of the landscape Matrix $\mathbf{M}$. Thus, the 'eigenvalue landscape functionality' is:

$$\mathcal{K}_{\text{eig}}(L) = \lambda(\mathbf{M}). \tag{A.4}$$

Originally, in Hanski and Ovaskainen (2000) the $q$ in $\mathbf{M}$ referred to the size of the patches and did not allow for self-colonization: i.e. $m_{ss} = 0$; further developments, however, dropped that assumption and allowed for self-rescue (Schnell et al., 2010): $m_{ss} = q_s^2$. Moreover, the proximity between patches $s$ and $t$ was originally derived from the Euclidean distance ($k_{st}^{\text{Eucl}} = \exp(-\alpha d_{st}^{\text{Eucl}})$), as with the NHAI above, this can easily be extended to ecological distances.

While the dominant eigenvalue of the landscape Matrix $\mathbf{M}$ has a solid foundation in metapopulation theory as a metric of long-term persistence of a metapopulation, it is computationally demanding for large landscapes. Fortunately, as presented by Song et al. (2022), the dominant eigenvalue ($\lambda(\mathbf{M})$, which is here also its spectral radius) of $\mathbf{M}$ (where $m_{st} \geq 0$ for all $s$-$t$), through the Perron-Frobenius theorem, satisfies:

$$\min_{1\le s\le n}\Big(\sum_{t=1}^{n} M_{st}\Big) \le \lambda(\mathbf{M}) \le \max_{1\le s\le n}\Big(\sum_{t=1}^{n} M_{st}\Big). \tag{A.5}$$

As the sum of all matrix entries is the sum of all row sums, the simple inequality above also implies:

$$\lambda(\mathbf{M}) \le \mathcal{K}_{\text{sum}}(L). \tag{A.6}$$

This bound is in general not tight. But because a landscape can be seen as a realisation of a stochastic process, entries of the landscape matrix are draws of a random variable. If these draws are sufficiently independent with finite mean and variance, the law of large numbers ensures that row sums concentrate near their mean $\mathcal{K}_{\text{sum}}(L)/n$:

$$\min_{1\le s\le n}\Big(\sum_{t=1}^{n} M_{st}\Big) \approx \lambda(\mathbf{M}) \approx \max_{1\le s\le n}\Big(\sum_{t=1}^{n} M_{st}\Big) \to \mathcal{K}_{\text{sum}}(L)/n \;\text{ as }\; n\to\infty, \tag{A.7}$$

Hence for large landscapes, $\mathcal{K}_{\text{sum}}(L)/n$ is an approximation of the metapopulation capacity.

Similarly, as shown in Appendix B, the summation-based landscape functionality $\mathcal{K}_{\text{sum}}(L)$ can be directly linked to metapopulation theory through a stochastic colonization–extinction model (Hanski, 1999; Hanski and Ovaskainen, 2000). In this framework, the condition

$$\frac{\mathcal{K}_{\text{sum}}(L)}{n\overline{q^{-1}}} > \frac{e}{c} \tag{A.8}$$

expresses the macroscopic mean-field persistence threshold for a metapopulation distributed across a heterogeneous landscape. Persistence occurs when the landscape-scale connectivity – represented by the summation-based functionality $\mathcal{K}_{\text{sum}}(L)$ (with a correction factor $1/(n\overline{q^{-1}})$) – exceeds the ratio of extinction to colonization rates ($e/c$). This mirrors Hanski & Ovaskainen's classical condition $\lambda(\mathbf{M}) > e/c$, where $\lambda(\mathbf{M})$ is the dominant eigenvalue (the metapopulation capacity) of the landscape matrix $\mathbf{M}$. It thus establishes $\mathcal{K}_{\text{sum}}(L)$ as an alternative but theoretical grounded metric of metapopulation persistence.

#### Appendix B. Mean-field approximation for metapopulations in a heterogeneous environment

##### *Appendix B.1. Purpose and context*

The purpose of this appendix is to place the summation-based landscape functionality $\mathcal{K}(L) = \sum_t \sum_s k_{st} q_s q_t$ in the formal context of metapopulation theory. We show that summation-based $\mathcal{K}(L)$ arises naturally in a landscape-level mean-field approximation of a stochastic colonization–extinction model, and that it plays the same structural role as the dominant eigenvalue – metapopulation capacity – in Hanski's formulation (Hanski and Ovaskainen, 2000). This establishes a theoretical basis for using summation-based $\mathcal{K}(L)$ as an indicator of landscape-level persistence potential and for computing its sensitivities to environmental change. We note that Hanski's formulation also rests on a mean-field assumption, but at the patch instead of the landscape scale. The advantage of working at the landscape scale is that we will not have to calculate eigenvalues of a large matrix.

##### *Appendix B.2. Notation and definitions*

We consider a lattice of $n$ cells. Each cell $i$ has an occupancy variable $o_i \in \{0, 1\}$, equal to 1 when occupied and 0 when unoccupied.

*Dispersal kernel.* The weight $k_{st}$ describes the relative contribution of source cell $s$ to target cell $t$ and satisfies $\forall s : \sum_{t=1}^{n} k_{st} = 1$. For mathematical simplicity we assume $k_{tt} = 0$, meaning that colonization of a site cannot originate from itself. This allows all sums to be written over $s$ without explicitly excluding $s = t$; the diagonal terms never contribute to colonization because they multiply $(1 - o_t)$, which is zero when $o_t = 1$.

From an ecological perspective, however, some formulations (e.g., Ovaskainen and Cornell, 2006; Schnell et al., 2010, 2013) allow $k_{tt} > 0$ to represent self-colonization – local persistence or demographic rescue following temporary extinction. Including such terms would add small diagonal contributions $k_{tt} q_t^2$ to $\mathcal{K}(L)$ but does not alter the qualitative persistence condition derived below.

*Reaction rules.* Change in occupancy state $o$ of cell $t$ through colonization and extinction are assumed to occur probabilistically following the processes (Hanski, 1999):

$$o_t = \begin{cases} 0 \to 1 & \text{at rate } c_t(1 - o_t) \sum_{s=1}^{n} k_{st} q_s o_s, \\ 1 \to 0 & \text{at rate } e_t, \end{cases} \tag{B.1}$$

here $q_s$ measures habitat quality or fecundity at the source, $c_t$ recruitment potential at the target, and $e_t$ local extinction rate.

*Ensemble and spatial means.* Because these reactions are stochastic, $o_t$ is a random variable. The ensemble mean (average over realizations) of cell occupation is equal to the probability of a cell being occupied i.e., $\langle o_t \rangle = \mathbb{E}[o_t] = P(o_t{=}1)$ because

$$\langle o_t \rangle := \sum_{x \in \{0,1\}} x P(o_t{=}x) = 0 \times P(o_t{=}0) + 1 \times P(o_t{=}1) = P(o_t{=}1), \tag{B.2}$$

The spatial mean over all cells is

$$\bar{o} := \frac{1}{n} \sum_{t=1}^{n} o_t$$

Angular brackets $\langle . \rangle$ denote ensemble averages; overbars denote spatial averages.

*Appendix B.3. Macroscopic mean field approximation*

The expected occupancy of cell $t$ evolves as:

$$\frac{\mathrm{d}}{\mathrm{d}\tau}\langle o_t \rangle = c_t \sum_s k_{st} q_s \langle o_s (1 - o_t) \rangle - e_t \langle o_t \rangle. \tag{B.3}$$

Summing over all cells gives a macroscopic equation for the mean occupancy:

$$n \frac{\mathrm{d}}{\mathrm{d}\tau}\langle \bar{o} \rangle = \sum_t c_t \sum_s k_{st} q_s \langle o_s (1 - o_t) \rangle - \sum_t e_t \langle o_t \rangle. \tag{B.4}$$

Starting from the macroscopic equation (B.4), we make a series of simplifying assumptions that permit analysis. We first define the following quantities:

$$\overline{M} := n^{-2} \sum_t \sum_s k_{st} q_s q_t, \tag{B.5}$$

$$\overline{\langle o_s \rangle (1 - \langle o_t \rangle)} := n^{-2} \sum_t \sum_s \langle o_s \rangle (1 - \langle o_t \rangle), \tag{B.6}$$

$$\overline{q^{-1}} := n^{-1} \sum_t 1/q_t. \tag{B.7}$$

The assumptions we will make are:

1. Pairwise independence between cells among ensembles:

$$\langle o_s (1 - o_t) \rangle \approx \langle o_s \rangle (1 - \langle o_t \rangle), \tag{B.8}$$

2. $c_t = c q_t$ and $e_t = e/q_t$,
3. Statistical independence in space between parameters and occupation:

$$n^{-2} \sum_t \sum_s k_{st} q_s q_t \langle o_s \rangle (1 - \langle o_t \rangle) \approx \overline{M}\ \overline{\langle o_s \rangle (1 - \langle o_t \rangle)}, \tag{B.9}$$

$$n^{-1} \sum_t o_t / q_t \approx \overline{q^{-1}} \bar{o}, \tag{B.10}$$

4. Statistical independence of occupation in space:

$$\overline{\langle o_s \rangle (1 - \langle o_t \rangle)} \approx \langle \bar{o} \rangle (1 - \langle \bar{o} \rangle), \tag{B.11}$$

Applying these standard mean-field assumptions, equation (B.4) reduces to:

$$\frac{\mathrm{d}}{\mathrm{d}\tau}\langle \bar{o} \rangle = c n \overline{M} \langle \bar{o} \rangle (1 - \langle \bar{o} \rangle) - e \overline{q^{-1}} \langle \bar{o} \rangle. \tag{B.12}$$

This equation describes the deterministic mean-field dynamics of the spatially averaged occupancy.

*Appendix B.4. Persistence threshold and connection to summation-based $\mathcal{K}(L)$*

Linearizing (B.12) near extinction ($\langle\bar{0}\rangle \approx 0$) gives a Jacobian:

$$J = cn\overline{M} - e\overline{q^{-1}}, \tag{B.13}$$

The trivial state becomes unstable —- and the metapopulation persists —- when

$$n\overline{M}/\overline{q^{-1}} > e/c. \tag{B.14}$$

Using summation-based $\mathcal{K}(L) = n^2\overline{M}$, this can be written:

$$\frac{\mathcal{K}(L)}{n\overline{q^{-1}}} > e/c. \tag{B.15}$$

This mirrors Hanski's classical condition $\lambda > e/c$, where $\lambda$ is the dominant eigenvalue (the metapopulation capacity) of the landscape matrix $\mathbf{M}$.

Here $\mathcal{K}(L)/(nq^{-1})$ plays the role of an aggregated mean-field metapopulation capacity. Its derivative $\partial\mathcal{K}(L)/\partial x_i$ therefore quantifies how sensitive persistence is to parameter $x_i$, directly analogous to the eigenvalue sensitivities in Hanski and Ovaskainen's framework.

*Appendix B.5. Limiting cases and consistency checks*

When mortality is spatially homogeneous ($\forall t : e_t = e$), the condition simplifies to

$$\frac{\mathcal{K}(L)}{n} > e/c. \tag{B.16}$$

When instead fecundity is homogeneous ($\forall t : c_t = c$), it becomes

$$\frac{\overline{q}}{\overline{q^{-1}}} > e/c. \tag{B.17}$$

In a fully uniform environment ($\forall t : c_t = c,\ q_t = 1$), this reduces to the classical mean-field threshold $c/e > 1$. These limits confirm that the mean-field approximation is consistent with established metapopulation theory.

*Appendix B.6. Derivatives of the persistence term*

The persistence term in the mean-field threshold (Eq. B.15) is:

$$\Phi = \frac{\mathcal{K}(L)}{n\overline{q^{-1}}}, \tag{B.18}$$

with $\mathcal{K}(L) = \sum_t \sum_s k_{st} q_s q_t$ and $\overline{q^{-1}} = \frac{1}{n}\sum_t \frac{1}{q_t}$.

*Derivative with respect to habitat quality.* Differentiating $\Phi$ with respect to the quality $q_t$ of a given cell gives:

$$\frac{\partial \Phi}{\partial q_t} = \frac{1}{n\overline{q^{-1}}} \frac{\partial \mathcal{K}(L)}{\partial q_t} - \frac{\mathcal{K}(L)}{n(\overline{q^{-1}})^2} \frac{\partial \overline{q^{-1}}}{\partial q_t}. \tag{B.19}$$

The first term represents the direct contribution of site $t$ to the overall structural connectivity, while the second term accounts for how changes in $q_t$ alter $\overline{q^{-1}}$. From the definition of $\overline{q^{-1}}$:

$$\frac{\partial \overline{q^{-1}}}{\partial q_t} = -\frac{1}{n q_t^2}. \tag{B.20}$$

and hence

$$\frac{\partial \Phi}{\partial q_t} = \frac{1}{n\overline{q^{-1}}} \frac{\partial \mathcal{K}(L)}{\partial q_t} + \frac{\mathcal{K}(L)}{n^2(\overline{q^{-1}})^2 q_t^2}. \tag{B.21}$$

The second term decreases in magnitude roughly as $1/n$ and becomes negligible in large landscapes, unless quality heterogeneity is extreme or $q_t$ is very small.

*Derivative with respect to habitat permeability.* Differentiating $\Phi$ with respect to the permeability $a_t$ of a given cell gives:

$$\frac{\partial \Phi}{\partial a_t} = \frac{1}{n\overline{q^{-1}}} \frac{\partial \mathcal{K}(L)}{\partial a_t} - \frac{\mathcal{K}(L)}{n(\overline{q^{-1}})^2} \frac{\partial \overline{q^{-1}}}{\partial a_t}. \tag{B.22}$$

The second term is zero, hence:

$$\frac{\partial \Phi}{\partial a_t} = \frac{1}{n\overline{q^{-1}}} \frac{\partial \mathcal{K}(L)}{\partial a_t}. \tag{B.23}$$

So the sensitivity of $\mathcal{K}(L)$ to the permeability is multiplied by the constant factor $1/(n\overline{q^{-1}})$. This rescales magnitudes but does not change rankings or spatial patterns, since $n\overline{q^{-1}}$ doesn't depend on habitat permeability.

##### *Appendix B.7. Interpretation summary*

This appendix shows that the summation-based $\mathcal{K}(L)$ is not merely a heuristic aggregation of the landscape matrix M. Within a landscape-level mean-field framework, it represents the landscape's aggregate capacity to sustain occupancy, playing an analogous structural role to Hanski & Ovaskainen's metapopulation capacity. The resulting persistence threshold, based on $\mathcal{K}(L)$, is mathematically parallel to the eigenvalue formulation but easier to compute and differentiate, providing a transparent bridge between network-based metrics and metapopulation theory.

The full persistence term $\Phi = \mathcal{K}(L)/(n\overline{q^{-1}})$ includes a correction factor based on the mean of inverse habitat quality. This factor is ecologically interpretable – it emphasizes low-quality sites – but its quantitative effect on sensitivities is small except under extreme heterogeneity. For the purposes of the main text, we therefore focus on sensitivities of $\mathcal{K}(L)$ itself, which capture the dominant structural dependence on permeability and habitat quality. Correction factors in the denominator are neglected unless explicitly stated.

#### Appendix C. Sensitivity derivations

In this Appendix, we present the derivations of the sensitivity of the 'landscape functionality' (see Appendix A) w.r.t. (with respect to) habitat degradation and habitat fragmentation. It extends results published in (Kivimäki et al., 2024) by considering measures defined on the whole landscape instead of simple $s$-$t$ (single source and target) points on that landscape. The new derived results of this appendix therefore rely on previous results, developed in that paper. First, in Section Appendix C.1, we provide the formal definition of all concepts and symbols using graph theory. In subsequent Sections Appendix C.2, Appendix C.4, Appendix C.5, and Appendix C.6 we present the actual derivations of the sensitivity of the landscape functionality w.r.t. quality and permeability. Section Appendix C.2 presents the derivations for the expected-cost in the randomized shortest paths framework and the survival probability from absorbing random walks (the algorithms are presented in section Appendix C.3) for summation-based landscape functionality metrics, section Appendix C.4 extends these results to eigen-based landscape functionality metrics. Sections Appendix C.5 and Appendix C.6 derive the sensitivity for the circuit and the least-cost-path model, respectively.

##### *Appendix C.1. Definitions and notation*

We derived the sensitivity for the *Randomized Shortest Paths* framework (RSP; Yen et al., 2008; Saerens et al., 2009), which interpolates between the most commonly used distances (i.e. least-cost and hitting-cost distance) using an optimality parameter $\theta$ (for an ecological discussion see: Van Moorter et al., 2021). In addition to the RSP framework generalizing prior ecological distances (Van Moorter et al., 2021), all the quantities involved in the RSP model are differentiable (in contrast to the minimizing function from the least-cost distance Klein, 2010), which facilitates a sensitivity analysis. We now introduce the RSP framework following Van Moorter et al. (2021), and present the notation and definition of all symbols used for the sensitivity analysis (Table C.2).

The RSP framework was inspired by traffic models developed in transportation sciences (Akamatsu, 1996), and can be used to compute the proximity $k_{st}$ for $\mathbf{M}$ based on, for instance, the 'expected cost' or the 'survival probability' (Kivimäki et al., 2014; Van Moorter et al., 2021). Two elements contribute to the permeability of a landscape element: the *likelihood* of the agent traversing it, and the energetic or survival *cost* of doing so (Fletcher Jr et al., 2019; Van Moorter et al., 2021). The RSP framework uses both the natural random walk transition probabilities ($p_{ij} = a_{ij}/d_{i\rightarrow}$; where we call $a_{ij}$ the step probability between $i$ and $j$ and $d_{i\rightarrow} = \sum_k a_{ik}$ is the out-degree of node $i$) and the cost ($c_{ij}$) of moving between two adjacent nodes $i$ and $j$. The step probability and cost of a transition can be linked, e.g. through a logarithmic transformation ($c_{ij} = -\log a_{ij}$), or they can be independent variables. Recently, killed or absorbing random walks (Madras, 1989) have been introduced to ecology (Fletcher Jr et al., 2019), where the cost of movement is related to the mortality risk. Under optimal movement, the likelihood and cost of movement are negatively related; however, Fletcher Jr et al. (2019) showed that movement likelihood can be positively related to mortality, giving rise to a 'dispersal trap'. The RSP is a generalized version of the model presented by Fletcher Jr et al. (2019); by including a scaling parameter ($\theta$) it allows for the interpolation between pure random walk (i.e. without mortality), over absorbing/killed random walks (Fletcher Jr et al., 2019), to optimal movements (i.e. least-cost paths). We refer to Van Moorter et al. (2021) and Van Moorter et al. (2023a) for a further description and computation of the RSP framework in an ecological context.

Following Van Moorter et al. (2023b), we extend the previous Euclidean or least-cost distance-based landscape functionality metrics, $\mathcal{K}(L)$, with two alternative proximity metrics derived from the RSP framework: (1) the survival probability quantifying the accessibility between node $s$ and node $t$, $\mathcal{Z}_{st}$ (see: Van Moorter et al., 2021, 2023a):

$$k_{st}^{\mathrm{Surv}} = \mathcal{Z}_{st} = \sum_{\wp \in \mathcal{P}_{st}} \mathrm{P}^{\mathrm{rw}}(\wp) \exp(-\theta c(\wp)), \tag{C.1}$$

where survival probabilities can be easily computed from the fundamental matrix (see, e.g., Kivimäki et al. (2024), Equations (28)-(29)), and (2) the proximity-transformed expected cost, $k_{st}^{\mathrm{EC}}$:

$$k_{st}^{\mathrm{EC}} = \exp(-\alpha \bar{c}_{st}), \tag{C.2}$$

where $\bar{c}_{st} = \sum_{\wp \in \mathcal{P}_{st}} \mathrm{P}^{\mathrm{RSP}}(\wp) c(\wp)$ is the expected cost for reaching target node $t$ from source node $s$ along paths $\wp \in \mathcal{P}_{st}$ and $\alpha$ is a scaling factor; see Van Moorter et al. (2021, 2023a); Kivimäki et al. (2024) for further discussion.

Using the survival probability from the RSP framework, we can define landscape functionality as:

$$\mathcal{K}_{\mathrm{sum}}^{\mathrm{Surv}}(L) = \sum_{s=1}^{n} \sum_{t=1}^{n} q_s q_t k_{st}^{\mathrm{Surv}}, \tag{C.3}$$

where $q$ is the quality of source $s$ and target $t$ node, and $k_{st}^{\mathrm{Surv}} = \mathcal{Z}_{st}$ is the proximity between $s$ and $t$ based on the survival probability from $s$ to $t$ (see Equation (C.1)). Also the eigenvalue-based landscape functionality can be computed using the survival probability, $\mathcal{K}_{\mathrm{eig}}^{\mathrm{Surv}}(L)$.

Alternatively, from the RSP, we can also derive the proximity between $s$ and $t$ from the expected cost (EC) of moving from $s$ to $t$ according to the RSP distribution, denoted by $\bar{c}_{st}$. The EC, $\bar{c}_{st}$, can be transformed into a proximity $k_{st}^{\mathrm{EC}}$ by some decreasing function, and the corresponding landscape functionality is:

$$\mathcal{K}_{\mathrm{sum}}^{\mathrm{EC}}(L) = \sum_{s=1}^{n} \sum_{t=1}^{n} q_s q_t k_{st}^{\mathrm{EC}}. \tag{C.4}$$

In this work, we consider a negative exponential, Eq. (C.2), for the proximity transformation. As above, the eigenvalue-based landscape functionality, $\mathcal{K}_{\mathrm{eig}}^{\mathrm{EC}}(L)$, can be computed also using the EC.

Relying on previous work (Kivimäki et al., 2024), the next sections present the sensitivity analysis of these four RSP-based landscape functionality metrics ($\mathcal{K}_{\mathrm{sum}}^{\mathrm{Surv}}$, $\mathcal{K}_{\mathrm{sum}}^{\mathrm{EC}}$, $\mathcal{K}_{\mathrm{eig}}^{\mathrm{Surv}}$, and $\mathcal{K}_{\mathrm{eig}}^{\mathrm{EC}}$) based on the derivative with respect to $q$, $a$, and $c$. These derivatives provide important information of the impact of the quantities on the landscape functionality measures, quantifying the overall quality of the landscape.

See Table C.2 for the notation and definition of all symbols used for the sensitivity analysis. In this work, a landscape is represented as graph $L = (\mathcal{V}, \mathcal{E})$ consisting of a set of $n$ nodes (or cells, or vertices) identified by natural numbers, $\mathcal{V} = \{1, 2, \ldots, n\}$, that are abstractions of landscape patches corresponding to habitat areas or pixels on a raster map of the landscape – and a set of edges (or arcs) defined as ordered pairs of nodes $\mathcal{E} = \{(i, j)\}$, where an edge from node $i$ to node $j$ exists if the landscape patch represented as node $i$ is connected to (adjacent to) the patch represented by node $j$. For any node $i \in \mathcal{V}$, we denote by $ⓘ\!\rightarrow$ the set of *successor nodes* of $i$, i.e. nodes for which an edge from $i$ exists, and by $\rightarrow\!ⓘ$ the set of *predecessor nodes* of $i$, i.e. nodes from which an edge to $i$ exists; formally

$$ⓘ\!\rightarrow = \{k \in \mathcal{V} \mid (i, k) \in \mathcal{E}\} \tag{C.5}$$

$$\rightarrow\!ⓘ = \{k \in \mathcal{V} \mid (k, i) \in \mathcal{E}\}. \tag{C.6}$$

| Term & definition | Description |
|---|---|
| $q_i$ | quality of node $i$ |
| $c_j$ | cost of movement into node $j$ (non-negative) |
| $a_j$ | step likelihood (random walk likelihood of movement into node $j$ – a non-negative affinity) |
| $\theta$ | randomness parameter (inverse of the temperature of the system, $T$) |
| $\rightarrow\!ⓙ$ | set of predecessor nodes of $j$ |
| $ⓘ\!\rightarrow$ | set of successor nodes of $i$ |
| $p_{ij} = a_j \Big/ \sum_{k \in ⓘ\rightarrow} a_k$ | unbiased, reference, transition probability over edge $(i, j)$ |
| $k_{st}^{\text{Surv}} = \mathcal{Z}_{st}$ | survival probability proximity |
| $\bar{c}_{st}$ | expected cost of paths w.r.t. the RSP distribution from $s$ to $t$ |
| $\alpha$ | scaling factor of expected costs |
| $k_{st}^{\text{EC}} = \exp(-\alpha \bar{c}_{st})$ | expected cost proximity |
| $m_{st} = q_s \cdot k_{st} \cdot q_t$ | landscape matrix $\mathbf{M}$ where $k_{st}$ is either $k_{st}^{\text{Surv}}$ or $k_{st}^{\text{EC}}$ |
| $\mathcal{K}_{\text{sum}}(L) = \sum_{s,t=1}^{n} m_{st}$ | summation-based landscape functionality where $\mathcal{K}_{\text{sum}}$, $m_{st}$, are $\mathcal{K}_{\text{sum}}^{\text{Surv}}$, $m_{st}^{\text{Surv}}$ or $\mathcal{K}_{\text{sum}}^{\text{EC}}$, $m_{st}^{\text{EC}}$ |
| $\mathcal{K}_{\text{eig}}(L) = \lambda(\mathbf{M})$ | eigenvalue landscape functionality where $\lambda$ is the dominant eigenvalue of $\mathbf{M}$ |
| $n_{ij}(\wp)$ | number of times that path $\wp$ traverses edge $(i, j)$ |
| $c(\wp)$ | total cost along path $\wp$ |
| $\overline{n}_{ij}^{st}$ | expected number of passages over edge $(i, j)$ when considering $s$-$t$ hitting paths |
| $\overline{n}_{i\rightarrow}^{st} = \sum_{j \in ⓘ\rightarrow} \overline{n}_{ij}^{st}$ | expected number of passages *out of* node $i$ |
| $\overline{n}_{\rightarrow j}^{st} = \sum_{i \in \rightarrowⓙ} \overline{n}_{ij}^{st}$ | expected number of passages *into* node $j$ |
| $\mathring{n}_{ij}^{st}$, $\mathring{n}_{i\rightarrow}^{st}$, $\mathring{n}_{\rightarrow j}^{st}$ | same expected number of passages, but now considering regular paths |
| $p_{ij}^{(\text{t})} = \overline{n}_{ij}^{st} / \overline{n}_{i\rightarrow}^{st}$ | the RSP transition probability over edge $(i, j)$, biased towards target node $t$ |
| $\sigma_{ij}^{st}$ | negative covariance between $n_{ij}(\wp)$ and $c(\wp)$ when considering hitting paths |
| $\sigma_{i\rightarrow}^{st} = \sum_{j \in ⓘ\rightarrow} \sigma_{ij}^{st}$ | negative covariance between $n_{i\rightarrow}(\wp)$ and $c(\wp)$ |
| $\sigma_{\rightarrow j}^{st} = \sum_{i \in \rightarrowⓙ} \sigma_{ij}^{st}$ | negative covariance between $n_{\rightarrow j}(\wp)$ and $c(\wp)$ |
| $\mathring{\sigma}_{ij}^{st}$, $\mathring{\sigma}_{i\rightarrow}^{st}$, $\mathring{\sigma}_{\rightarrow j}^{st}$ | negative covariances when considering regular paths |

Table C.2: Notation; all the expected values and covariances are considered w.r.t. the RSP distribution over paths (both hitting, absorbing, and regular, non-hitting paths), from source node $s$ to target node $t$. These quantities can readily be computed from the fundamental matrix in the RSP framework; see (Kivimäki et al., 2024) for details.

Each node $j \in \mathcal{V}$ of the landscape graph $L$ is associated with three quantities; a *quality*, $q_j$, a step probability, $a_j$, and a cost, $c_j$. Therefore, it is assumed here that the step probability and the cost associated to each edge $(i, j)$, $a_{ij}$ and $c_{ij}$, are simply provided by the end node $j$ (the successor node), that is, $a_{ij} = a_j$ and $c_{ij} = c_j$. In other words, in this section, the step probabilities and the costs are defined on the nodes[1]. The step probabilities and costs will be

[1]Note that this assumption will be removed for the derivation of the algorithms, assuming there that these quantities are defined on the edges; see Section Appendix C.3.

used to define the *proximity*, $k_{st}$, between any two nodes $s$ and $t$ on the landscape graph $L$ according to the RSP framework (Van Moorter et al., 2021). These quantities then define the *landscape matrix*, $\mathbf{M}$, whose elements are $m_{st} = q_s{\cdot}q_t{\cdot}k_{st}$, thus the rows representing the source $s$ and the columns the target $t$ of the paths. As already mentioned before, for any proximity measure $k$ on $L$, the *summation-based landscape functionality index* is defined as $\mathcal{K}_{\text{sum}}(L) = \sum_{s,t=1}^{n} m_{st}$, and the *eigenvalue landscape functionality index* as the dominant eigenvalue of matrix $\mathbf{M}$.

Throughout the work, we assume for each parameter that the parameter values associated to any two nodes are defined independently of each other, i.e. that for any $i, j \in \mathcal{V}$, such that $i \neq j$, we assume $x_i \perp x_j$, where $x$ is either $q$, $c$, or $a$. Moreover, in general, we assume for each node $j \in \mathcal{V}$ that $q_j \perp a_j$ and $q_j \perp c_j$. Instead, $a_j$ and $c_j$ can either depend on each other or not. If $a_j$ and $c_j$ are defined independently of one another, then the appropriate way of quantifying the sensitivity of a landscape functionality index with respect to one of them is by computing the partial derivative w.r.t. said quantity, e.g. for the summation-based index as

$$\frac{\partial \mathcal{K}_{\text{sum}}(L)}{\partial x_j} = \frac{\partial}{\partial x_j}\left(\sum_{s,t=1}^{n} q_s q_t k_{st}\right) = \sum_{s,t=1}^{n} \frac{\partial (q_s q_t k_{st})}{\partial x_j}, \tag{C.7}$$

where $\mathcal{K}$ is either $\mathcal{K}_{\text{sum}}$ or $\mathcal{K}_{\text{eig}}$ and $x_j$ is one of parameters $q_j$, $a_j$ or $c_j$, associated to node $j$. Instead, in cases where $a_j$ and $c_j$ are defined based on one another, it is more appropriate to consider the total derivative, which takes into account the effect on one parameter caused by perturbation of the other. This is done in Section Appendix C.2.4. When considering perturbation of $q_j$, the total and partial derivatives are equivalent due to the assumption of independence between $q_j$ and all other parameters.

For the eigensensitivity analysis, note that matrix $\mathbf{M}$ is square, non-negative and irreducible. Therefore, based on the Perron-Frobenius Theorem (see, e.g. Meyer, 2000), $\mathbf{M}$ has a unique dominant eigenpair, i.e. an eigenvalue $\lambda$ with modulus larger than that of any other eigenvalue, and unique left and right eigenvectors, $\mathbf{v}$ and $\mathbf{w}$, respectively, in addition to which $\lambda$, $\mathbf{v}$ and $\mathbf{w}$ are guaranteed to be real and positive. In ecology, when $\mathbf{M}$ represents e.g. a population projection matrix (as in Caswell, 2019), its left eigenvector represents the stable distribution, whereas its right eigenvector represents the reproductive value. Ovaskainen and Hanski (2003) adopted the same terminology for metapopulations and refer to the stable distribution over and reproductive value of population patches. Note that the matrix defined by, e.g., Ovaskainen and Hanski (2003) and Caswell (2019) is transposed compared to here, hence the left $\mathbf{v}$ and right $\mathbf{w}$ eigenvector in their work represent respectively the reproductive value and stable distribution (i.e. the opposite of here).

In the following, we derive the formulae for computing the sensitivity analysis expressions presented in Table C.3.

##### *Appendix C.2. Sensitivity of summation-based landscape functionality*

The expressions for the sensitivity of the summation-based landscape functionality, $\mathcal{K}_{\text{sum}}(L)$ in Table C.3, are obtained by inserting the expressions derived in this section of the partial sensitivities of the proximities $k^{\text{Surv}}$ and $k^{\text{EC}}$ w.r.t. $q_j$, $a_j$, $c_j$, and the total sensitivity w.r.t. $c_j$ into Equation (C.7).

| Connectivity | Proximity | w.r.t. | Expression |
|---|---|---|---|
| | | $q$ | $\frac{\mathrm{d}\mathcal{K}^{\mathrm{Surv}}_{\mathrm{sum}}(L)}{\mathrm{d}q_j} = \sum_{t=1}^{n} q_t \cdot \left(k^{\mathrm{Surv}}_{tj} + k^{\mathrm{Surv}}_{jt}\right)$ |
| | $k^{\mathrm{Surv}}$ | $c$ | $\frac{\partial\mathcal{K}^{\mathrm{Surv}}_{\mathrm{sum}}(L)}{\partial c_j} = -\theta \cdot \sum_{s,t=1}^{n} q_s \cdot q_t \cdot k^{\mathrm{Surv}}_{st} \cdot \overline{n}^{st}_{\to j}$ |
| | | $a$ | $\frac{\partial\mathcal{K}^{\mathrm{Surv}}_{\mathrm{sum}}(L)}{\partial a_j} = \frac{1}{a_j} \cdot \sum_{s,t=1}^{n} q_s \cdot q_t \cdot k^{\mathrm{Surv}}_{st} \cdot \left(\overline{n}^{st}_{\to j} - \sum_{i \in \to\textcircled{j}} \overline{n}^{st}_{i\to} \cdot p_{ij}\right)$ |
| summation | | $a = e^{-c}$ | $\frac{\mathrm{d}\mathcal{K}^{\mathrm{Surv}}_{\mathrm{sum}}(L)}{\mathrm{d}c_j} = -\sum_{s,t=1}^{n} q_s \cdot q_t \cdot k^{\mathrm{Surv}}_{st} \cdot \left((\theta+1)\overline{n}^{st}_{\to j} - \sum_{i \in \to\textcircled{j}} \overline{n}^{st}_{i\to} \cdot p_{ij}\right)$ |
| | | $q$ | $\frac{\mathrm{d}\mathcal{K}^{\mathrm{EC}}_{\mathrm{sum}}(L)}{\mathrm{d}q_j} = \sum_{t=1}^{n} q_t \cdot \left(k^{\mathrm{EC}}_{tj} + k^{\mathrm{EC}}_{jt}\right)$ |
| | $k^{\mathrm{EC}}$ | $c$ | $\frac{\partial\mathcal{K}^{\mathrm{EC}}_{\mathrm{sum}}(L)}{\partial c_j} = -\alpha \sum_{s,t=1}^{n} q_s \cdot q_t \cdot k^{\mathrm{EC}}_{st} \cdot \left(\overline{n}^{st}_{\to j} + \theta \cdot \sigma^{st}_{\to j}\right)$ |
| | | $a$ | $\frac{\partial\mathcal{K}^{\mathrm{EC}}_{\mathrm{sum}}(L)}{\partial a_j} = \frac{\alpha}{a_j} \cdot \sum_{s,t=1}^{n} q_s \cdot q_t \cdot k^{\mathrm{EC}}_{st} \cdot \left(\sigma^{st}_{\to j} - \sum_{i \in \to\textcircled{j}} \sigma^{st}_{i\to} \cdot p_{ij}\right)$ |
| | | $a = e^{-c}$ | $\frac{\mathrm{d}\mathcal{K}^{\mathrm{EC}}_{\mathrm{sum}}(L)}{\mathrm{d}c_j} = -\alpha \cdot \sum_{s,t=1}^{n} q_s \cdot q_t \cdot k^{\mathrm{EC}}_{st} \cdot \left(\overline{n}^{st}_{\to j} + (\theta+1)\sigma^{st}_{\to j} - \sum_{i \in \to\textcircled{j}} \sigma^{st}_{i\to} \cdot p_{ij}\right)$ |
| | | $q$ | $\frac{\partial\mathcal{K}^{\mathrm{Surv}}_{\mathrm{eig}}(L)}{\partial q_j} = \frac{1}{\mathbf{v}^{\mathsf{T}}\mathbf{w}} \cdot \sum_{t=1}^{n} q_t \cdot \left(v_t \cdot w_j \cdot k^{\mathrm{Surv}}_{tj} + v_j \cdot w_t \cdot k^{\mathrm{Surv}}_{jt}\right)$ |
| | $k^{\mathrm{Surv}}$ | $c$ | $\frac{\partial\mathcal{K}^{\mathrm{Surv}}_{\mathrm{eig}}(L)}{\partial c_j} = -\frac{\theta}{\mathbf{v}^{\mathsf{T}}\mathbf{w}} \cdot \sum_{s,t=1}^{n} v_s \cdot w_t \cdot q_s \cdot q_t \cdot k^{\mathrm{Surv}}_{st} \cdot \overline{n}^{st}_{\to j}$ |
| | | $a$ | $\frac{\partial\mathcal{K}^{\mathrm{Surv}}_{\mathrm{eig}}(L)}{\partial a_j} = \frac{1}{a_j \cdot \mathbf{v}^{\mathsf{T}}\mathbf{w}} \cdot \sum_{s,t=1}^{n} v_s \cdot w_t \cdot q_s \cdot q_t \cdot k^{\mathrm{Surv}}_{st} \cdot \left(\overline{n}^{st}_{\to j} - \sum_{i \in \to\textcircled{j}} \overline{n}^{st}_{i\to} \cdot p_{ij}\right)$ |
| eigen | | $a = e^{-c}$ | $\frac{\mathrm{d}\mathcal{K}^{\mathrm{Surv}}_{\mathrm{eig}}(L)}{\mathrm{d}c_j} = \frac{1}{\mathbf{v}^{\mathsf{T}}\mathbf{w}} \cdot \sum_{s,t=1}^{n} v_s \cdot w_t \cdot q_s \cdot q_t \cdot k^{\mathrm{Surv}}_{st} \cdot \left((1-\theta)\overline{n}^{st}_{\to j} - \sum_{i \in \to\textcircled{j}} \overline{n}^{st}_{i\to} \cdot p_{ij}\right)$ |
| | | $q$ | $\frac{\mathrm{d}\mathcal{K}^{\mathrm{EC}}_{\mathrm{eig}}(L)}{\mathrm{d}q_j} = \frac{1}{\mathbf{v}^{\mathsf{T}}\mathbf{w}} \cdot \sum_{t=1}^{n} q_t \cdot \left(v_t \cdot w_j \cdot k^{\mathrm{EC}}_{tj} + v_j \cdot w_t \cdot k^{\mathrm{EC}}_{jt}\right)$ |
| | $k^{\mathrm{EC}}$ | $c$ | $\frac{\partial\mathcal{K}^{\mathrm{EC}}_{\mathrm{eig}}(L)}{\partial c_j} = -\frac{\alpha}{\mathbf{v}^{\mathsf{T}}\mathbf{w}} \cdot \sum_{s,t=1}^{n} v_s \cdot w_t \cdot q_s \cdot q_t \cdot k^{\mathrm{EC}}_{st} \cdot \left(\overline{n}^{st}_{\to j} + \theta \cdot \sigma^{st}_{\to j}\right)$ |
| | | $a$ | $\frac{\partial\mathcal{K}^{\mathrm{EC}}_{\mathrm{eig}}(L)}{\partial a_j} = \frac{\alpha}{a_j \cdot \mathbf{v}^{\mathsf{T}}\mathbf{w}} \cdot \sum_{s,t=1}^{n} v_s \cdot w_t \cdot q_s \cdot q_t \cdot k^{\mathrm{EC}}_{st} \cdot \left(\sigma^{st}_{\to j} - \sum_{i \in \to\textcircled{j}} \sigma^{st}_{i\to} \cdot p_{ij}\right)$ |
| | | $a = e^{-c}$ | $\frac{\mathrm{d}\mathcal{K}^{\mathrm{EC}}_{\mathrm{eig}}(L)}{\mathrm{d}c_j} = -\frac{\alpha}{\mathbf{v}^{\mathsf{T}}\mathbf{w}} \cdot \sum_{s,t=1}^{n} v_s \cdot w_t \cdot q_s \cdot q_t \cdot k^{\mathrm{EC}}_{st} \cdot \left(\overline{n}^{st}_{\to j} + (\theta+1)\sigma^{st}_{\to j} - \sum_{i \in \to\textcircled{j}} \sigma^{st}_{i\to} \cdot p_{ij}\right)$ |

Table C.3: Sensitivity analysis expressions. The notation used here is reviewed in Table C.2. The partial derivatives (with $\partial$'s) are meant for use in cases where $a_j$ and $c_j$ (defined on nodes) do not depend upon each other, whereas the total derivatives (with d's) are applicable when the perturbation of a cost $c_j$ also causes a perturbation of the step probability according to $a_j = \exp(-c_j)$.

*Appendix C.2.1. Sensitivity of $\mathcal{K}_{\text{sum}}(L)$ to habitat degradation (i.e. change in $q$)*

The sensitivity of $\mathcal{K}_{\text{sum}}(L)$ to a change in the quality of a node $j$ can be considered using the derivative

$$\frac{\mathrm{d}\mathcal{K}_{\text{sum}}(L)}{\mathrm{d}q_j} = \sum_{s,t=1}^{n} \frac{\mathrm{d}(q_s q_t)}{\mathrm{d}q_j} \cdot k_{st}. \tag{C.8}$$

Note that the derivative does not affect the proximity $k_{st}$, as the proximities are defined independently of qualities, as stated earlier. Thus, the result here holds for any proximity measure $k_{st}$. The summand in Equation (C.8) is nonzero only if either $s = j$ or $t = j$ or both $s = t = j$. Accordingly, Equation (C.8) can be written as

$$\frac{\mathrm{d}\mathcal{K}_{\text{sum}}(L)}{\mathrm{d}q_j} = \sum_{\substack{s=1\\ s\neq j}}^{n} \frac{\mathrm{d}(q_s q_j)}{\mathrm{d}q_j} \cdot k_{sj} + \sum_{\substack{t=1\\ t\neq j}}^{n} \frac{\mathrm{d}(q_j q_t)}{\mathrm{d}q_j} \cdot k_{jt} + \frac{\mathrm{d}q_j^2}{\mathrm{d}q_j} \cdot k_{jj} \tag{C.9}$$

$$= \sum_{\substack{s=1\\ s\neq j}}^{n} q_s \cdot k_{sj} + \sum_{\substack{t=1\\ t\neq j}}^{n} q_t \cdot k_{jt} + 2q_j \cdot k_{jj} \tag{C.10}$$

$$= \sum_{s=1}^{n} q_s \cdot k_{sj} + \sum_{t=1}^{n} q_t \cdot k_{jt} \tag{C.11}$$

$$= \sum_{t=1}^{n} q_t (k_{tj} + k_{jt}). \tag{C.12}$$

This shows that the quality-sensitivity of the summation-based landscape functionality corresponds to a type of *closeness centrality* measure on a graph; more exactly, it generalizes the *harmonic centrality* (Boldi and Vigna, 2014) of node $j$, which is obtained when the proximities are defined as $k_{st} = 1/d_{st}^{\text{LC}}$ and qualities are set to unity. Thus, the sensitivity of the landscape functionality to a perturbation of the quality of cell $j$ is simply the sum of its contribution as both a source and a target for all cells. Therefore, the computation of the sensitivity of $\mathcal{K}_{\text{sum}}(L)$ w.r.t. $q_j$ is fairly straightforward, as it is the sum of the qualities of all pixels ($q_t$) on the landscape weighted by the summed proximity to and from those pixels (respectively $k_{jt}$ and $k_{tj}$) – obviously , with the proximity metric matching the one for the landscape functionality (e.g. $k_{st}^{\text{Surv}}$ or $k_{st}^{\text{EC}}$, but also $k_{st}^{\text{Eucl}}$ or $k_{st}^{\text{LC}}$).

*Appendix C.2.2. Sensitivity of $\mathcal{K}_{\text{sum}}(L)$ to habitat fragmentation through movement costs (i.e. change in c)*

The sensitivity of $\mathcal{K}_{\text{sum}}(L)$ to perturbation of a cost $c_j$, assuming that the perturbation of $c_j$ does not affect $a_j$ (nor $q_j$), can be quantified with the partial derivative

$$\frac{\partial\mathcal{K}_{\text{sum}}(L)}{\partial c_j} = \sum_{s,t=1}^{n} q_s q_t \cdot \frac{\partial k_{st}}{\partial c_j}. \tag{C.13}$$

Note that by inserting $1 = \big(\delta_{sj} + (1-\delta_{sj})\big)\big(\delta_{tj} + (1-\delta_{tj})\big)$ as a factor within the summation, it can easily be shown that this expression can be decomposed into a sum of four components:

$$\begin{aligned}\frac{\partial \mathcal{K}_{\text{sum}}(L)}{\partial c_j} &= q_j^2 \cdot \frac{\partial k_{jj}}{\partial c_j} + q_j \cdot \sum_{\substack{t=1\\t\neq j}}^{n} q_t \cdot \frac{\partial k_{jt}}{\partial c_j} + q_j \cdot \sum_{\substack{s=1\\s\neq j}}^{n} q_s \cdot \frac{\partial k_{sj}}{\partial c_j} + \sum_{\substack{s,t=1\\s\neq j \wedge t\neq j}}^{n} q_s q_t \cdot \frac{\partial k_{st}}{\partial c_j}\\ &= q_j \cdot \sum_{\substack{t=1\\t\neq j}}^{n} q_t \cdot \frac{\partial k_{jt}}{\partial c_j} + q_j \cdot \sum_{\substack{s=1\\s\neq j}}^{n} q_s \cdot \frac{\partial k_{sj}}{\partial c_j} + \sum_{\substack{s,t=1\\s\neq j \wedge t\neq j}}^{n} q_s q_t \cdot \frac{\partial k_{st}}{\partial c_j},\end{aligned} \quad \text{(C.14)}$$

where $\partial k_{jj}/\partial c_j = 0$ because the distance of a node to itself is zero so that the elements on the diagonal of the proximity matrix are constant, and of course the derivative of proximity $k$ w.r.t. the step costs ($c_j$) depends on the type of proximity used (i.e. $k^{\text{Surv}}$ or $k^{\text{EC}}$).

*Survival probability proximity.* Let us first consider the sensitivity of the survival probability proximity this is called the power mean proximity., defined as $k_{st}^{\text{Surv}} = \mathcal{Z}_{st}$. It was shown by Kivimäki et al. (2024), Equation (53), that for any edge cost $c_{ij}$,

$$\frac{\partial \mathcal{Z}_{st}}{\partial c_{ij}} = -\theta \mathcal{Z}_{st} \overline{n}_{ij}^{st}, \quad \text{(C.15)}$$

where

$$\overline{n}_{ij}^{st} = \sum_{\wp \in \mathcal{P}_{st}} \mathrm{P}^{\text{RSP}}(\wp) n_{ij}(\wp) \quad \text{(C.16)}$$

is the expected number of traversals over edge $(i,j)$ when moving from $s$ to $t$ according to the RSP probability distribution (for details on the computation of $\overline{n}_{ij}^{st}$, see Kivimäki et al., 2016, 2024, Equations (9)-(10) in the later reference).

As edge costs in this work are defined by costs on the nodes, the sensitivity of the survival proximity w.r.t. the cost on node $j$ is given by

$$\frac{\partial k_{st}^{\text{Surv}}}{\partial c_j} = \frac{\partial \mathcal{Z}_{st}}{\partial c_j} = \sum_{i \in \to \textcircled{j}} \frac{\partial \mathcal{Z}_{st}}{\partial c_{ij}} = -\theta k_{st}^{\text{Surv}} \sum_{i \in \to \textcircled{j}} \overline{n}_{ij}^{st}, \quad \text{(C.17)}$$

where the sum is taken over the set of predecessor nodes of node $j$ (denoted by '$\to\textcircled{j}$'). The sum appearing in the last expression of Equation (C.17) contains the expected number of visits to node $j$ based on the incoming flow, which we denote, accordingly, as

$$\overline{n}_{\to j}^{st} = \sum_{i \in \to \textcircled{j}} \overline{n}_{ij}^{st} \quad \text{(C.18)}$$

for brevity. The cost-sensitivity of the survival probability can thus be written as

$$\frac{\partial k_{st}^{\text{Surv}}}{\partial c_j} = -\theta k_{st}^{\text{Surv}} \overline{n}_{\to j}^{st}. \quad \text{(C.19)}$$

Note that, normally, the number of visits to a node is defined in the RSP framework (see, e.g., Kivimäki et al., 2016) considering the outgoing flow, i.e. the sum over successor nodes,

$$\overline{n}_{j \to}^{st} = \sum_{k \in \textcircled{j} \to} \overline{n}_{jk}^{st}. \quad \text{(C.20)}$$

The main reason for opting for the outgoing flow over the incoming one here is simply notational, as the outgoing flow has a simpler computational expression. However, the two visit rates are equal for all nodes except for the source $s$ and target $t$, as they must satisfy a variant of Kirchhoff's law reflecting the conservation of flows,

$$\overline{n}^{st}_{\to j} + \delta_{sj} = \overline{n}^{st}_{j\to} + \delta_{jt}, \tag{C.21}$$

for all $j = 1, \ldots, n$, where the Kronecker delta is $\delta_{ab} = 1$, if $a = b$ and 0 otherwise.

Thus, the cost sensitivity of landscape functionality based on survival proximity for a node is determined by the betweenness of this node (expected number of visits) weighted by the qualities and proximity of source and target nodes. In Van Moorter et al. (2023a), we termed this $k$-weighted betweenness, which is similar in idea to the 'generalized betweenness' introduced by Bodin and Saura (2010).

*Expected cost proximity.* Let us then consider the sensitivity of the landscape functionality based on the expected cost proximity, $k^{\mathrm{EC}}_{st} = \exp(-\bar{c}_{st})$. For this, as shown by Kivimäki et al. (2024) (Equation (63)), the sensitivity of $k^{\mathrm{EC}}_{st}$ w.r.t. a perturbation of an edge cost $c_{ij}$ is given by the partial derivative

$$\frac{\partial \bar{c}_{st}}{\partial c_{ij}} = \overline{n}^{st}_{ij} + \theta \sigma^{st}_{ij}, \tag{C.22}$$

where $\sigma^{st}_{ij}$ is the negative covariance between the cost of a path, $c(\wp)$, and the number of times that the path traverses edge $(i, j)$. Formally (Equations (64-66) of the same paper),

$$\sigma^{st}_{ij} = -\mathrm{cov}(c(\wp), n_{ij}(\wp)) = \bar{c}_{st}\overline{n}^{st}_{ij} - \overline{cn}^{st}_{ij}, \tag{C.23}$$

where $\overline{cn}^{st}_{ij} = \sum_{\wp \in \mathcal{P}_{st}} \mathrm{P}^{\mathrm{RSP}}(\wp) n_{ij}(\wp) c(\wp)$ is the expectation of the product $n_{ij}(\wp)c(\wp)$ over the paths distribution $\mathrm{P}^{\mathrm{RSP}}$. The negative covariance for a fixed $s$-$t$-pair, $\sigma^{st}_{ij}$ is positive for an edge $(i, j)$ if $s$-$t$-paths that are likely to traverse edge $(i, j)$ also likely have a low cost, and, on the other hand, if high-cost $s$-$t$-paths rarely visit edge $(i, j)$. On the contrary, $\sigma^{st}_{ij}$ is negative if paths that are likely to visit edge $(i, j)$ also likely have high cost, whereas low-cost paths do not tend to visit $(i, j)$. This happens when edge $(i, j)$ lies 'far away' from the low-cost paths between $s$ and $t$. Thus, $\sigma^{st}_{ij}$ can be considered as a signed betweenness measure, as a positive value of $\sigma^{st}_{ij}$ indicates that edge $(i, j)$ lies more likely on low-cost paths between $s$ and $t$ than on high-cost paths. It is not surprising that the sensitivivity of the expected cost depends upon $\sigma^{st}_{ij}$, as a positive value for $\sigma^{st}_{ij}$ also implies stronger sensitivity of the expected cost $\bar{c}_{st}$ to perturbations of the edge cost $c_{ij}$.

Based on Equation (C.22), the sensitivity of $\bar{c}_{st}$ w.r.t. a perturbation of node cost $c_j$ is

$$\frac{\partial \bar{c}_{st}}{\partial c_j} = \sum_{i \in \to\textcircled{j}} \frac{\partial \bar{c}_{st}}{\partial c_{ij}} = \sum_{i \in \to\textcircled{j}} (\overline{n}^{st}_{ij} + \theta \sigma^{st}_{ij}) = \overline{n}^{st}_{\to j} + \theta \sigma^{st}_{\to j}, \tag{C.24}$$

where we have defined $\sigma^{st}_{\to j} = \sum_{i \in \to\textcircled{j}} \sigma^{st}_{ij}$ (similarly to the definition of $\overline{n}^{st}_{\to j}$ in Equation (C.18)). The term $\sigma^{st}_{\to j}$ in fact quantifies the negative covariance between the quantities $n_{\to j}(\wp)$ (expected number of visits to $j$ along path $\wp$) and $c(\wp)$ w.r.t. the RSP distribution, as we have, based on the definition in Equation (C.23), that

$$\sigma^{st}_{\to j} = \sum_{i \in \to \textcircled{j}} \sigma^{st}_{ij} = \sum_{i \in \to \textcircled{j}} (\bar{c}_{st}\bar{n}^{st}_{ij} - \overline{cn}^{st}_{ij}) \tag{C.25}$$

$$= \sum_{i \in \to \textcircled{j}} \left[ \left( \sum_{\wp \in \mathcal{P}_{st}} \mathrm{P}^{\mathrm{RSP}}(\wp) c(\wp) \right) \left( \sum_{\wp \in \mathcal{P}_{st}} \mathrm{P}^{\mathrm{RSP}}(\wp) n_{ij}(\wp) \right) - \sum_{\wp \in \mathcal{P}_{st}} \mathrm{P}^{\mathrm{RSP}}(\wp) c(\wp) n_{ij}(\wp) \right] \tag{C.26}$$

$$= \sum_{\wp \in \mathcal{P}_{st}} \mathrm{P}^{\mathrm{RSP}}(\wp) c(\wp) \sum_{\wp \in \mathcal{P}_{st}} \mathrm{P}^{\mathrm{RSP}}(\wp) n_{\to j}(\wp) - \sum_{\wp \in \mathcal{P}_{st}} \mathrm{P}^{\mathrm{RSP}}(\wp) c(\wp) n_{\to j}(\wp) \tag{C.27}$$

$$= -\mathrm{cov}(c(\wp), n_{\to j}(\wp)). \tag{C.28}$$

Thus, using Equation (C.24) and the chain rule, the sensitivity of the expected cost proximity, $k^{\mathrm{EC}}_{st}$, w.r.t. $c_j$ becomes

$$\frac{\partial k^{\mathrm{EC}}_{st}}{\partial c_j} = \frac{\partial \exp(-\alpha \bar{c}_{st})}{\partial \bar{c}_{st}} \cdot \frac{\partial \bar{c}_{st}}{\partial c_j} = -\alpha \cdot \exp(-\alpha \bar{c}_{st}) \cdot \frac{\partial \bar{c}_{st}}{\partial c_j} = -\alpha k^{\mathrm{EC}}_{st} \left( \bar{n}^{st}_{\to j} + \theta \sigma^{st}_{\to j} \right). \tag{C.29}$$

Whereas the sensitivity of the survival-based landscape functionality w.r.t. movement cost is directly dependent upon the betweenness (see Equation (C.19)), the sensitivity of the EC-based landscape functionality depends upon a novel 'betweenness' metric: the negative covariance between the cost and the betweenness. Another way to understand this negative covariance is in terms of a 'detour cost'. Indeed, Kivimäki et al. (2024) showed that the negative covariance can be written as the product of the betweenness and a detour cost (Equations (75)-(77) in Kivimäki et al., 2024). As explained in that paper, this detour cost can be negative, which corresponds to a detour benefit. The detour cost associated to a pixel $i$ is the difference in expected cost of moving directly from $s$ to $t$ and the cumulative expected cost of moving first from $s$ to $i$ and then from $i$ to $t$. Thus, $\sigma^{st}_i$ can be considered as a signed betweenness measure, as a positive value of $\sigma^{st}_i$ indicates that node $i$ lies more likely on low-cost paths between $s$ and $t$ than on high-cost paths. The absolute value of $\sigma^{st}_i$ for a node $i$ is low, when it has low betweenness (i.e. few paths are expected to pass) or when the cost of the passing paths is not particularly high or low. It is not surprising that the sensitivivity of the expected cost depends upon $\sigma^{st}_i$, as a high absolute value for $\sigma^{st}_i$ also implies stronger sensitivity of the expected cost $\bar{c}_{st}$ to perturbations of the edge cost $c_{ij}$.

*Appendix C.2.3. Sensitivity of $\mathcal{K}_{\mathrm{sum}}(L)$ to habitat fragmentation through behavioral avoidance (i.e. change in $a$)*

Similarly to the decomposition of Equation (C.14), we can decompose the sensitivity of $\mathcal{K}_{\mathrm{sum}}(L)$ w.r.t. $a_j$ as:

$$\frac{\partial \mathcal{K}_{\mathrm{sum}}(L)}{\partial a_j} = q_j^2 \cdot \frac{\partial k_{jj}}{\partial a_j} + q_j \cdot \sum_{\substack{t=1 \\ t \neq j}}^{n} q_t \cdot \frac{\partial k_{jt}}{\partial a_j} + q_j \cdot \sum_{\substack{s=1 \\ s \neq j}}^{n} q_s \cdot \frac{\partial k_{sj}}{\partial a_j} + \sum_{\substack{s,t=1 \\ s \neq j \wedge t \neq j}}^{n} q_s q_t \cdot \frac{\partial k_{st}}{\partial a_j}$$

$$= q_j \cdot \sum_{\substack{t=1 \\ t \neq j}}^{n} q_t \cdot \frac{\partial k_{jt}}{\partial a_j} + q_j \cdot \sum_{\substack{s=1 \\ s \neq j}}^{n} q_s \cdot \frac{\partial k_{sj}}{\partial a_j} + \sum_{\substack{s,t=1 \\ s \neq j \wedge t \neq j}}^{n} q_s q_t \cdot \frac{\partial k_{st}}{\partial a_j}, \tag{C.30}$$

which ultimately depends on the type of proximity used (i.e. $k^{\mathrm{Surv}}$ or $k^{\mathrm{EC}}$).

*Survival probability proximity.* Let us then consider the sensitivity of the survival proximity, $k_{st}^{\text{Surv}} = \mathcal{Z}_{st}$, to perturbations of a step probability $a_j$ of node $j$. We know from Kivimäki et al. (2024) (Equation (54)) that the partial derivative of the survival proximity w.r.t. a step probability of an edge $(i, j)$ is

$$\frac{\partial k_{st}^{\text{Surv}}}{\partial a_{ij}} = \frac{\partial \mathcal{Z}_{st}}{\partial a_{ij}} = \mathcal{Z}_{st}\left(\frac{\overline{n}_{ij}^{st}}{a_{ij}} - \frac{\overline{n}_{i\rightarrow}^{st}}{a_{i\rightarrow}}\right) = k_{st}^{\text{Surv}}\left(\frac{\overline{n}_{ij}^{st}}{a_{ij}} - \frac{\overline{n}_{i\rightarrow}^{st}}{a_{i\rightarrow}}\right), \tag{C.31}$$

where $a_{i\rightarrow} = \sum_{k\in\textcircled{i}\rightarrow} a_{ik}$ is simply the out-degree of node $i$. As we assume that $a_{ij} = a_j$ for all $i \in \rightarrow\!\textcircled{j}$, the sensitivity w.r.t. the node step probability $a_j$ is given by the sum of the edgewise partial derivatives over incoming edges to node $j$:

$$\frac{\partial k_{st}^{\text{Surv}}}{\partial a_j} = \sum_{i\in\rightarrow\textcircled{j}} \frac{\partial \mathcal{Z}_{st}}{\partial a_{ij}} \cdot \frac{\partial a_{ij}}{\partial a_j} = \sum_{i\in\rightarrow\textcircled{j}} \frac{\partial \mathcal{Z}_{st}}{\partial a_{ij}} = k_{st}^{\text{Surv}}\left(\sum_{i\in\rightarrow\textcircled{j}}\left(\frac{\overline{n}_{ij}^{st}}{a_{ij}} - \frac{\overline{n}_{i\rightarrow}^{st}}{a_{i\rightarrow}}\right)\right) \tag{C.32}$$

$$= k_{st}^{\text{Surv}}\left(\frac{\overline{n}_{\rightarrow j}^{st}}{a_j} - \sum_{i\in\rightarrow\textcircled{j}} \frac{\overline{n}_{i\rightarrow}^{st}}{a_{i\rightarrow}}\right) = \frac{k_{st}^{\text{Surv}}}{a_j}\left(\overline{n}_{\rightarrow j}^{st} - \sum_{i\in\rightarrow\textcircled{j}} \overline{n}_{i\rightarrow}^{st} p_{ij}\right), \tag{C.33}$$

and we recall from Table C.2 the definition of the unbiased random walk transition probabilities as $p_{ij} = a_j/a_{i\rightarrow}$.

Thus, for the sensitivity w.r.t. the step probability we find a similar dependence on the betweenness of a cell (i.e. $k$-weighted betweenness) as w.r.t. the step cost; however, the betweenness comes in the term $(\overline{n}_{\rightarrow j}^{st} - \sum_{i\in\rightarrow\textcircled{j}} \overline{n}_{i\rightarrow}^{st} \cdot p_{ij})/a_j$ provided by the RSP framework. This term can also be written as $\sum_{i\in\rightarrow\textcircled{j}} \overline{n}_{i\rightarrow}^{st}\big(p_{ij}^{(\text{t})} - p_{ij}\big)/a_j$, where $p_{ij}^{(\text{t})} = \overline{n}_{ij}^{st}/\overline{n}_{i\rightarrow}^{st}$ is the *biased* (towards the target node $t$) transition probability provided by the RSP Kivimäki et al. (2024). Thus, the sensitivity corresponds to the betweenness weighted with the change in transition probability caused by considering the long-term movement cost instead of moving according to only local preferences. In other words, it quantifies the difference in how often paths enter node $j$ from a predecessor node $i \in \rightarrow\!\textcircled{j}$ when comparing target-biased movement with unbiased random movement. This value is positive when the bias causes more visits to $j$ from predecessors, and negative when paths that visit a predecessor node tend to drift away from node $j$. Hence, all else being equal, the sensitivity will be higher to perturbations in nodes that have a larger difference between the biased and unbiased transition probabilities, especially when the movement is more optimal ($\theta \rightarrow \infty$). Thus, nodes that are used more when global information is being used (rather than moving randomly) will have a large positive sensitivity, whereas nodes that are used less when global information is being used have a large negative sensitivity.

*Expected cost proximity.* For the sensitivity of the expected cost proximity $k_{st}^{\text{EC}}$, we again use a result by Kivimäki et al. (2024), Equation (84), according to which the partial derivative of the expected cost $\bar{c}_{st}$ w.r.t. a step probability of an edge $(i, j)$ is

$$\frac{\partial \bar{c}_{st}}{\partial a_{ij}} = \frac{\sigma_{i\rightarrow}^{st}}{a_{i\rightarrow}} - \frac{\sigma_{ij}^{st}}{a_{ij}}, \tag{C.34}$$

and we recall that $\sigma_{i\rightarrow}^{st} = \sum_{k\in\textcircled{i}\rightarrow} \sigma_{ik}^{st}$. Again, as we assume that $a_{ij} = a_j$ for all $i \in \rightarrow\!\textcircled{j}$, the sensitivity w.r.t. the step probability of node $j$ is

$$\frac{\partial \bar{c}_{st}}{\partial a_j} = \sum_{i\in\rightarrow\textcircled{j}}\left(\frac{\sigma_{i\rightarrow}^{st}}{a_{i\rightarrow}} - \frac{\sigma_{ij}^{st}}{a_j}\right) = \frac{1}{a_j}\left(\sum_{i\in\rightarrow\textcircled{j}} \sigma_{i\rightarrow}^{st} p_{ij} - \sigma_{\rightarrow j}^{st}\right). \tag{C.35}$$

Thus, the sensitivity of the Expected cost proximity, $k_{st}^{\mathrm{EC}} = \exp(-\alpha \bar{c}_{st})$, w.r.t. a perturbation of $a_j$, is obtained using the chain rule, as in Equation (C.29):

$$\frac{\partial k_{st}^{\mathrm{EC}}}{\partial a_j} = -\alpha k_{st}^{\mathrm{EC}} \frac{\partial \bar{c}_{st}}{\partial a_j} = -\frac{\alpha k_{st}^{\mathrm{EC}}}{a_j} \left( \sum_{i \in \rightarrow \textcircled{j}} \sigma_{i\rightarrow}^{st} p_{ij} - \sigma_{\rightarrow j}^{st} \right). \tag{C.36}$$

Notice the similarity with Equation (C.33) where the node betweenness is replaced by the node covariance.

*Appendix C.2.4. Sensitivity of $\mathcal{K}_{\mathrm{sum}}(L)$ to habitat fragmentation through both movement costs and an adaptive behavioral response to it (i.e. change in $c$ and $a$)*

Many species may respond behaviorally to landscape perturbations affecting their cost of movement, hence, a perturbation of the cost of movement may not be independent of the probability of movement. Especially, when costs and step probabilities are considered to have a functional relationship, then it is reasonable to also consider that the perturbation of one will affect the other. In such a case, the sensitivity w.r.t. a perturbation should be considered using the total derivative, instead of the partial derivatives considered in the preceding sections. Here, we focus on a model, where the step probabilities are defined from costs as $a_j = f(c_j)$, for all $j$, where $f$ is some differentiable function, and that a perturbation of a cost $c_j$ will cause a change in step probability $a_j$ according to the function $f$. Then, the sensitivity of a proximity $k_{st}$ to a perturbation of a cost $c_j$ is given by the total derivative

$$\frac{\mathrm{d}k_{st}}{\mathrm{d}c_j} = \frac{\partial k_{st}}{\partial c_j} + \frac{\partial k_{st}}{\partial a_j} \cdot \frac{\mathrm{d}a_j}{\mathrm{d}c_j}, \tag{C.37}$$

where the latter form is obtained by applying the chain rule of the total derivative, and where $\mathrm{d}a_j/\mathrm{d}c_j = \mathrm{d}f(c_j)/\mathrm{d}c_j = f'$ is the derivative of function $f$ that defines step probabilities from costs. We refer to such sensitivity measures, where perturbations of costs and step probabilities are inter-dependent, as *total sensitivity*.

Moreover, let us specifically consider here that the step probability $a_j$ is defined from movement cost $c_j$ as $a_j = \exp(-c_j)$. The derivative of $a_j$ is then $\mathrm{d}a_j/\mathrm{d}c_j = -a_j$, and the sensitivity of $k_{st}$ to a perturbation of $c_j$ from Equation (C.37) becomes

$$\frac{\mathrm{d}k_{st}}{\mathrm{d}c_j} = \frac{\partial k_{st}}{\partial c_j} - \frac{\partial k_{st}}{\partial a_j} \cdot a_j. \tag{C.38}$$

*Survival probability proximity.* With these assumptions, the total sensitivity of the survival probability $k_{st}^{\mathrm{Surv}} = \mathcal{Z}_{st}$ to a perturbation of a cost $c_j$ can be expressed by inserting Equations (C.19) and (C.33) into Equation (C.38), which gives

$$\frac{\mathrm{d}k_{st}^{\mathrm{Surv}}}{\mathrm{d}c_j} = \frac{\partial \mathcal{Z}_{st}}{\partial c_j} - \frac{\partial \mathcal{Z}_{st}}{\partial a_j} \cdot a_j \tag{C.39}$$

$$= -\theta k_{st}^{\mathrm{Surv}} \overline{n}_{\rightarrow j}^{st} - \frac{k_{st}^{\mathrm{Surv}}}{a_j} \left( \overline{n}_{\rightarrow j}^{st} - \sum_{i \in \rightarrow \textcircled{j}} \overline{n}_{i\rightarrow}^{st} p_{ij} \right) \cdot a_j \tag{C.40}$$

$$= -k_{st}^{\mathrm{Surv}} \left( (\theta + 1) \overline{n}_{\rightarrow j}^{st} - \sum_{i \in \rightarrow \textcircled{j}} \overline{n}_{i\rightarrow}^{st} p_{ij} \right), \tag{C.41}$$

where, as before, the unbiased random walk transition probability of edge $(i, j)$ results from $p_{ij} = a_j / a_{i\rightarrow}$.

*Expected cost proximity.* The total sensitivity of $k_{st}^{\mathrm{EC}}$ can be derived similarly, by inserting Equations (C.29) and (C.36) into the expression of the total derivative of Equation (C.38):

$$\frac{\mathrm{d}k_{st}^{\mathrm{EC}}}{\mathrm{d}c_j} = -\alpha k_{st}^{\mathrm{EC}}\left(\overline{n}_{\to j}^{st} + \theta\sigma_{\to j}^{st}\right) + \frac{\alpha k_{st}^{\mathrm{EC}}}{a_j}\left(\sum_{i\in\to(j)} \sigma_{i\to}^{st} p_{ij} - \sigma_{\to j}^{st}\right) \cdot a_j \tag{C.42}$$

$$= -\alpha k_{st}^{\mathrm{EC}}\left(\overline{n}_{\to j}^{st} + (\theta+1)\sigma_{\to j}^{st} - \sum_{i\in\to(j)} \sigma_{i\to}^{st} p_{ij}\right). \tag{C.43}$$

*Appendix C.2.5. Sensitivity of $\mathcal{K}_{\mathrm{sum}}(L)$ to habitat fragmentation, when movement cost is derived from observed behavior (i.e. change in $c$ and $a$)*

While in the previous Section Appendix C.2.4 we considered an adaptive behavioral response to a changing movement cost ($c_j$), here we consider the case where cost is considered a function of behavior ($a_j$) for pragmatic reasons (see Van Moorter et al., 2021, for a discussion). Thus, similar as before, if one defines costs from step probabilities with a function $f$ as $c_j = f(a_j)$, then the sensitivity of $k_{st}$ to a perturbation of $a_j$ is given by the total derivative

$$\frac{\mathrm{d}k_{st}}{\mathrm{d}a_j} = \frac{\partial k_{st}}{\partial a_j} + \frac{\partial k_{st}}{\partial c_j} \cdot \frac{\mathrm{d}c_j}{\mathrm{d}a_j}. \tag{C.44}$$

When $c_j = -\log a_j$ and, accordingly, $\mathrm{d}c_j/\mathrm{d}a_j = -1/a_j$, the sensitivity of $k_{st}$ to a perturbation of $a_j$, from last Equation (C.44), can be written further as

$$\frac{\mathrm{d}k_{st}}{\mathrm{d}a_j} = \frac{\partial k_{st}}{\partial a_j} - \frac{\partial k_{st}}{\partial c_j} \cdot \frac{1}{a_j}. \tag{C.45}$$

*Survival probability proximity.* With these assumptions, the total sensitivity of the survival probability $k_{st}^{\mathrm{Surv}} = \mathcal{Z}_{st}$ to a perturbation of a step probability $a_j$ can be expressed by inserting Equations (C.33) and (C.19) into Equation (C.45), which gives

$$\frac{\mathrm{d}k_{st}^{\mathrm{Surv}}}{\mathrm{d}a_j} = \frac{\partial \mathcal{Z}_{st}}{\partial a_j} - \frac{\partial \mathcal{Z}_{st}}{\partial c_j} \cdot \frac{1}{a_j} \tag{C.46}$$

$$= k_{st}^{\mathrm{Surv}}\left(\frac{\overline{n}_{\to j}^{st}}{a_j} - \sum_{i\in\to(j)} \frac{\overline{n}_{i\to}^{st}}{a_{i\to}}\right) + \theta k_{st}^{\mathrm{Surv}}\overline{n}_{\to j}^{st} \cdot \frac{1}{a_j} \tag{C.47}$$

$$= \frac{k_{st}^{\mathrm{Surv}}}{a_j}\left((\theta+1)\overline{n}_{\to j}^{st} - \sum_{i\in\to(j)} \overline{n}_{i\to}^{st} p_{ij}\right). \tag{C.48}$$

*Expected cost proximity.* Again, the total sensitivity of $k_{st}^{\mathrm{EC}}$ can be derived similarly by inserting Equations (C.36) and (C.29) into the expression of the total derivative of Equation (C.45):

$$\frac{\mathrm{d}k_{st}^{\mathrm{EC}}}{\mathrm{d}a_j} = -\frac{\alpha k_{st}^{\mathrm{EC}}}{a_j}\left(\sum_{i\in\to(j)} \sigma_{i\to}^{st} p_{ij} - \sigma_{\to j}^{st}\right) + \alpha k_{st}^{\mathrm{EC}}\left(\overline{n}_{\to j}^{st} + \theta\sigma_{\to j}^{st}\right) \cdot \frac{1}{a_j} \tag{C.49}$$

$$= \frac{\alpha k_{st}^{\mathrm{EC}}}{a_j}\left(\overline{n}_{\to j}^{st} + (\theta+1)\sigma_{\to j}^{st} - \sum_{i\in\to(j)} \sigma_{i\to}^{st} p_{ij}\right). \tag{C.50}$$

In fact, because $a_j = \exp(-c_j)$, these derivatives with respect to $a_j$ can easily be obtained heuristically from the previous ones (computed for $c_j$) by considering

$$\frac{\mathrm{d}k_{st}}{\mathrm{d}c_j} = \frac{\mathrm{d}k_{st}}{\mathrm{d}a_j} \cdot \frac{\mathrm{d}a_j}{\mathrm{d}c_j} = \frac{\mathrm{d}k_{st}}{\mathrm{d}a_j} \cdot (-a_j) = -a_j \frac{\mathrm{d}k_{st}}{\mathrm{d}a_j}.$$

Therefore,

$$\frac{\mathrm{d}k_{st}}{\mathrm{d}a_j} = -\frac{1}{a_j} \frac{\mathrm{d}k_{st}}{\mathrm{d}c_j},$$

which is easily verified for both $k_{st}^{\mathrm{EC}}$ and $k_{st}^{\mathrm{Surv}}$.

*Appendix C.2.6. Sensitivity of the Equivalent Connected Habitat (ECH)*

The sensitivity of the Equivalent Connected Habitat (ECH) is similar to the sensitivity of the summation-based landscape functionality, as:

$$ECH = \sqrt{\mathcal{K}_{\mathrm{sum}}(L)} \tag{C.51}$$

By simply applying the chain rule we obtain:

$$\frac{\mathrm{d}ECH}{\mathrm{d}x_i} = \frac{\mathrm{d}\sqrt{\mathcal{K}_{\mathrm{sum}}(L)}}{\mathrm{d}x_i} = \frac{1}{2\sqrt{\mathcal{K}_{\mathrm{sum}}(L)}} \cdot \frac{\mathrm{d}\mathcal{K}_{\mathrm{sum}}(L)}{\mathrm{d}x_i} \tag{C.52}$$

Hence, the sensitivity of the ECH to local perturbations in habitat quality or permeability is proportional to the sensitivity of the summation-based landscape functionality to these perturbations, which are described in Section Appendix C.2.

*Appendix C.3. Algorithms for the sensitivity computation of $\mathcal{K}_{\mathrm{sum}}^{\mathrm{EC}}(L)$, $\mathcal{K}_{\mathrm{sum}}^{\mathrm{Surv}}(L)$, $\mathcal{K}_{\mathrm{sum}}^{\mathrm{PM}}(L)$*

In this section, we present the algorithms for computing the different sensitivity metrics included in the `ConScape` package that are based on the derivatives of the landscape functionality metric with respect to the parameters on the nodes of the network involved in the RSP model. The algorithms are based on the methods derived by Kivimäki et al. (2024) and the present paper. Note that the algorithms implemented in the `ConScape` library are for the landmark approach (i.e. $n \times m$ source and target nodes, respectively; see main text), whereas for ease of presentation, the algorithms presented here are for the full case (i.e. $n \times n$ source and target nodes).

*Appendix C.3.1. Sensitivity computation of $\mathcal{K}_{\mathrm{sum}}^{\mathrm{EC}}(L)$*

The main Algorithm 1 computes the sensitivity of $\mathcal{K}_{\mathrm{sum}}^{\mathrm{EC}}$, as defined by Equation (C.4), i.e. the summation-based landscape connectivity index based on the RSP expected cost proximity, $k_{st}^{\mathrm{EC}}$. More precisely, the computation of $\partial_{c_{ij}}\mathcal{K}_{\mathrm{sum}}^{\mathrm{EC}}(L)$ and $\partial_{a_{ij}}\mathcal{K}_{\mathrm{sum}}^{\mathrm{EC}}(L)$ is detailed in this section. It is implemented in the `randomizedshortestpath` module of the `ConScape` package as the function `EC_sensitivity`. As noted above, the algorithms here present the general principles, the exact algorithms in the `ConScape` package use a landmark approach (see main text).

There are a few differences with the presentation here and the one in the main text. First, the main text only considers the scenario where edge costs and step probabilities are defined by node costs and node step probabilities by defining $c_{ij} = c_j$ and $a_{ij} = a_j$. Here, we, however consider, for generality, that costs and step probabilities are defined separately for all edges. Although considering node parameters makes the presentation in the main text a bit cleaner, it does not benefit the computation; the sensitivity must in both cases be computed separately for each edge, and node sensitivity can be then computed from the edge sensitivities e.g. by summing the values on the incoming edges of a node.

Also, although the main text defines the proximity as $k_{st}^{\mathrm{EC}} = \exp(-\alpha\bar{c}_{st})$ (Yen et al., 2008; Saerens et al., 2009; Kivimäki et al., 2014; Kivimäki, 2018), and the dependency between step probabilities and costs is only considered based on the transformation $a_{ij} = \exp(-c_{ij})$, here, Algorithm 1 is instead written for general transformations defining step probabilities from costs and proximities from the expected costs. For these, we assume that $k : \mathbb{R}_{\geq 0} \to \mathbb{R}_{\geq 0}$ and $a : \mathbb{R}_{\geq 0} \to \mathbb{R}_{>0}$ are decreasing differentiable functions, and consider the proximity as $k_{st}^{\mathrm{EC}} = k(\bar{c}_{st})$ and the step probabilities as $a_{ij} = a(c_{ij})$. For ease of notation, $k_{st}^{\mathrm{EC}}$ will be simply denoted as $k_{st}$ in this subsection.

The algorithm 1 uses the derivatives of both of the transformations $k$ and $a$. The derivatives are hard-coded for the transformations that are implemented in the `ConScape` package, but for custom transformations the derivatives must be computed separately and the matrices of values $a'_{ij} = \partial a_{ij}/\partial c_{ij}$ and $k'_{st} = \partial k(\bar{c}_{st})/\partial \bar{c}_{st}$ must be given as input to `EC_sensitivity`.

With the above assumptions and assuming that a perturbation of edge costs and step probabilities are independent of each other, the sensitivity of $\mathcal{K}_{\mathrm{sum}}^{\mathrm{EC}}(L)$ w.r.t. an edge cost, $c_{ij}$, can be written, based on Kivimäki et al. (2024), Equation (63), as the partial derivative (see Equation (C.22))

$$s_{ij}^{c} \triangleq \frac{\partial \mathcal{K}_{\mathrm{sum}}^{\mathrm{EC}}(L)}{\partial c_{ij}} = \sum_{s,t=1}^{n} q_s q_t \frac{\partial k(\bar{c}_{st})}{\partial c_{ij}} = \sum_{s,t=1}^{n} q_s q_t k'_{st} \frac{\partial \bar{c}_{st}}{\partial c_{ij}} \tag{C.53}$$

$$= \sum_{s,t=1}^{n} q_s q_t k'_{st} \big(\bar{n}_{ij}^{st} + \theta \sigma_{ij}^{st}\big), \tag{C.54}$$

where $\sigma_{ij}^{st}$ is the negative covariance between the functions $n_{ij}(\wp)$ and $c(\wp)$ considered based on drawing a path $\wp$ according to the RSP distribution $\mathrm{P}^{\mathrm{RSP}}$ (for more information on $\sigma_{ij}^{st}$, see the discussion following Equation (C.23) as well as Kivimäki et al., 2024).

In addition, the sensitivity of $\mathcal{K}_{\mathrm{sum}}^{\mathrm{EC}}(L)$ to a perturbation of $a_{ij}$ is given, based on Equation (C.34), as

$$s_{ij}^{a} \triangleq \frac{\partial \mathcal{K}_{\mathrm{sum}}^{\mathrm{EC}}(L)}{\partial a_{ij}} = \sum_{s,t=1}^{n} q_s q_t k'_{st} \left( \frac{\sigma_{i\to}^{st}}{a_{i\to}} - \frac{\sigma_{ij}^{st}}{a_{ij}} \right), \tag{C.55}$$

where, as before, $a_{i\to} = \sum_j a_{ij}$ is the out-degree of node $i$.

Equations (C.54) and (C.55) show that the computation of the sensitivity of $\mathcal{K}_{\mathrm{sum}}^{\mathrm{EC}}(L)$ to perturbations of $c_{ij}$ or $a_{ij}$ is similar to the computation of the $k$-weighted betweenness presented in Van Moorter et al. (2023a). For convenience, the computation of the $k$-weighted RSP edge betweenness from Van Moorter et al. (2023a) is presented again in Algorithm 3.

However, compared to the betweenness case, here we have the derivative of the RSP expected cost proximity, $k'_{st}$, in place of the proximity $k_{st}$. In addition, the computation here is performed for each edge separately. Algorithm 1, derived below, computes the sensitivity matrices $\mathbf{S}^c$ and $\mathbf{S}^a$ whose each element $i, j$, such that $(i, j) \in \mathcal{E}$, contains the sensitivities $\partial_{c_{ij}} \mathcal{K}_{\mathrm{sum}}^{\mathrm{EC}}(L)$ and $\partial_{a_{ij}} \mathcal{K}_{\mathrm{sum}}^{\mathrm{EC}}(L)$, respectively, and other elements are $0$.

When considering dependence between the step probabilities and edge costs, i.e. that the perturbation of one causes a perturbation of the other, then the sensitivity is best quantified by the total derivative (e.g. with respect to $c_{ij}$), which can be computed using the chain rule as (see Kivimäki et al., 2024, for details)

$$\frac{\mathrm{d}\mathcal{K}^{\mathrm{EC}}_{\mathrm{sum}}(L)}{\mathrm{d}c_{ij}} = \frac{\partial\mathcal{K}^{\mathrm{EC}}_{\mathrm{sum}}(L)}{\partial c_{ij}} + \frac{\partial\mathcal{K}^{\mathrm{EC}}_{\mathrm{sum}}(L)}{\partial a_{ij}} \cdot \frac{\mathrm{d}a_{ij}}{\mathrm{d}c_{ij}}. \tag{C.56}$$

The computation of the sensitivity expressed in Equations (C.54) and (C.55) requires the processing of a proximity-weighted overall negative covariance which needs to be computed,

$$\sigma_{ij}^{(k)} = \sum_{s=1}^{n}\sum_{t=1}^{n} k_{st}\sigma_{ij}^{st}. \tag{C.57}$$

Note that, in place of $k_{st}$, Equations (C.54), (C.55) actually contain the term $q_s q_t k'_{st}$. Nevertheless, these terms can be precomputed and the result can be defined as a new 'proximity' including $q_s$ and $q_t$, after which the computation of (the part involving the term $\sigma_{ij}^{st}$ of) Equations (C.54) and (C.55) indeed become equivalent to computing Equation (C.57). For simplicity, we will use the shortcut notation $k_{st}$ in place of $q_s q_t k'^{\mathrm{EC}}_{st}$ in the sequel, which is equivalent $m_{st}$.

Let us now derive a method for computing Equation (C.57) in an efficient way (resembling the computation of the betweenness values in Appendix C from Van Moorter et al. (2023a)), which is slightly involved. The idea is to obtain a matrix expression for this quantity, avoiding the use of a loop over all $s$-$t$ pairs, which would be prohibitive. This will also allow the use of sparse matrices in the implementation of the algorithms. We start by using the fact that the negative covariance $\sigma_{ij}^{st}$ can be expressed (see Equation (83) in Kivimäki et al. (2024)) as

$$\sigma_{ij}^{st} = \mathring{\sigma}_{ij}^{st} - \mathring{\sigma}_{ij}^{tt} = \mathring{n}_{ij}^{st}(\mathring{c}_{st} - \mathring{c}_{si}) - (\mathring{n}_{ij}^{st} - \mathring{n}_{ij}^{tt})(c_{ij} + \mathring{c}_{jt}) - \mathring{n}_{ij}^{tt}(\mathring{c}_{tt} - \mathring{c}_{ti}), \tag{C.58}$$

where $\mathring{n}_{ij}^{st}$, $\mathring{c}_{st}$, and $\mathring{\sigma}_{ij}^{st}$ are, respectively, the expected number of passages, the expected cost from $s$ to $t$, and the negative covariance computed from the RSP distribution over regular (non-hitting) paths. The matrix expression for computing the expected cost matrix $\mathring{\mathbf{C}}$ on all $s$-$t$ pairs can be found in Equation (21) of Kivimäki et al. (2024).

Using this together with Equation (11) in (Kivimäki et al., 2024) (coming initially from Kivimäki et al. (2016)) providing the expected number of passages through edge $(i, j)$ for both regular and hitting paths, which is recalled here for convenience,

$$\mathring{n}_{ij}^{st} = \frac{z_{si}w_{ij}z_{jt}}{z_{st}}(\text{ regular paths}), \quad \overline{n}_{ij}^{st} = \left(\frac{z_{si}}{z_{st}} - \frac{z_{ti}}{z_{tt}}\right) w_{ij} z_{jt} = \mathring{n}_{ij}^{st} - \mathring{n}_{ij}^{tt}(\text{ hitting paths}), \tag{C.59}$$

where $\mathbf{W}$ is a $n \times n$ weight matrix containing elements $[\mathbf{W}]_{ij} = w_{ij} = [\mathbf{W}]_{ij} = p_{ij}\exp(-\theta c_{ij})$ and $\mathbf{Z} = (\mathbf{I} - \mathbf{W})^{-1}$ is called the fundamental matrix depending on costs $c_{ij}$ and transition probabilities $p_{ij}$ of the natural random walk.

After rearranging a bit we obtain, for each element $\sigma_{ij}^{(k)}$ defined in Equation (C.57),

$$\sigma_{ij}^{(k)} = \sum_{s=1}^{n}\sum_{t=1}^{n} k_{st}\sigma_{ij}^{st} \tag{C.60}$$

$$= \sum_{s,t=1}^{n} k_{st}\Big(\mathring{n}_{ij}^{st}(\mathring{c}_{st} - \mathring{c}_{si}) - \overline{n}_{ij}^{st}(c_{ij} + \mathring{c}_{jt}) - \mathring{n}_{ij}^{tt}(\mathring{c}_{tt} - \mathring{c}_{ti})\Big) \tag{C.61}$$

$$\begin{aligned} &= \sum_{s,t=1}^{n} k_{st}\left(\frac{z_{si}}{z_{st}} w_{ij} z_{jt}(\mathring{c}_{st} - \mathring{c}_{si}) - \left(\frac{z_{si}}{z_{st}} - \frac{z_{ti}}{z_{tt}}\right) w_{ij} z_{jt}(c_{ij} + \mathring{c}_{jt})\right) \\ &\quad - \sum_{t=1}^{n} k_{\to t}\frac{z_{ti}}{z_{tt}} w_{ij} z_{jt}(\mathring{c}_{tt} - \mathring{c}_{ti}) \end{aligned} \tag{C.62}$$

$$\begin{aligned} &= w_{ij}\Bigg[\sum_{s,t=1}^{n} z_{jt}\frac{k_{ts}^{\mathsf{T}}\mathring{c}_{ts}^{\mathsf{T}}}{z_{ts}^{\mathsf{T}}} z_{si} - \sum_{s,t=1}^{n} z_{jt}\frac{k_{ts}^{\mathsf{T}}}{z_{ts}^{\mathsf{T}}}\mathring{c}_{si} z_{si} - c_{ij}\sum_{t=1}^{n} z_{jt}\left(\sum_{s=1}^{n}\frac{k_{ts}^{\mathsf{T}}}{z_{ts}^{\mathsf{T}}} z_{si} - \frac{k_{\to t}}{z_{tt}} z_{ti}\right) \\ &\quad - \sum_{t=1}^{n}\mathring{c}_{jt} z_{jt}\left(\sum_{s=1}^{n}\frac{k_{ts}^{\mathsf{T}}}{z_{ts}^{\mathsf{T}}} z_{si} - \frac{k_{\to t}}{z_{tt}} z_{ti}\right) - \sum_{t=1}^{n} z_{jt}\left(\frac{k_{\to t}\mathring{c}_{tt}}{z_{tt}} z_{ti} - \frac{k_{\to t}}{z_{tt}}\mathring{c}_{ti} z_{ti}\right)\Bigg] \end{aligned} \tag{C.63}$$

$$\begin{aligned} &= w_{ij}\Bigg[\sum_{s,t=1}^{n} z_{jt}\hat{k}_{ts}^{\mathsf{T}}\mathring{c}_{ts}^{\mathsf{T}} z_{si} - \sum_{s,t=1}^{n} z_{jt}\hat{k}_{ts}^{\mathsf{T}} y_{si} - c_{ij}\sum_{t=1}^{n} z_{jt}\left(\sum_{s=1}^{n}\hat{k}_{ts}^{\mathsf{T}} z_{si} - \hat{k}_{t}^{\Sigma} z_{ti}\right) \\ &\quad - \sum_{t=1}^{n} y_{jt}\left(\sum_{s=1}^{n}\hat{k}_{ts}^{\mathsf{T}} z_{si} - \hat{k}_{t}^{\Sigma} z_{ti}\right) - \sum_{t=1}^{n} z_{jt}\left(\hat{k}_{t}^{\Sigma}\mathring{c}_{tt} z_{ti} - \hat{k}_{t}^{\Sigma} y_{ti}\right)\Bigg], \end{aligned} \tag{C.64}$$

in which we have defined, for all $s$, $t$,

$$\hat{k}_{st} = k_{st}/z_{st}, \quad \text{i.e. the matrix} \quad \widehat{\mathbf{K}} = \mathbf{K} \div \mathbf{Z}, \tag{C.65}$$

with $\div$ the elementwise division,

$$\hat{k}_{t}^{\Sigma} = k_{\to t}/z_{tt}, \quad \text{i.e. the column vector} \quad \hat{\mathbf{k}}^{\Sigma} = (\mathbf{K}^{\mathsf{T}}\mathbf{e}) \div \mathbf{diag}(\mathbf{Z}), \tag{C.66}$$

and from Equation (20) in Kivimäki et al. (2024),

$$y_{st} = \mathring{c}_{st} z_{st} = \sum_{k,l=1}^{n} z_{sk} w_{kl} c_{kl} z_{lt}, \quad \text{i.e. the matrix} \quad \mathbf{Y} = \mathbf{Z}(\mathbf{C}\circ\mathbf{W})\mathbf{Z}. \tag{C.67}$$

Equation (C.64) can then be further rewritten as

$$\sigma_{ij}^{(k)} = w_{ij}\left[\sum_{t=1}^{n} z_{jt}x_{ti}^{(1)} - \sum_{t=1}^{n} z_{jt}x_{ti}^{(2)} - c_{ij}\sum_{t=1}^{n} z_{jt}x_{ti}^{(3)} - \sum_{t=1}^{n} y_{jt}x_{ti}^{(3)} - \sum_{t=1}^{n} z_{jt}x_{ti}^{(4)}\right] \tag{C.68}$$

by defining four helper matrices as

$$\mathbf{X}^{(1)} = (\widehat{\mathbf{K}}\circ\mathring{\mathbf{C}})^{\mathsf{T}}\mathbf{Z}, \tag{C.69}$$

$$\mathbf{X}^{(2)} = \widehat{\mathbf{K}}^{\mathsf{T}}\mathbf{Y}, \tag{C.70}$$

$$\mathbf{X}^{(3)} = \widehat{\mathbf{K}}^{\mathsf{T}}\mathbf{Z} - \mathbf{Diag}(\hat{\mathbf{k}}^{\Sigma})\mathbf{Z} = \Big(\widehat{\mathbf{K}}^{\mathsf{T}} - \mathbf{Diag}(\hat{\mathbf{k}}^{\Sigma})\Big)\mathbf{Z}, \tag{C.71}$$

$$\mathbf{X}^{(4)} = \mathbf{Diag}\Big(\hat{\mathbf{k}}^{\Sigma}\circ\mathbf{diag}(\mathring{\mathbf{C}})\Big)\mathbf{Z} - \mathbf{Diag}(\hat{\mathbf{k}}^{\Sigma})\mathbf{Y}. \tag{C.72}$$

Note that, again from Equation (11) in Kivimäki et al. (2024) (see also Equation (C.59)), the third term of the right side of this last Equation (C.68) (the one multiplied by $c_{ij}$ and involving $\mathbf{X}^{(3)}$, defined in Equation (C.71)) is proportional to $w_{ij}\sum_{t=1}^{n} z_{jt}x_{ti}^{(3)} = \sum_{s,t=1}^{n} k_{st}\bar{n}_{ij}^{st} = \mathrm{bet}_{ij}^{k-\mathrm{RSP}}$ (with $k_{st} = q_s q_t k_{st}^{\prime\mathrm{EC}}$ as discussed before), which is nothing else than the $i$, $j$ element of the $k$-weighted edge betweenness[2] matrix, $\mathbf{Bet}^{k-\mathrm{RSP}}$ introduced in the Appendix C of Van Moorter et al. (2023a), Equation (31). This can be easily seen from Equations (C.61) and (C.62).

Using these, Equation (C.68) can be rewritten in matrix form as

$$\sigma_{ij}^{(k)} = w_{ij}\mathbf{e}_j^{\mathsf{T}}\big(\mathbf{Z}\mathbf{X}^{(1)} - \mathbf{Z}\mathbf{X}^{(2)} - c_{ij}\mathbf{Z}\mathbf{X}^{(3)} - \mathbf{Y}\mathbf{X}^{(3)} - \mathbf{Z}\mathbf{X}^{(4)}\big)\mathbf{e}_i \tag{C.73}$$

$$= w_{ij}\mathbf{e}_j^{\mathsf{T}}\big(\mathbf{Z}\mathbf{X}^{(5)} - c_{ij}\mathbf{Z}\mathbf{X}^{(3)} - \mathbf{Y}\mathbf{X}^{(3)}\big)\mathbf{e}_i, \tag{C.74}$$

$$= w_{ij}\big(\mathbf{e}_j^{\mathsf{T}}\mathbf{Z}\mathbf{X}^{(5)}\mathbf{e}_i - c_{ij}\mathbf{e}_j^{\mathsf{T}}\mathbf{Z}\mathbf{X}^{(3)}\mathbf{e}_i - \mathbf{e}_j^{\mathsf{T}}\mathbf{Y}\mathbf{X}^{(3)}\mathbf{e}_i\big), \tag{C.75}$$

where $\mathbf{e}_i$ is a column indicator vector with 0's everywhere except at position $i$, containing a 1. After this last step, only $\mathbf{Y}$, $\mathbf{X}^{(3)}$ and $\mathbf{X}^{(5)}$ are used after having further simplified the expression by defining

$$\mathbf{X}^{(5)} = \mathbf{X}^{(1)} - \mathbf{X}^{(2)} - \mathbf{X}^{(4)}. \tag{C.76}$$

Finally, the elements $\sigma_{ij}^{(k)}$ are gathered in a new matrix called $\mathbf{\Sigma}^{k-\mathrm{RSP}}$. The corresponding pseudocode is depicted in Algorithm 1. In this code, the elements of the matrix are computed by performing a loop on the edges and using Equation (C.75). However, one could also use the following closed-form matrix solution without a loop, depending on the efficiency of the sparse matrix multiplication,

$$\mathbf{\Sigma}^{k-\mathrm{RSP}} = \mathbf{W} \circ \Big(\big(\mathbf{Z}\mathbf{X}^{(5)}\big)^{\mathsf{T}} - \mathbf{C} \circ \big(\mathbf{Z}\mathbf{X}^{(3)}\big)^{\mathsf{T}} - \big(\mathbf{Y}\mathbf{X}^{(3)}\big)^{\mathsf{T}}\Big). \tag{C.77}$$

*Appendix C.3.2. Sensitivity computation of $\mathcal{K}_{\mathrm{sum}}^{\mathrm{Surv}}(L)$*

The derivation of the sensitivity for the survival proximity (Equation (C.3) with $k_{st}^{\mathrm{Surv}} = \mathcal{Z}_{st}$) is much simpler than for the expected cost. Proceeding as for the expected cost and using Equation (C.15) provides, for the sensitivity w.r.t. edge cost,

$$s_{ij}^{c} \triangleq \frac{\partial \mathcal{K}_{\mathrm{sum}}^{\mathrm{Surv}}(L)}{\partial c_{ij}} = \frac{\partial \sum_{s,t=1}^{n} q_s q_t k_{st}^{\mathrm{Surv}}}{\partial c_{ij}} \tag{C.78}$$

$$= \sum_{s,t=1}^{n} q_s q_t \frac{\partial \mathcal{Z}_{st}}{\partial c_{ij}} = \sum_{s,t=1}^{n} q_s q_t (-\theta \bar{n}_{ij}^{st} \mathcal{Z}_{st}) \tag{C.79}$$

$$= -\theta \sum_{s,t=1}^{n} q_s q_t \mathcal{Z}_{st} \bar{n}_{ij}^{st} = -\theta\, \mathrm{bet}_{ij}^{k-\mathrm{RSP}}, \tag{C.80}$$

where, in the last line, $\mathrm{bet}_{ij}^{k-\mathrm{RSP}}$ is the already discussed $k$-weighted node betweenness (weighted by qualities $q_s$ and $q_t$, $k_{st}^{\mathrm{Surv}} = q_s \mathcal{Z}_{st} q_t$ here) described in the Postscript section, at the end of this paper (see Equation (C.110)), where an algorithm for computing it is proposed (Algorithm

[2] Also computed in another way in Algorithm 3 of the Postscript added to this paper – see Equation (C.110).

**Algorithm 1** Computation of the sensitivities of the summation-based landscape connectivity index based on the RSP expected cost proximity (Equation (C.4)) w.r.t. step probability (edge affinity) or cost of movement, $\mathbf{S}^a$ and $\mathbf{S}^c$, respectively (Equations (C.54) and (C.55)), as implemented in the `EC_sensitivity` function.

**Input:**

– A directed, strongly connected, graph $G$ containing $n$ nodes.
– The $n \times n$ non-negative affinity matrix $\mathbf{A}$ associated to $G$.
– The $n \times n$ non-negative cost matrix $\mathbf{C}$ associated to $G$.
– The inverse temperature parameter $\theta > 0$.
– The $n \times n$ non-negative diagonal matrices $\mathbf{Q}^s$ and $\mathbf{Q}^t$ containing the qualities of nodes $s$ and $t$, resp., on the diagonals.
– The distance scaling parameter $\alpha > 0$.

**Output:**

– The $n \times n$ sparse matrices $\mathbf{S}^a$ and $\mathbf{S}^c$ containing, for each edge $(i,j) \in \mathcal{E}$, the sensitivities $\partial_{c_{ij}} \mathcal{K}^{\mathrm{EC}}_{\mathrm{sum}}(L)$ and $\partial_{a_{ij}} \mathcal{K}^{\mathrm{EC}}_{\mathrm{sum}}(L)$, respectively, at the corresponding elements $(i,j)$, and 0 elsewhere, computed from Equations (C.54) and (C.55).

1. $\mathbf{D} \leftarrow \mathbf{Diag}(\mathbf{Ae})$ {the outdegree row-normalization diagonal matrix; $\mathbf{e}$ is a column vector of 1's; see Kivimäki et al. (2024), Eq. (1)}
2. $\mathbf{P}^{\mathrm{rw}} \leftarrow \mathbf{D}^{-1}\mathbf{A}$ {the reference, natural, random walk transition probabilities matrix; see Kivimäki et al. (2024), Eq. (1)}
3. $\mathbf{W} \leftarrow \mathbf{P}^{\mathrm{rw}} \circ \exp(-\theta \mathbf{C})$ {the $\mathbf{W}$ substochastic matrix involving elementwise exponential and multiplication $\circ$; see Kivimäki et al. (2024), Eq. (5)}
4. $\mathbf{Z} \leftarrow (\mathbf{I} - \mathbf{W})^{-1}$ {the fundamental matrix; see Kivimäki et al. (2024), Eq. (8)}
5. $\mathbf{Y} \leftarrow \mathbf{Z}(\mathbf{C} \circ \mathbf{W})\mathbf{Z}$ {see Eq. (C.67); will be used later}
6. $\mathring{\mathbf{C}} \leftarrow \mathbf{Y} \div \mathbf{Z}$ {elementwise division; computes the matrix containing expected costs for regular (non-hitting) paths; see Kivimäki et al. (2024), Eq. (21), or Kivimäki et al. (2014)}
7. $\mathbf{d}_{\mathbf{C}} \leftarrow \mathbf{diag}(\mathring{\mathbf{C}})$ {the column vector of diagonal values of $\mathring{\mathbf{C}}$; see Kivimäki et al. (2024), Eq. (23)}
8. $\overline{\mathbf{C}} \leftarrow \mathring{\mathbf{C}} - \mathbf{e}\mathbf{d}_{\mathbf{C}}^{\mathsf{T}}$ {the matrix of expected costs for hitting paths, with $\mathbf{e}$ a column vector full of 1's; see Kivimäki et al. (2024), Eq. (23)}
9. $\mathbf{K} \leftarrow \exp(-\alpha \overline{\mathbf{C}})$ {the proximity matrix as an elementwise exponential transformation of the weighted RSP expected cost; see the paragraph just before Eq. (C.36)}
10. $\mathbf{K}' \leftarrow -\alpha \mathbf{K}$ {the derivative of the proximity matrix w.r.t. the RSP expected costs; see Eq. (C.29)}
11. $\mathbf{K}' \leftarrow \mathbf{Q}^s \mathbf{K}' \mathbf{Q}^t$ {the $q$-weighted proximity matrix (optional; only if source-target quality weighting is desired); see Eq. (C.4)}
12. $\widehat{\mathbf{K}} \leftarrow \mathbf{K}' \div \mathbf{Z}$ {see Eq. (C.65)}
13. $\hat{\mathbf{k}}^{\Sigma} \leftarrow (\mathbf{K}'^{\mathsf{T}}\mathbf{e}) \div \mathbf{diag}(\mathbf{Z})$ {see Eq. (C.66)}
14. $\mathbf{X}^{(5)} \leftarrow \Big( (\widehat{\mathbf{K}} \circ \mathring{\mathbf{C}})^{\mathsf{T}} - \widehat{\mathbf{K}}^{\mathsf{T}} \mathbf{Z}(\mathbf{C} \circ \mathbf{W}) + \mathbf{Diag}(\hat{\mathbf{k}}^{\Sigma}) \mathbf{Z}(\mathbf{C} \circ \mathbf{W}) \Big) \mathbf{Z} - \mathbf{Diag}\big(\hat{\mathbf{k}}^{\Sigma} \circ \mathbf{diag}(\mathring{\mathbf{C}})\big)\mathbf{Z}$ {this is $\mathbf{X}^{(1)} - \mathbf{X}^{(2)} - \mathbf{X}^{(4)}$ in Eq. (C.76)}
15. $\mathbf{X}^{(3)} \leftarrow \Big( \widehat{\mathbf{K}}^{\mathsf{T}} - \mathbf{Diag}(\hat{\mathbf{k}}^{\Sigma}) \Big) \mathbf{Z}$ {see Eq. (C.71)}
16. $\boldsymbol{\Sigma}^{k-\mathrm{RSP}} \leftarrow \mathbf{0}$ {initialize the negative covariance matrix as a $n \times n$ matrix of 0's}
17. **for all** $(i,j) \in \mathcal{E}$ **do** {perform a loop over edges in order to fill in the elements $i$, $j$ of the negative covariance matrix, the $\sigma_{ij}^{(k)}$}
18. $\quad$ $\mathbf{z}_j^{\mathrm{r}} \leftarrow \mathbf{Z}^{\mathsf{T}}\mathbf{e}_j$ {row $j$ of $\mathbf{Z}$ as column vector; see Eq. (C.75)}
19. $\quad$ $\mathbf{x}_i^{(3)\mathrm{c}} \leftarrow \mathbf{X}^{(3)}\mathbf{e}_i$ {column $i$ of $\mathbf{X}^{(3)}$; see Eq. (C.75)}
20. $\quad$ $\mathrm{bet}_{ij}^{k-\mathrm{RSP}} \leftarrow w_{ij} \cdot (\mathbf{z}_j^{\mathrm{r}})^{\mathsf{T}} \mathbf{x}_i^{(3)\mathrm{c}}$ {betweenness of edge $(i,j)$, i.e. element $i$, $j$ of matrix $\mathbf{Bet}^{k-\mathrm{RSP}}$; see Eq. (C.71) and the note following the equation}
21. $\quad$ $\mathbf{y}_j^{\mathrm{r}} \leftarrow \mathbf{Y}^{\mathsf{T}}\mathbf{e}_j$ {row $j$ of $\mathbf{Y}$ as column vector; see Eq. (C.75)}
22. $\quad$ $\mathbf{x}_i^{(5)\mathrm{c}} \leftarrow \mathbf{X}^{(5)}\mathbf{e}_i$ {column $i$ of $\mathbf{X}^{(5)}$; see Eq. (C.75)}
23. $\quad$ $\sigma_{ij}^{k-\mathrm{RSP}} \leftarrow w_{ij} \cdot \Big( (\mathbf{z}_j^{\mathrm{r}})^{\mathsf{T}} \mathbf{x}_i^{(5)\mathrm{c}} - c_{ij} \cdot (\mathbf{z}_j^{\mathrm{r}})^{\mathsf{T}} \mathbf{x}_i^{(3)\mathrm{c}} - (\mathbf{y}_j^{\mathrm{r}})^{\mathsf{T}} \mathbf{x}_i^{(3)\mathrm{c}} \Big)$ {negative covariance of edge $(i,j)$, i.e. element $i$, $j$ of $\boldsymbol{\Sigma}^{k-\mathrm{RSP}}$, $\sigma_{ij}^{(k)}$ in the text; see Eq. (C.75)}
24. **end for**
25. $\mathbf{S}^c \leftarrow \mathbf{Bet}^{k-\mathrm{RSP}} + \theta \boldsymbol{\Sigma}^{k-\mathrm{RSP}}$ {sensitivities w.r.t the costs of movement; see Eqs. (C.54) and (C.57)}
26. $\mathbf{S}^a \leftarrow (\mathbf{W} > 0) \circ \big( \big( (\boldsymbol{\Sigma}^{k-\mathrm{RSP}}\mathbf{e}) \div (\mathbf{Ae}) \big) \mathbf{e}^{\mathsf{T}} - \boldsymbol{\Sigma}^{k-\mathrm{RSP}} \div \mathbf{A} \big)$ {sensitivities w.r.t the step probabilities, see Eqs. (C.55) and (C.57); note that we must avoid missing links (division by 0)}
27. **return** $\mathbf{S}^a, \mathbf{S}^c$

**Algorithm 2** Computation of the sensitivities of the summation-based landscape connectivity index based on the weighted power mean proximity w.r.t. both step probability and cost of movement, $\mathbf{S}^a$ and $\mathbf{S}^c$, respectively (Equations (C.87) and (C.83)), as implemented in the `PM_sensitivity` function.

**Input:**

– A directed, strongly connected graph $G$ containing $n$ nodes.
– The $n \times n$ non-negative affinity matrix $\mathbf{A}$ associated to $G$.
– The $n \times n$ non-negative cost matrix $\mathbf{C}$ associated to $G$.
– The inverse temperature parameter $\theta > 0$.
– The $n \times n$ non-negative diagonal matrices $\mathbf{Q}^s$ and $\mathbf{Q}^t$ containing the qualities of nodes $s$ and $t$, resp., on the diagonals.

**Output:**

– The $n \times n$ sparse matrices $\mathbf{S}^a$ and $\mathbf{S}^c$ containing, for each edge $(i,j) \in \mathcal{E}$, the sensitivities $\partial_{c_{ij}} \mathcal{K}^{\mathrm{PM}}_{\mathrm{sum}}(L)$ and $\partial_{a_{ij}} \mathcal{K}^{\mathrm{PM}}_{\mathrm{sum}}(L)$, respectively, at the corresponding elements $(i,j)$, and 0 elsewhere, computed from Equations (C.87) and (C.83).

1. $\mathbf{D} \leftarrow \mathbf{Diag}(\mathbf{Ae})$ {the outdegree row-normalization diagonal matrix; $\mathbf{e}$ is a column vector of 1's; see Kivimäki et al. (2024), Eq. (1)}
2. $\mathbf{P}^{\mathrm{rw}} \leftarrow \mathbf{D}^{-1}\mathbf{A}$ {the reference random walk transition probabilities matrix; see Kivimäki et al. (2024), Eq. (1)}
3. $\mathbf{W} \leftarrow \mathbf{P}^{\mathrm{rw}} \circ \exp(-\theta \mathbf{C})$ {the $\mathbf{W}$ substochastic matrix involving elementwise exponential and multiplication $\circ$; see Kivimäki et al. (2024), Eq. (5)}
4. $\mathbf{Z} \leftarrow (\mathbf{I} - \mathbf{W})^{-1}$ {the fundamental matrix; see Kivimäki et al. (2024), Eq. (8)}
5. $\mathbf{Z}^h \leftarrow \mathbf{Z}\,(\mathbf{Diag}(\mathbf{Z}))^{-1}$ {the survival probability matrix for hitting paths; see Van Moorter et al. (2023a), Eq. (11) in Appendix B, Kivimäki et al. (2024), Eqs. (29-30), or Kivimäki et al. (2014)}
6. $\mathbf{K} \leftarrow \mathbf{Q}^s (\mathbf{Z}^h)^{(1/\theta)} \mathbf{Q}^t$ {the $q$-weighted power mean proximity matrix with elementwise power $(1/\theta)$ (source-target quality weighting is optional; see Eq. (C.4)}
7. $\mathbf{Y}^{(k)} \leftarrow \Big( (\mathbf{K} \div \mathbf{Z})^{\mathsf{T}} - \mathbf{Diag}\big((\mathbf{K}^{\mathsf{T}}\mathbf{e}) \div \mathbf{diag}(\mathbf{Z})\big) \Big) \mathbf{Z}$ {needed for the computation of the $k$-weighted betweenness, see Eq. (C.116); $\mathbf{diag}(\mathbf{Z})$ is a column vector containing the diagonal of $\mathbf{Z}$ and $\div$ is the elementwise division}
8. $\mathbf{Bet}^{k-\mathrm{RSP}} \leftarrow \mathbf{0}$ {initialize the $k$-weighted edge betweenness matrix as a $n \times n$ matrix of 0's}
9. **for all** $(i,j) \in \mathcal{E}$ **do** {perform a loop over edges in order to fill in the elements $i$, $j$ of the $k$-weighted edge betweenness matrix, $\mathrm{bet}_{ij}^{k-\mathrm{RSP}}$}
10. $\quad \mathbf{z}_j^{\mathrm{r}} \leftarrow \mathbf{Z}^{\mathsf{T}}\mathbf{e}_j$ {row $j$ of $\mathbf{Z}$ as column vector; see Eq. (C.118)}
11. $\quad \mathbf{y}_i^{\mathrm{c}} \leftarrow \mathbf{Y}^{(k)}\mathbf{e}_i$ {column $i$ of $\mathbf{Y}^{(k)}$; see Eq. (C.118)}
12. $\quad \mathrm{bet}_{ij}^{k-\mathrm{RSP}} \leftarrow w_{ij} \cdot (\mathbf{z}_j^{\mathrm{r}})^{\mathsf{T}} \mathbf{y}_i^{\mathrm{c}}$ {betweenness of edge $(i,j)$, i.e. element $i, j$ of matrix $\mathbf{Bet}^{k-\mathrm{RSP}}$; see Eq. (C.118)}
13. **end for**
14. $\mathbf{S}^c \leftarrow -\mathbf{Bet}^{k-\mathrm{RSP}}$ {sensitivities w.r.t the costs of movement; see Eq. (C.87)}
15. $\mathbf{S}^a \leftarrow \frac{1}{\theta}(\mathbf{W} > 0) \circ \big( \mathbf{Bet}^{k-\mathrm{RSP}} \div \mathbf{A} - \big( (\mathbf{Bet}^{k-\mathrm{RSP}}\mathbf{e}) \div (\mathbf{Ae}) \big) \mathbf{e}^{\mathsf{T}} \big)$ {sensitivities w.r.t the step probabilities; see Eq. (C.83); we must avoid missing links (division by 0)}
16. **return** $\mathbf{S}^a, \mathbf{S}^c$

3). It follows that the sensitivity w.r.t. edge costs is essentially proportional to the $k$-weighted node betweenness.

Therefore, for the sensitivity w.r.t. edge affinity, we obtain from Equations (C.31) and (C.110) with $k_{st} = k_{st}^{\text{Surv}} = q_s \mathcal{Z}_{st} q_t$,

$$s_{ij}^a \triangleq \frac{\partial \mathcal{K}_{\text{sum}}^{\text{Surv}}(L)}{\partial a_{ij}} = \frac{\partial \sum_{s,t=1}^n q_s q_t k_{st}^{\text{Surv}}}{\partial a_{ij}} \tag{C.81}$$

$$= \sum_{s,t=1}^n q_s q_t \frac{\partial \mathcal{Z}_{st}}{\partial a_{ij}} = \sum_{s,t=1}^n q_s q_t \left( \frac{\bar{n}_{ij}^{st}}{a_{ij}} - \frac{\bar{n}_{i\rightarrow}^{st}}{a_{i\rightarrow}} \right) \mathcal{Z}_{st} \tag{C.82}$$

$$= \frac{\text{bet}_{ij}^{k-\text{RSP}}}{a_{ij}} - \frac{\text{bet}_{i\rightarrow}^{k-\text{RSP}}}{a_{i\rightarrow}}. \tag{C.83}$$

*Appendix C.3.3. Extention to the sensitivity computation of* $\mathcal{K}_{\text{sum}}^{\text{PM}}(L)$

Let us now derive the sensitivity w.r.t. another interesting quantity, called the *weighted power mean* proximity (PM), $k_{st}^{\text{PM}} = \mathcal{Z}_{st}^{1/\theta}$, which was introduced in Kivimäki et al. (2024) (see this paper for more information) and which benefits from some nice properties. As for the expected cost proximity, it considers an exponential link between the proximity and the cost (see Equation (C.2)), but this cost is now the directed *free energy distance*, $\phi_{st} = -\frac{1}{\theta} \log \mathcal{Z}_{st}$, instead of the expected cost, $k_{st}^{\text{PM}} = \exp(-\phi_{st})$ Kivimäki et al. (2024). It turns out that the weighted power mean proximity is closely related to the survival probability, but is better behaved in the limits $\theta \to 0^+$ and $\theta \to \infty$.

By using Equations (C.15) and (C.31), we readily obtain

$$\frac{\partial k_{st}^{\text{PM}}}{\partial c_{ij}} = \frac{\partial \mathcal{Z}_{st}^{1/\theta}}{\partial c_{ij}} = -\bar{n}_{ij}^{st} \mathcal{Z}_{st}^{1/\theta} \quad \text{and} \quad \frac{\partial k_{st}^{\text{PM}}}{\partial a_{ij}} = \frac{\partial \mathcal{Z}_{st}^{1/\theta}}{\partial a_{ij}} = \frac{1}{\theta} \left( \frac{\bar{n}_{ij}^{st}}{a_{ij}} - \frac{\bar{n}_{i\rightarrow}^{st}}{a_{i\rightarrow}} \right) \mathcal{Z}_{st}^{1/\theta} \tag{C.84}$$

which, by proceeding as for the survival proximity, leads to the following sensitivities,

$$s_{ij}^c \triangleq \frac{\partial \mathcal{K}_{\text{sum}}^{\text{PM}}(L)}{\partial c_{ij}} = \frac{\partial \sum_{s,t=1}^n q_s q_t k_{st}^{\text{PM}}}{\partial c_{ij}} \tag{C.85}$$

$$= \sum_{s,t=1}^n q_s q_t \frac{\partial \mathcal{Z}_{st}^{1/\theta}}{\partial c_{ij}} = \sum_{s,t=1}^n q_s q_t (-\bar{n}_{ij}^{st} \mathcal{Z}_{st}^{1/\theta}) \tag{C.86}$$

$$= -\sum_{s,t=1}^n q_s q_t \mathcal{Z}_{st}^{1/\theta} \bar{n}_{ij}^{st} = -\text{bet}_{ij}^{k-\text{RSP}}, \tag{C.87}$$

This quantity is thus also proportional to the $k$-weighted RSP betweenness computed from the power mean (see Equation (C.110)). Moreover, for sensitivity w.r.t. affinities, we obtain

$$s_{ij}^a \triangleq \frac{\partial \mathcal{K}_{\text{sum}}^{\text{PM}}(L)}{\partial a_{ij}} = \frac{\partial \sum_{s,t=1}^n q_s q_t k_{st}^{\text{PM}}}{\partial a_{ij}} \tag{C.88}$$

$$= \sum_{s,t=1}^n q_s q_t \frac{\partial \mathcal{Z}_{st}^{1/\theta}}{\partial a_{ij}} = \frac{1}{\theta} \sum_{s,t=1}^n q_s q_t \left( \frac{\bar{n}_{ij}^{st}}{a_{ij}} - \frac{\bar{n}_{i\rightarrow}^{st}}{a_{i\rightarrow}} \right) \mathcal{Z}_{st}^{1/\theta} \tag{C.89}$$

$$= \frac{1}{\theta} \left( \frac{\text{bet}_{ij}^{k-\text{RSP}}}{a_{ij}} - \frac{\text{bet}_{i\rightarrow}^{k-\text{RSP}}}{a_{i\rightarrow}} \right), \tag{C.90}$$

which is very close to the result obtained for the survival proximity (Equation C.83), but now weighted by $1/\theta$ and the proximity is computed from the power mean similarity instead of the survival probabilities. In matrix form, we have

$$\mathbf{S}^a = \frac{1}{\theta}\Big(\big(\mathbf{Bet}^{k-\mathrm{RSP}} \div \mathbf{A}\big) - \big((\mathbf{Bet}^{k-\mathrm{RSP}}\mathbf{e}) \div (\mathbf{Ae})\big)\mathbf{e}^\mathsf{T}\Big) \tag{C.91}$$

The algorithm for computing the power mean sensitivities is presented in Algorithm 2. Note that, in this equation, missing nodes must be avoided when computing the elementwise division by matrix $\mathbf{A}$.

*Appendix C.4. Sensitivity of eigenvalue landscape functionality*

Recall that the eigenvalue landscape functionality index is defined in Equation (A.4) as $\mathcal{K}_{\mathrm{eig}} = \lambda$, where $\lambda$ is the leading eigenvalue of the landscape matrix $\mathbf{M}$. As shown e.g. by Caswell (2019, Equation (3.29)), if $\mathbf{v}$ and $\mathbf{w}$ are, respectively, the left and right eigenvectors associated with the largest eigenvalue $\lambda$ of a positive-valued matrix $\mathbf{M}$, then the sensitivity of $\lambda$ to a perturbation of an element $m_{st}$ for any $s, t \in \{1, \ldots, n\}$ is

$$\frac{\mathrm{d}\lambda}{\mathrm{d}m_{st}} = \frac{v_s w_t}{\mathbf{v}^\mathsf{T}\mathbf{w}} \tag{C.92}$$

given that the change in $m_{st}$ does not imply a change in other elements of matrix $\mathbf{M}$. Ovaskainen and Hanski (2003) used this relationship for the sensitivity of the MC to the perturbation of a landscape patch. Hence, the sensitivity of $\mathcal{K}$ to an arbitrary parameter $x_j$ on node $j$ (where $x$ is one of $q$, $a$ or $c$) can be computed with the help of Equation (C.92) and the chain rule, as here the perturbation of $x_j$ can cause a change involving multiple elements of the matrix $\mathbf{M}$. Thus,

$$\frac{\mathrm{d}\lambda}{\mathrm{d}x_j} = \sum_{s,t=1}^{n} \frac{\partial\lambda}{\partial m_{st}} \cdot \frac{\mathrm{d}m_{st}}{\mathrm{d}x_j} = \frac{1}{\mathbf{v}^\mathsf{T}\mathbf{w}} \sum_{s,t=1}^{n} v_s w_t \frac{\mathrm{d}m_{st}}{\mathrm{d}x_j}, \tag{C.93}$$

where Equation (C.92) gives the second equality, as each partial derivative $\partial\lambda/\partial m_{st}$, by definition of the partial derivative, can be considered independently of the changes in other elements of matrix $\mathbf{M}$. Equation (C.93) can also be obtained by writing the expression by Caswell (2019, Equation (3.42)) in elementwise form.

The expressions for the sensitivity of the eigenvalue landscape functionality, $\mathcal{K}_{\mathrm{eig}}(L)$ in Table C.3, are obtained by inserting the expressions of the partial sensitivities of the proximities $k^{\mathrm{Surv}}$ and $k^{\mathrm{EC}}$ w.r.t. $q_j$, $a_j$, $c_j$, and the total sensitivity w.r.t. $c_j$, from the previous sections, into Equation (C.93) and (C.7), where $m_{st} = q_s k_{st} q_t$ and $k_{st}$ can be any of the proximities studied before (expected cost, survival probability, power mean) – the resulting expression is therefore of the same form as before, see Equation (C.7). In other words, here, we do not consider the sum of the weighted proximities (elements of the $\mathbf{M}$ matrix) as in Equations (C.3), (C.4) any more, but the derivative of the leading eigenvalue of the non-negative $\mathbf{M}$ matrix.

*Appendix C.4.1. Sensitivity of $\mathcal{K}_{\mathrm{eig}}(L)$ to change in $q$*

Here, the sensitivity of the eigenvalue landscape functionality index $\mathcal{K}_{\mathrm{eig}}$ w.r.t. the quality $q_j$ of node $j$ is derived. For this, recall that in our consideration, the elements of the landscape matrix, $\mathbf{M}$, are defined as $m_{st} = q_s q_t k_{st}$. Also, note that the derivative of $m_{st}$ w.r.t. $q_j$ is nonzero only if $s = j$ or $t = j$ (or both), and that the proximities $k_{st}$ are independent of $q_j$

and thus unaffected by the derivative. Accordingly, from Equation (C.12), the eigensensitivity expression of Equation (C.93) can be written w.r.t. the quality $q_j$ of node $j$, as

$$\frac{\mathrm{d}\mathcal{K}_{\mathrm{eig}(L)}}{\mathrm{d}q_j} = \frac{1}{\mathbf{v}^{\mathsf{T}}\mathbf{w}} \sum_{s,t=1}^{n} v_s w_t k_{st} \frac{\mathrm{d}(q_s q_t)}{\mathrm{d}q_j} \tag{C.94}$$

$$= \frac{1}{\mathbf{v}^{\mathsf{T}}\mathbf{w}} \sum_{t=1}^{n} q_t (v_t w_j k_{tj} + v_j w_t k_{jt}). \tag{C.95}$$

This result produces the expressions for the sensitivity of $\mathcal{K}_{\mathrm{eig}}(L)$ w.r.t. $q_j$ presented in Table C.3 for both proximity measures $k^{\mathrm{Surv}}$ and $k^{\mathrm{EC}}$.

*Appendix C.4.2. Sensitivity of $\mathcal{K}_{\mathrm{eig}}(L)$ to change in $a$ or $c$*

Then, the sensitivity of the eigenvalue landscape functionality index $\mathcal{K}_{\mathrm{eig}}$ w.r.t. the cost $c_j$, and the step probability $a_j$ of node $j$ are derived, when assuming that $c_j$ and $a_j$ are independent of each other. Remembering also that $q_s$ and $q_t$ are independent of both $c_j$ and $a_j$, and that we consider $m_{st} = q_s q_t k_{st}$, these sensitivity measures can be written based on Equation (C.93) as the partial derivatives

$$\frac{\partial\mathcal{K}_{\mathrm{eig}(L)}}{\partial c_j} = \frac{1}{\mathbf{v}^{\mathsf{T}}\mathbf{w}} \sum_{s,t=1}^{n} q_s v_s q_t w_t \frac{\partial k_{st}}{\partial c_j}, \tag{C.96}$$

and

$$\frac{\partial\mathcal{K}_{\mathrm{eig}(L)}}{\partial a_j} = \frac{1}{\mathbf{v}^{\mathsf{T}}\mathbf{w}} \sum_{s,t=1}^{n} q_s v_s q_t w_t \frac{\partial k_{st}}{\partial a_j}. \tag{C.97}$$

Combining Equations (C.96) and (C.97) with the corresponding equations derived in Appendix C.2 for the partial and total derivatives of the different forms of $k_{st}$ results in the expressions presented in Table C.3 for the eigensensitivities w.r.t. $a_j$ and $c_j$.

*Appendix C.4.3. Connection between sensitivity of summation- and eigenvalue-based landscape functionality*

From the Equations presented here, it is evident that the expression for the sensitivities of the summation- and eigenvalue-based landscape functionality indices are highly similar. In fact, a direct connection between the two can be drawn by considering a scenario where matrix $\mathbf{M}$ is replaced by a new matrix, $\widehat{\mathbf{M}}$, with elements $\widehat{m}_{st} = \frac{v_s w_t}{\mathbf{v}^{\mathsf{T}}\mathbf{w}} m_{st}$, and then assuming that the values $\widehat{m}_{st}$, for all $s, t \in \mathcal{V}$, are independent of the parameters $q_j$, $c_j$ and $a_j$ for all $j \in \mathcal{V}$. Of course, normally the eigenvectors $\mathbf{v}$ and $\mathbf{w}$ do depend on the parameter values, as they are eigenvectors of $\mathbf{M}$ which depends on all the parameters. Nevertheless, if we define matrix $\widehat{\mathbf{M}}$ as above, with the assumption of independence between $\widehat{m}_{st}$ and the parameters, i.e. that the values $v_j$ and $w_j$ for all $j \in \mathcal{V}$, are considered as constant when taking derivatives, then the eigen sensitivity based on matrix $\mathbf{M}$ is equivalent to the summation sensitivity based on matrix $\widehat{\mathbf{M}}$. Note that this is not the same as saying that the eigenvalue landscape functionality index based on $\mathbf{M}$ is the same as the summation-based landscape functionality index based on $\widehat{\mathbf{M}}$, but only that their derivatives are equivalent.

*Appendix C.5. Sensitivity of circuit-based landscape functionality*

In this section, we also briefly review the sensitivity of landscape functionality w.r.t. conductance in the context of circuit theory Klein (2010); Ghosh et al. (2008). Circuit theory is a popular framework for connectivity modeling (McRae and Beier, 2007; Dickson et al., 2019), with the landscape as a conductive surface where each cell has a conductivity (inverse of resistance) value representing ease of movement; it is mathematically equivalent to modeling random walks on an undirected graph Doyle and Snell (1984). A commonly used distance metric derived from a circuit theory perspective is the resistance distance (Klein and Randic (1993), which is proportional to the commute cost, i.e. the symmetrized hitting cost Kivimäki et al. (2014)), which can be used to define landscape functionality from circuit theory:

$$\mathcal{K}^{\mathrm{R}}_{\mathrm{sum}}(L) = \sum_{s=1}^{n}\sum_{t=1}^{n} q_s q_t k^R_{st} = \sum_{s=1}^{n}\sum_{t=1}^{n} q_s q_t \exp(-\alpha R_{st}) \tag{C.98}$$

where $k^R_{st}$ is the proximity based on the resistance distance, $R_{st}$, between $s$ and $t$.

As shown in Klein (2010) (see also Ghosh et al. (2008)), the sensitivity of the resistance distance $R_{st}$ w.r.t. a perturbation of the conductance, $a_{ij}$, is directly related to the power dissipated on an edge ($P_{ij} = I^2_{ij}/a_{ij}$, where $I_{ij}$ is the current through edge $(i, j)$:

$$\frac{\mathrm{d}R_{st}}{\mathrm{d}a_{ij}} = -\left(\frac{I_{ij}}{a_{ij}}\right)^2 = -\frac{P_{ij}}{a_{ij}}. \tag{C.99}$$

Using Equation C.99 and the chain rule, the sensitivity of the resistance proximity, $k^R_{st}$, w.r.t. $a_{ij}$ becomes

$$\frac{\mathrm{d}k^{\mathrm{R}}_{st}}{\mathrm{d}a_{ij}} = \frac{\partial \exp(-\alpha R_{st})}{\partial R_{st}} \cdot \frac{\partial R_{st}}{\partial a_{ij}} = -\alpha \cdot \exp(-\alpha R_{st}) \cdot \frac{\partial R_{st}}{\partial a_{ij}} = \alpha k^{\mathrm{R}}_{st} \frac{P^{st}_{ij}}{a_{ij}} = \frac{\alpha k^{\mathrm{R}}_{st}}{a_{ij}} P^{st}_{ij}. \tag{C.100}$$

Thus, the sensitivity of the circuit-based landscape functionality, $\mathcal{K}^{\mathrm{R}}_{\mathrm{sum}}(L)$, to perturbation of the conductance $a_{ij}$, can be quantified with the derivative:

$$\frac{\mathrm{d}\mathcal{K}^{\mathrm{R}}_{\mathrm{sum}}(L)}{\mathrm{d}a_{ij}} = \sum_{s,t=1}^{n} q_s q_t \cdot \frac{\mathrm{d}k^R_{st}}{\mathrm{d}a_{ij}} = \frac{1}{a_{ij}} \sum_{s,t=1}^{n} q_s q_t \cdot \alpha k^{\mathrm{R}}_{st} \cdot P^{st}_{ij}. \tag{C.101}$$

The scale-free sensitivity or elasticity of the circuit-based landscape functionality is then:

$$a_{ij} \cdot \frac{\mathrm{d}\mathcal{K}^{\mathrm{R}}_{\mathrm{sum}}(L)}{\mathrm{d}a_{ij}} = \sum_{s,t=1}^{n} q_s q_t \cdot \alpha k^{\mathrm{R}}_{st} \cdot P^{st}_{ij}. \tag{C.102}$$

which is closely related to power centrality Klein (2010).

Indeed, the result (C.99) has been previously used by both Klein (2010) and Ghosh et al. (2008). Let us briefly summarize the arguments. It is well-known that the resistance distance can be computed from the Moore-Penrose pseudoinverse of the positive semi-definite Laplacian matrix Klein and Randic (1993),

$$R_{st} = (\mathbf{e}_s - \mathbf{e}_t)^{\mathsf{T}} \mathbf{L}^{+} (\mathbf{e}_s - \mathbf{e}_t). \tag{C.103}$$

where $\mathbf{e}_k$ is a column vector containing a 1 at position $k$ and 0 elsewhere. From this equation, it can be shown Klein (2010) that the derivative of the resistance distance with respect to the conductance of an existing edge $(i, j) \in \mathcal{E}$ can be computed in terms of the elements of the Laplacian matrix as

$$\frac{\mathrm{d}R_{st}}{\mathrm{d}a_{ij}} = \left(l^+_{si} - l^+_{sj} + l^+_{tj} - l^+_{ti}\right)^2. \tag{C.104}$$

In addition, when deriving the power centrality Klein (2010), we obtain that the electrical power dissipated at the level of an edge $(i, j)$ (see, e.g., Fouss et al. (2016), Equation (4.119)) is

$$P^{st}_{ij} = a_{ij}\left(l^+_{si} - l^+_{sj} + l^+_{tj} - l^+_{ti}\right)^2, \tag{C.105}$$

which justifies Equation (C.99).

*Appendix C.6. Sensitivity of least-cost-based landscape functionality*

Finally, we consider the sensitivity of landscape functionality w.r.t. cost in the context of the shortest or least-cost path (Ahuja and Orlin, 2001; Klein, 2010), which is also a popular framework for connectivity modeling in ecology. The distance metric in this framework is the cost of the shortest path, which can be used to define landscape functionality:

$$\mathcal{K}^{\mathrm{R}}_{\mathrm{sum}}(L) = \sum_{s=1}^{n}\sum_{t=1}^{n} q_s q_t k^{LC}_{st} = \sum_{s=1}^{n}\sum_{t=1}^{n} q_s q_t \exp(-\alpha d_{st}) \tag{C.106}$$

where $k^{LC}_{st}$ is the proximity based on the least-cost distance, $d_{st}$, between $s$ and $t$.

Based on Equation 7.3 in Ahuja and Orlin (2001), and making the assumption that there is only one shortest path from source to target, the sensitivity of the least-cost distance $d_{st}$ w.r.t. a perturbation of the cost, $c_{ij}$, is 1 for edges on the shortest path and 0 elsewhere:

$$\frac{\mathrm{d}d_{st}}{\mathrm{d}c_{ij}} = \begin{cases} 1 & \text{if edge } (i, j) \in P^*_{st} \\ 0 & \text{otherwise} \end{cases} \tag{C.107}$$

with $P^*_{st}$ the unique shortest path from $s$ to $t$. The cost distance function is piecewise linear and non-differentiable at points where the shortest path structure changes (e.g., two paths are equally optimal). The extensions to several shortest paths are not trivial and are investigated in Ahuja and Orlin (2001); Klein (2010); these are not discussed in the present paper.

Using Equation C.107 and the chain rule, the sensitivity of the least-cost proximity, $k^{\mathrm{LC}}_{st}$, w.r.t. $c_{ij}$ becomes

$$\frac{\mathrm{d}k^{\mathrm{LC}}_{st}}{\mathrm{d}c_{ij}} = \frac{\partial \exp(-\alpha d_{st})}{\partial d_{st}} \cdot \frac{\partial d_{st}}{\partial c_{ij}} = -\alpha \cdot \exp(-\alpha d_{st}) \cdot \frac{\partial d_{st}}{\partial c_{ij}} = \begin{cases} -\alpha k^{\mathrm{LC}}_{st} & \text{if edge } i\text{-}j \in P^*_{st} \\ 0 & \text{otherwise} \end{cases}. \tag{C.108}$$

Thus, the sensitivity of the least-cost-based landscape functionality, $\mathcal{K}^{\mathrm{LC}}_{\mathrm{sum}}(L)$, to perturbation of the step cost $c_{ij}$ can be quantified with the derivative:

$$\frac{\mathrm{d}\mathcal{K}^{\mathrm{LC}}_{\mathrm{sum}}(L)}{\mathrm{d}c_{ij}} = \sum_{s,t=1}^{n} q_s q_t \cdot \frac{\mathrm{d}k^{\mathrm{LC}}_{st}}{\mathrm{d}c_{ij}} = -\sum_{s,t=1}^{n} q_s q_t \cdot \alpha k^{\mathrm{LC}}_{st} \cdot \begin{cases} 1 & \text{if edge } i\text{-}j \in P^*_{st} \\ 0 & \text{otherwise} \end{cases}. \tag{C.109}$$

Thus, the sensitivity of the least-cost landscape functionality w.r.t. perturbation of the step cost is the quality- and proximity-weighted betweenness, referred to as the ‘generalized betweenness’ centrality by Bodin and Saura (2010).

#### POSTSCRIPT

*Alternative derivation for the proximity-weighted RSP edge betweenness*

In Van Moorter et al. (2023a), we presented a derivation and algorithm for the 'proximity-weighted RSP edge betweenness', or '$k$-weighted betweenness' in short, in Section C.2.4 from Appendix C. While this derivation and algorithm are correct, they do not correspond exactly to the implementation in the ConScape library. Therefore, we present here the alternative derivation and algorithm corresponding to this implementation of RSP edge betweenness including a weighting of the flow between each source and target nodes with the proximity between them, $k_{st}$. Using again Equation (11) in Kivimäki et al. (2024) (see also Equation (C.59)) where $k_{st}$ can be any proximity measure (such as expected cost, survival, or weighted power mean),

$$\begin{aligned}
\mathrm{bet}_{ij}^{k-\mathrm{RSP}} &= \sum_{s,t=1}^{n} \bar{n}_{ij}^{st} k_{st} = \sum_{s,t=1}^{n} \left( \frac{z_{si}}{z_{st}} - \frac{z_{ti}}{z_{tt}} \right) w_{ij} z_{jt} k_{st} && \text{(C.110)} \\
&= \left( \sum_{s,t=1}^{n} z_{jt} \frac{k_{ts}^{\mathsf{T}}}{z_{ts}^{\mathsf{T}}} z_{si} - \sum_{s,t=1}^{n} z_{jt} \frac{z_{ti}}{z_{tt}} k_{st} \right) w_{ij} && \text{(C.111)} \\
&= \left( \sum_{s,t=1}^{n} z_{jt} \frac{k_{ts}^{\mathsf{T}}}{z_{ts}^{\mathsf{T}}} z_{si} - \sum_{t=1}^{n} z_{jt} \frac{z_{ti}}{z_{tt}} \left( \sum_{s=1}^{n} k_{st} \right) \right) w_{ij} && \text{(C.112)} \\
&= \left( \sum_{s,t=1}^{n} z_{jt} \frac{k_{ts}^{\mathsf{T}}}{z_{ts}^{\mathsf{T}}} z_{si} - \sum_{t=1}^{n} z_{jt} \frac{k_{\rightarrow t}}{z_{tt}} z_{ti} \right) w_{ij} && \text{(C.113)}
\end{aligned}$$

with $k_{\rightarrow t} = \sum_{s=1}^{n} k_{st}$. This can be written in matrix form as

$$\begin{aligned}
\mathrm{bet}_{ij}^{k-\mathrm{RSP}} &= \mathbf{e}_j^{\mathsf{T}} \Big( \mathbf{Z} \big( \mathbf{K} \div \mathbf{Z} \big)^{\mathsf{T}} \mathbf{Z} - \mathbf{Z}\, \mathbf{Diag}\big( (\mathbf{K}^{\mathsf{T}} \mathbf{e}) \div \mathbf{diag}(\mathbf{Z}) \big) \mathbf{Z} \Big) \mathbf{e}_i \cdot w_{ij} && \text{(C.114)} \\
&= \mathbf{e}_j^{\mathsf{T}} \mathbf{Z} \Big( \big( \mathbf{K} \div \mathbf{Z} \big)^{\mathsf{T}} - \mathbf{Diag}\big( (\mathbf{K}^{\mathsf{T}} \mathbf{e}) \div \mathbf{diag}(\mathbf{Z}) \big) \Big) \mathbf{Z} \mathbf{e}_i \cdot w_{ij} && \text{(C.115)}
\end{aligned}$$

where $\mathbf{Diag}(\mathbf{Z})$ is a diagonal matrix containing the diagonal of $\mathbf{Z}$ and $\mathbf{diag}(\mathbf{Z})$ is a column vector, also containing the diagonal of $\mathbf{Z}$. Recall also that $\div$ is the elementwise division. The computation can again benefit from the introduction of a helper matrix[3],

$$\mathbf{Y}^{(k)} = \Big( \big( \mathbf{K} \div \mathbf{Z} \big)^{\mathsf{T}} - \mathbf{Diag}\big( (\mathbf{K}^{\mathsf{T}} \mathbf{e}) \div \mathbf{diag}(\mathbf{Z}) \big) \Big) \mathbf{Z} \qquad \text{(C.116)}$$

Then,

$$\begin{aligned}
\mathrm{bet}_{ij}^{k-\mathrm{RSP}} &= w_{ij} \big( \mathbf{e}_j^{\mathsf{T}} \mathbf{Z} \big) \big( \mathbf{Y}^{(k)} \mathbf{e}_i \big) && \text{(C.117)} \\
&= w_{ij} \big( (\mathbf{z}_j^{\mathrm{r}})^{\mathsf{T}} \mathbf{y}_i^{\mathrm{c}} \big) && \text{(C.118)}
\end{aligned}$$

where $\mathbf{z}_j^{\mathrm{r}}$ is a column vector containing row $j$ of $\mathbf{Z}$ and $\mathbf{y}_i^{\mathrm{c}}$ is column $i$ of $\mathbf{Y}^{(k)}$. This computation is shown as pseudo-code in Algorithm 3 and is implemented in the ConScape library (instead of Algorithm 5 in Section C.2.4 from Appendix C, Van Moorter et al., 2023a).

[3]The helper matrix was chosen slightly differently in (Van Moorter et al., 2023a).

**Algorithm 3** Computation of the $k$-weighted RSP edge betweenness on edges of graph $G$ as defined in Equations (C.113) and (C.115).

**Input:**

– A directed, strongly connected, graph $G$ containing $n$ nodes.
– The $n \times n$ non-negative affinity matrix $\mathbf{A}$ associated to $G$.
– The $n \times n$ non-negative cost matrix $\mathbf{C}$ associated to $G$.
– The $n \times n$ non-negative diagonal matrices $\mathbf{Q}^s$ and $\mathbf{Q}^t$ containing the source and target qualities, respectively, on the diagonals.
– The $n \times n$ non-negative matrix $\mathbf{K}$ of proximities $k_{st}$ between each pair of source node $s$ and target node $t$.
– The inverse temperature parameter $\theta > 0$.

**Output:**

– The $n \times n$ sparse matrix $\mathbf{Bet}^{k-\mathrm{RSP}}$ containing, for each edge $(i, j) \in \mathcal{E}$, the $k$-weighted RSP edge betweenness, $\mathrm{bet}_{ij}^{k-\mathrm{RSP}}$ at the corresponding element $i$, $j$, and 0 elsewhere, computed from Equations (C.115) and (C.118).

$\mathbf{D} \leftarrow \mathbf{Diag}(\mathbf{Ae})$ {the outdegree row-normalization diagonal matrix; $\mathbf{e}$ is a column vector of 1's; see Kivimäki et al. (2024), Eq. (1)}
$\mathbf{P}^{\mathrm{rw}} \leftarrow \mathbf{D}^{-1}\mathbf{A}$ {the reference random walk transition probabilities matrix; see Kivimäki et al. (2024), Eq. (1)}
$\mathbf{W} \leftarrow \mathbf{P}^{\mathrm{rw}} \circ \exp(-\theta\mathbf{C})$ {the $\mathbf{W}$ substochastic matrix involving elementwise exponential and multiplication $\circ$; see Kivimäki et al. (2024), Eq. (6)}
$\mathbf{Z} \leftarrow (\mathbf{I} - \mathbf{W})^{-1}$ {the fundamental matrix; see Kivimäki et al. (2024), Eq. (8)}
$\mathbf{K} \leftarrow \mathbf{Q}^s\mathbf{K}\mathbf{Q}^t$ {the $q$-weighted proximity matrix (optional; only if source-target quality weighting is desired); see Eq. (C.4)}
$\mathbf{Y}^{(k)} \leftarrow \Big((\mathbf{K} \div \mathbf{Z})^{\mathsf{T}} - \mathbf{Diag}\big((\mathbf{K}^{\mathsf{T}}\mathbf{e}) \div \mathbf{diag}(\mathbf{Z})\big)\Big)\mathbf{Z}$ {see Eq. (C.116); we use elementwise division $\div$}
$\mathbf{Bet}^{k-\mathrm{RSP}} \leftarrow \mathbf{0}$ {initialize the $k$-weighted edge betweenness matrix as a $n \times n$ matrix of 0's}
**for all** $(i, j) \in \mathcal{E}$ **do** {perform a loop over edges in order to fill in the elements $i$, $j$ of the $k$-weighted edge betweenness matrix, i.e. $\mathrm{bet}_{ij}^{k-\mathrm{RSP}}$}
    $\mathbf{z}_j^{\mathrm{r}} \leftarrow \mathbf{Z}^{\mathsf{T}}\mathbf{e}_j$ {row $j$ of $\mathbf{Z}$ as column vector; see Eq. (C.118)}
    $\mathbf{y}_i^{\mathrm{c}} \leftarrow \mathbf{Y}^{(k)}\mathbf{e}_i$ {column $i$ of $\mathbf{Y}^{(k)}$; see Eq. (C.118)}
    $\mathrm{bet}_{ij}^{k-\mathrm{RSP}} \leftarrow w_{ij} \cdot (\mathbf{z}_j^{\mathrm{r}})^{\mathsf{T}}\mathbf{y}_i^{\mathrm{c}}$ {betweenness of edge $(i, j)$, i.e. element $i$, $j$ of matrix $\mathbf{Bet}^{k-\mathrm{RSP}}$; see Eq. (C.118)}
**end for**
**return** $\mathbf{Bet}^{k-\mathrm{RSP}}$ {returns the $k$-weighted RSP edge betweenness matrix}

### Appendix D. Demonstration notebook

#### Introduction

This notebook provides a reproducible demonstration of the analytical sensitivity framework developed in the main manuscript, implemented using the ConScape library in Julia. It illustrates how sensitivities and elasticities of landscape functionality with respect to habitat quality and permeability can be computed efficiently for large spatial networks. For background on ConScape and practical guidance on its use, we refer readers to Van Moorter et al. (2023; https://doi.org/10.1111/2041-210X.13850) and the project website (https://conscape.org/).

We first demonstrate the computation of sensitivities and elasticities based on the summation formulation of the landscape matrix, which is used throughout the main text. We then compare these infinitesimal sensitivity measures with the finite effects of complete node removal, referred to as node criticality, thereby illustrating how analytical sensitivities approximate the spatial patterns revealed by large perturbations.

Next, we extend the analysis to sensitivities derived from the eigenvalue-based formulation of landscape functionality corresponding to metapopulation capacity. This comparison illustrates the close correspondence between summation-based and eigenvalue-based sensitivity patterns and motivates the use of summation-based sensitivities as scalable surrogates in large, high-resolution landscapes. Finally, we demonstrate how the harmonic-mean correction term appearing in the mean-field persistence condition derived in Appendix B can be incorporated into the sensitivity calculations.

Together, these examples illustrate the practical implementation, interpretability, and computational advantages of analytical sensitivity analysis for assessing landscape functionality and supporting spatial prioritization.

#### Setup

Load the necessary libraries:

```
using Pkg
Pkg.add(url = "https://github.com/ConScape/ConScape.jl.git",
                                                rev = "sensitivity")
```

```
using ConScape#sensitivity
using SparseArrays

using DataFrames, Statistics, StatsBase, PrettyTables
using Plots

function NaNcor(x, y)
    non_nans = findall(@. !(isnan(x) | isnan(y)))
    return(cor(x[non_nans], y[non_nans]))
end
```

Read the input data (see Figure 1):

```
datadir = dirname(dirname(pathof(ConScape)))

mov_prob, meta_p =
        ConScape.readasc(joinpath(datadir,"data", "mov_prob_1000.asc"))
hab_qual, meta_q =
        ConScape.readasc(joinpath(datadir,"data", "hab_qual_1000.asc"))

non_matches = findall(xor.(isnan.(mov_prob), isnan.(hab_qual)))
mov_prob[non_matches] .= NaN
hab_qual[non_matches] .= NaN;
```

Generate the ConScape Grid (g) from the input data and the GridRSP (h) for a level of randomness ($\theta = 1.5$):

```
g = ConScape.Grid(size(mov_prob)...,
                    affinities=ConScape.graph_matrix_from_raster(mov_prob),
                    source_qualities=hab_qual,
                    target_qualities=hab_qual,
                    costs=ConScape.MinusLog())

h = ConScape.GridRSP(g,  =1.5);
```

```
[ Info: cost graph contains 4927 strongly connected subgraphs
[ Info: removing 4926 nodes from affinity and cost graphs
```

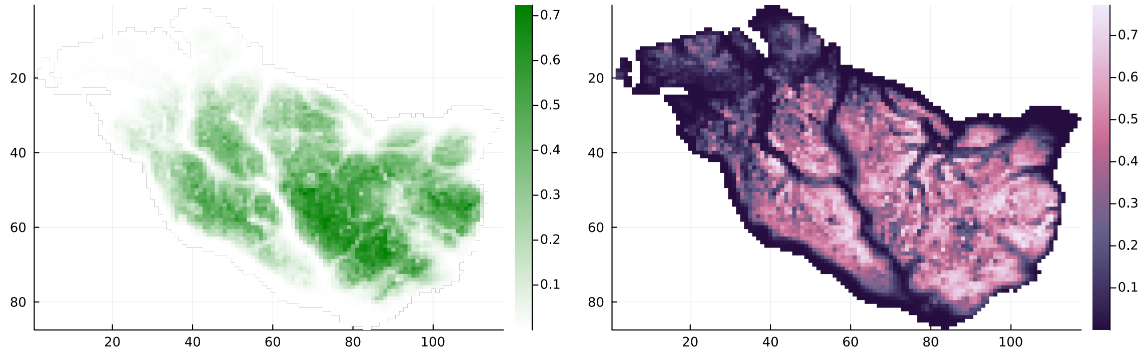


(a) Habitat quality

(b) Habitat permeability

Figure 1: Input maps of habitat quality and permeability.

Note that the greek symbols (here, $\theta$) do not get rendered properly in the code blocks (it should read $\theta = 1.5$ in the code above).

#### Sensitivity analysis

##### With respect to habitat quality

We compute the sensitivity w.r.t. quality by setting the 'wrt' argument to "Q":

```
sens_Q = ConScape.sensitivity(h,
                    connectivity_function=ConScape.expected_cost,
                    distance_transformation=ConScape.ExpMinus(),
                     =1/75.,
                    wrt="Q", unitless=false);
```

(see Figure 2), note that $\alpha$ symbol doesn't seem to be rendered properly in the code blocks (we set $\alpha = 1/75$ in this demonstration).

##### With respect to permeability

The effect of perturbation of the permeabilility can be computed in serval ways: only w.r.t. step cost (wrt="C"), only w.r.t. step likelihood (wrt="A"), w.r.t. the step cost as a function of the likelihood (and vice versa) by setting wrt="A&C=f(A)" (or wrt="C&A=f(C)" respectively). The functional relationship is derived from the 'g.costfunction' in the ConScape Grid g. Here we demonstrate the wrt="A&C=f(A)" version:

```
sens_AfC = ConScape.sensitivity(h,
                    connectivity_function=ConScape.expected_cost,
                    distance_transformation=ConScape.ExpMinus(),
                     =1/75.,
                    wrt="A&C=f(A)", unitless=false);
```

See Figure 2.

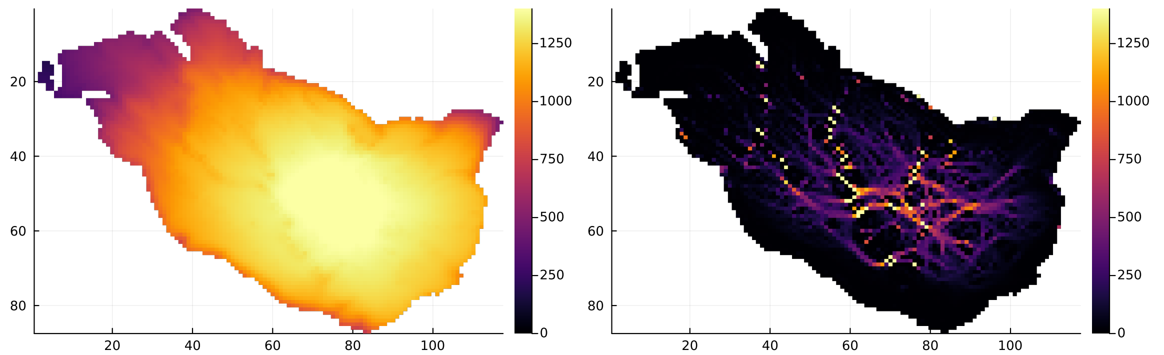


(a) w.r.t. quality

(b) w.r.t. permeability

Figure 2: Sensitivity w.r.t. quality and permeability.

#### Sensitivity analysis using elasticities

The elasticities represent the proportional change, whereas the sensitivities above are the effects of unit change. We compute elasticities by setting the 'unitless' argument to true.

##### With respect to habitat quality

```
elast_Q = ConScape.sensitivity(h,
                    connectivity_function=ConScape.expected_cost,
                    distance_transformation=ConScape.ExpMinus(),
                     =1/75.,
                    wrt="Q", unitless=true);
```

See Figure 3.

##### With respect to permeability

```
elast_AfC = ConScape.sensitivity(h,
                    connectivity_function=ConScape.expected_cost,
                    distance_transformation=ConScape.ExpMinus(),
                     =1/75.,
                    wrt="A&C=f(A)", unitless=true);
```

See Figure 3.

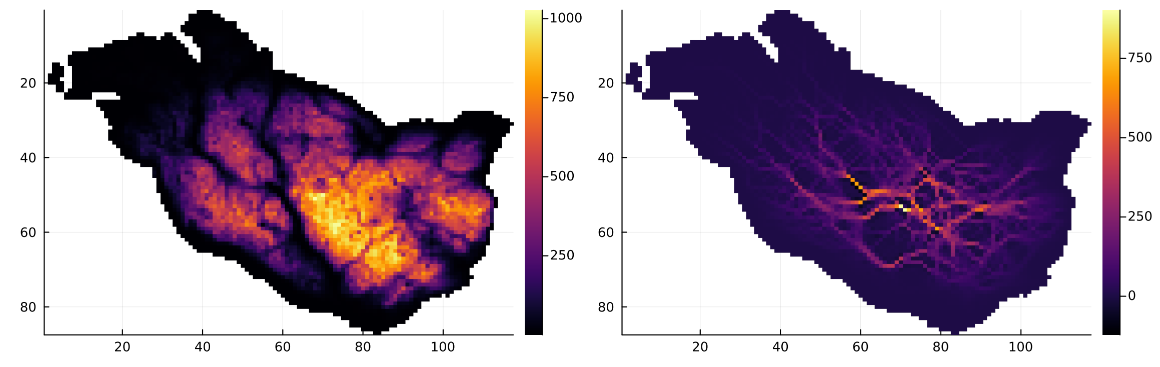


(a) w.r.t. quality

(b) w.r.t. permeability

Figure 3: Elasticity w.r.t. quality and permeability.

#### Criticality

The criticality of a node is the effect of the complete removal of a node (either only its habitat or its complete removal from the network). The criticality computation takes a long time, as it involves the iterative removal of nodes. We added the argument `one_out_of` to evaluate only one out of every so many nodes. Here we used 1 out of every 500 pixels for faster computation: `one_out_of`=500. However, to show reliable results, we load the results where every location is used (`one_out_of`=1), made available through precomputation (takes about 24 hours).

```
crit_Q = ConScape.criticality_simulation(h,  =1/75., wrt="Q", one_out_of=500);
```

See Figure 4. The correlation between the criticality and the sensitivity w.r.t. habitat quality is -0.645, and the elasticity -1.0

```
crit_AC = ConScape.criticality_simulation(h,  =1/75., wrt="all", one_out_of=500);
```

The correlation between the criticality w.r.t. quality and complete node removal is very high: 0.996, due to the fact that in both version habitat quality is removed.

We compute the difference between the complete removal and habitat only removal to isolate the effect of destruction of the connectivity from the destruction of habitat:

```
crit_dAC = crit_AC .- crit_Q;
```

See Figure 4. The correlation between the criticality and the sensitivity w.r.t. cost of movement: -0.002 and the elasticity: -0.699

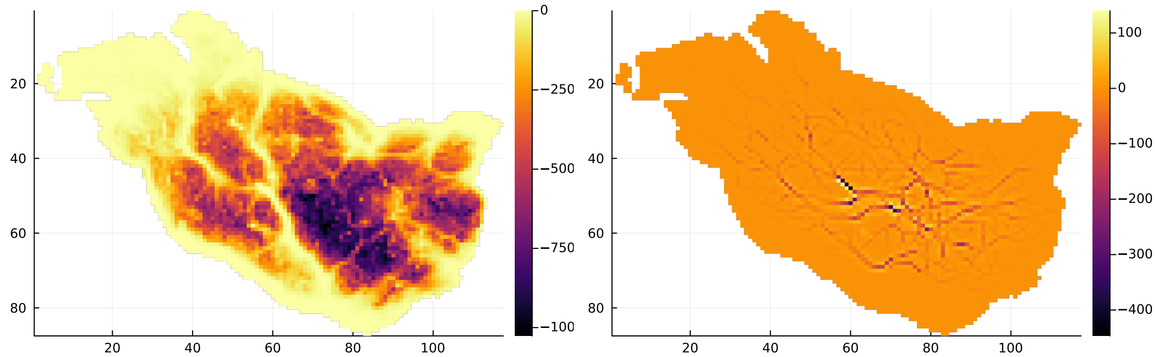


(a) w.r.t. quality (b) w.r.t. permeability

Figure 4: Criticality w.r.t. quality and permeability.

#### Sensitivity analysis for eigenanalysis based landscape summary

All previous sensitivity analysis were based on a summation-based topological measure, i.e. all elements in the landscape matrix were summed, which is the default. We now demonstrate the sensitivity computation for an eigenanalysis based topological measure, i.e. the dominant eigenvalue of the landscape matrix, which correponds to the metapopulation capacity (Hanski & Ovaskainen, 2000), we do this by setting the 'landscape_measure' argument to "eigenanalysis".

##### With respect to habitat quality

```
sens_Q_eigen = ConScape.sensitivity(h,
                          connectivity_function=ConScape.expected_cost,
                          distance_transformation=ConScape.ExpMinus(),
                           =1/75.,
                          wrt="Q",
                          landscape_measure="eigenanalysis",
                          unitless=false);
```

See Figure 5, correlation between the summation and eigenanalysis based sensitivity: 0.638.

###### With respect to permeability

```
sens_AfC_eigen = ConScape.sensitivity(h,
                          connectivity_function=ConScape.expected_cost,
                          distance_transformation=ConScape.ExpMinus(),
                           =1/75.,
                          wrt="A&C=f(A)",
                          landscape_measure="eigenanalysis",
                          unitless=false);
```

See Figure 5, correlation between the summation and eigenanalysis based sensitivity: 0.815.

##### Correction for the Harmonic-Mean Term in the Persistence Condition

As derived in Appendix B, the mean-field persistence condition can be written as:

$$\Phi = \frac{\mathcal{K}_{\text{sum}}(L)}{n\,\overline{q^{-1}}},$$

where the denominator $(n\,\overline{q^{-1}})$ is the harmonic mean of habitat quality.
This term introduces a small correction that gives slightly greater weight to low-quality sites. In practice the effect is very minor, but it is simple to compute and can be included for completeness.

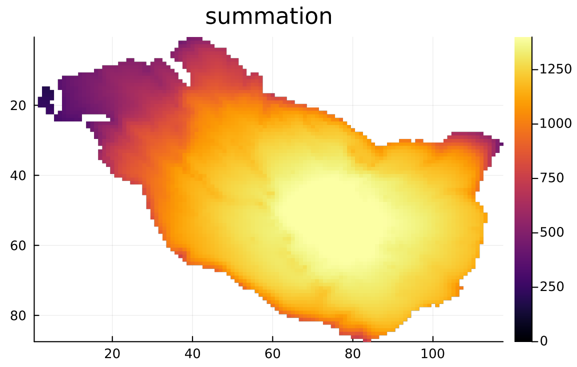


(a) summation-based w.r.t. quality

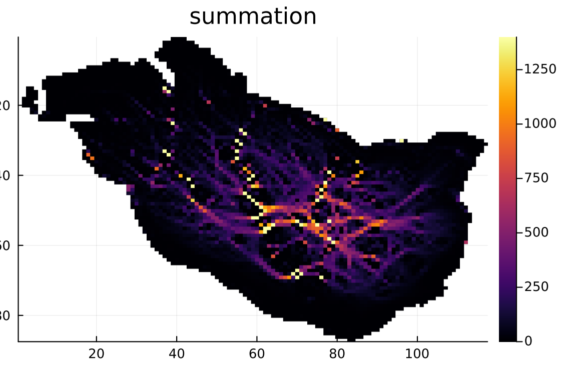


(b) summation-based w.r.t. permeability

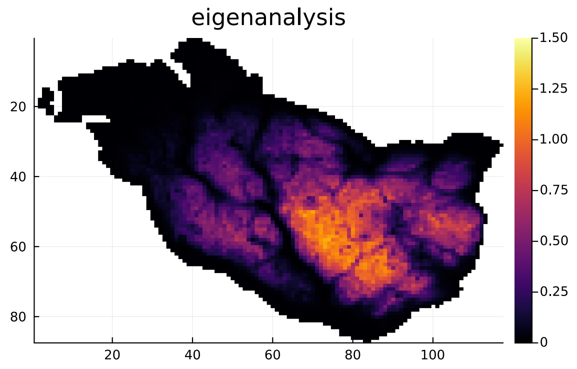


(c) eigenanalysis w.r.t. quality

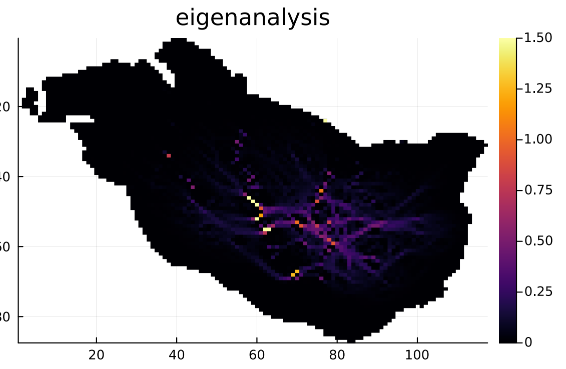


(d) eigenanalysis w.r.t. permeability

Figure 5: Sensitivity w.r.t. quality and permeability based on summation versus on eigenanalysis.

#### Computation

```
# Number of habitat cells
nan_mask = .!isnan.(g.source_qualities)
qualities = g.source_qualities[nan_mask]
n = length(qualities)

# habitat functionality
func = ConScape.connected_habitat(h,
                    connectivity_function = ConScape.expected_cost,
                    distance_transformation=x -> exp(-x/75));

# summation-based landscape functionality
K_sum = sum(func[nan_mask]);

# Harmonic mean of habitat quality
q_inv = mean(1 ./(qualities))

# Mean-field persistence term
phi = K_sum / (n * q_inv)

# Correction term for the derivative with respect to quality
correction_Q = K_sum ./ (n^2 * q_inv^2 .* g.source_qualities.^2)

# Total derivative including the correction
dPhi_dQ = (1 / (n * q_inv)) .* sens_Q .+ correction_Q;
```

This correction term becomes larger for smaller quality values ($q < 10^{-4}$) to very large for extremely small quality values ($q < 10^{-11}$):

| q_min | q_max | n | max correction | median_correction |
|---|---|---|---|---|
| 3.35e-12 | 4.55e-11 | 4 | 90700.0 | 48300.0 |
| 4.55e-11 | 6.19e-10 | 1 | 5.63 | 5.63 |
| 6.19e-10 | 8.42e-9 | 4 | 1.64 | 0.0741 |
| 8.42e-9 | 1.14e-7 | 17 | 0.0113 | 0.00103 |
| 1.14e-7 | 1.56e-6 | 54 | 2.87e-5 | 2.45e-6 |
| 1.56e-6 | 2.12e-5 | 118 | 4.12e-7 | 2.22e-8 |
| 2.12e-5 | 0.000288 | 192 | 2.27e-9 | 1.12e-10 |
| 0.000288 | 0.00391 | 586 | 1.22e-11 | 5.14e-13 |

| | | | | |
|---|---|---|---|---|
| 0.00391 | 0.0532 | 1119 | 6.64e-14 | 4.21e-15 |
| 0.0532 | 0.723 | 3158 | 3.59e-16 | 1.34e-17 |

The total sensitivity including the correction term is very highly correlated with the original sensitivity for pixels with $q > 10^{-4}$: 1.0

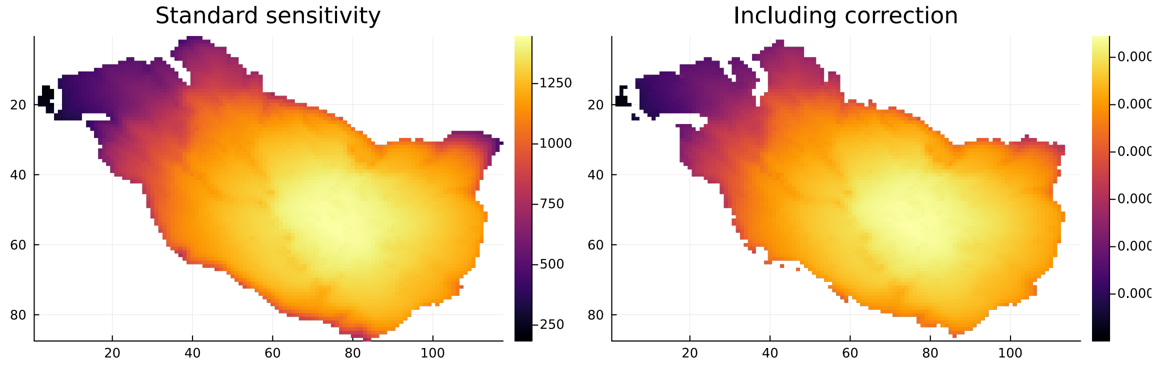


(a) original sensitivities

(b) corrected sensitivities

Figure 6: Compare original and corrected sensitivities.

The corrected derivative corresponds exactly to the theoretical expression derived in Appendix B:

$$\frac{\partial \Phi}{\partial q_t} = \frac{1}{n, \overline{q^{-1}}} \frac{\partial \mathcal{K}_{\text{sum}}}{\partial q_t} * \frac{\mathcal{K}_{\text{sum}}}{n^2, (\overline{q^{-1}})^2, q_t^2}.$$

Figure 6 compares the standard and corrected sensitivities: the standard sensitivity (left) and the version including the harmonic-mean correction (right). Spatial patterns remain virtually identical, confirming that the correction affects only scaling.

For the main text, we report sensitivities of $\mathcal{K}_{\text{sum}}(L)$ alone, as they capture the dominant structural dependence on permeability and habitat quality while keeping the presentation concise. This section shows, however, that incorporating the harmonic-mean correction is both straightforward and fully consistent with the mean-field derivation.

#### Summary

This notebook demonstrated the practical computation of analytical sensitivities and elasticities of landscape functionality using the ConScape framework. We showed that sensitivities and elasticities—quantifying the effects of infinitesimal changes in habitat quality and permeability – provide informative approximations of the finite impacts associated with complete node removal, which we refer to as node criticality. Despite their fundamentally different perturbation scales, sensitivity- and elasticity-based rankings closely reproduce the spatial patterns obtained from node-removal analyses, while being far more computationally efficient.

We further demonstrated that sensitivity analysis can be applied consistently to different landscape functionality measures, including both summation-based formulations and eigenanalysis-based formulations corresponding to metapopulation capacity. Sensitivity patterns derived from these alternative summaries are strongly correlated, indicating that summation-based sensitivities can serve as effective surrogates for eigenvalue-based sensitivities in large, high-resolution landscapes.

Finally, we illustrated how the harmonic-mean correction term arising in the mean-field persistence condition can be incorporated into sensitivity calculations. This correction affects only the scaling of sensitivities and does not alter their spatial structure, confirming that the uncorrected sensitivities reported in the main text capture the dominant structural dependence of landscape functionality on habitat quality and permeability. Together, these demonstrations show that analytical sensitivity analysis provides a robust, interpretable, and scalable basis for connectivity assessment and spatial prioritization.

#### Appendix E. Landmarks

The sensitivity computations are implemented in the `ConScape` library (Van Moorter et al., 2023a). `ConScape` is implemented in the high-performance open-source Julia language, the combination of Julia's 'just-in-time' compiler, efficient algorithms, and 'landmarks' to reduce the computational load allow `ConScape` to compute landscape ecological metrics – originally developed in metapopulation ecology, such as 'metapopulation capacity' (Hanski and Ovaskainen, 2000) and 'probability of connectivity' (Saura and Pascual-Hortal, 2007) – for large landscapes. For a landscape with $n$ cells, the computation of the landscape matrix $\mathbf{M}$ requires $n \times n = n^2$ pairwise proximities ($\mathbf{K}$), which is demanding both in terms of computation and memory for high-resolution large-extent landscapes. Therefore, we developed landmark-based algorithms (Van Moorter et al., 2023a), where we compute the distances/proximities from $n$ nodes to $m$ landmarks (see for the $n$-to-$m$ method: pp. 111 in Kivimäki, 2018), which depending on the number of landmarks ($m$) can lead to a substantial reduction in the computational load with limited loss in accuracy (Van Moorter et al., 2023a). Here we explore this same approach to (1) investigate how the accuracy of the estimated sensitivity declines with decreasing number of landmarks $m$, and (2) assess the accuracy of the sensitivity estimated from separating all $n$ pixels over $n/m$ processes, which enables the parallelization of the sensitivity computation over multiple processes.

Figure E.4 shows that reducing the number of pixels used as target pixels can substantially speed-up the computation with, depending on the application, an acceptable loss in accuracy. Moreover, the landmark-based computation can also be used to split the analysis into multiple processes (results not shown here). Such splitting of the analysis can be used to parallelize the analysis over multiple processes, which can significantly reduce computation time. Furthermore, such splitting can also be used for looping over sets of landmarks to increase performance in terms of memory footprint and sometimes even in terms of processing time; however most often processing speed will decrease due to the associated 'overhead'.

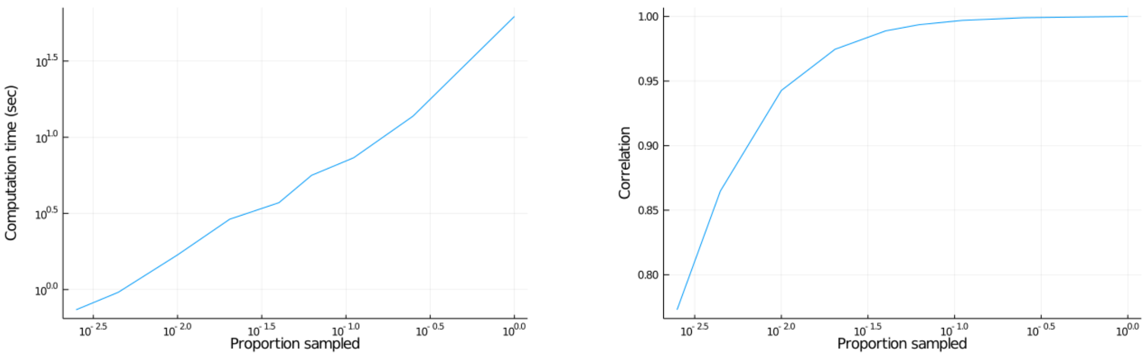


(a) (b)

Figure E.4: Computation time and accuracy of the landmark-based approach. Panel a shows the increase in computation time as the proportion of landmarks increases, and panel b shows the associated increase in accuracy (i.e. correlation between the landmark-based and full sensitivity).
